# Germline-targeting chimpanzee SIV Envelopes induce V2-apex broadly neutralizing-like B cell precursors in a rhesus macaque infection model

**DOI:** 10.1101/2023.09.21.558743

**Authors:** Rami Musharrafieh, Yana Safonova, Ge Song, Ryan S. Roark, Fang-Hua Lee, Shiyu Zhang, Jonathan Hurtado, Peter Yong, Shuyi Wang, Ronnie M. Russell, Wenge Ding, Yingying Li, Juliette Rando, Alexander I. Murphy, Emily Lindemuth, Chengyan Zhao, Andrew Jesse Connell, Wen-Hsin Lee, Nitesh Mishra, Gabriel Avillion, Wanting He, Sean Callaghan, Katharina Dueker, Anh L. Vo, Xuduo Li, Tazio Capozzola, Collin Joyce, Fangzhu Zhao, Fabio Anzanello, Weimin Liu, Frederic Bibollet-Ruche, Alejandra Ramos, Hui Li, Mark G. Lewis, Gabriel Ozorowski, Elise Landais, Brian T. Foley, Kshitij Wagh, Devin Sok, Bryan Briney, Andrew B. Ward, Beatrice H. Hahn, Dennis R. Burton, George M. Shaw, Raiees Andrabi

**Author notes:** Corresponding authors. (Y.S.); (G.M.S.); (R.A.). These authors contributed equally to this work (Rami Musharrafieh, Yana Safonova, Ge Song, Ryan S. Roark).

## Abstract

Eliciting broadly neutralizing antibodies-(bnAbs) remains a major goal of HIV-1 vaccine research. Previously, we showed that a soluble chimpanzee SIV Envelope-(Env) trimer, MT145K, bound several human V2-apex bnAb-precursors and stimulated an appropriate response in V2-apex bnAb precursor-expressing knock-in mice. Here, we tested the immunogenicity of three MT145 variants (MT145, MT145K, MT145K.dV5) expressed as chimeric simian-chimpanzee-immunodeficiency-viruses-(SCIVs) in rhesus macaques-(RMs). All three viruses established productive infections with high setpoint vRNA titers. RMs infected with the germline-targeting SCIV_MT145K and SCIV_MT145K.dV5 exhibited larger and more clonally expanded B cell lineages featuring long anionic heavy chain complementary-determining-regions-(HCDR3s) compared with wildtype SCIV_MT145. Moreover, antigen-specific B cell analysis revealed enrichment for long-CDHR3-bearing antibodies in SCIV_MT145K.dV5 infected animals with paratope features resembling prototypic V2-apex bnAbs and their precursors. Although none of the animals developed bnAbs, these results show that germline-targeting SCIVs can activate and preferentially expand B cells expressing V2-apex bnAb-like precursors, the first step in bnAb elicitation.

**Graphical Abstract:** 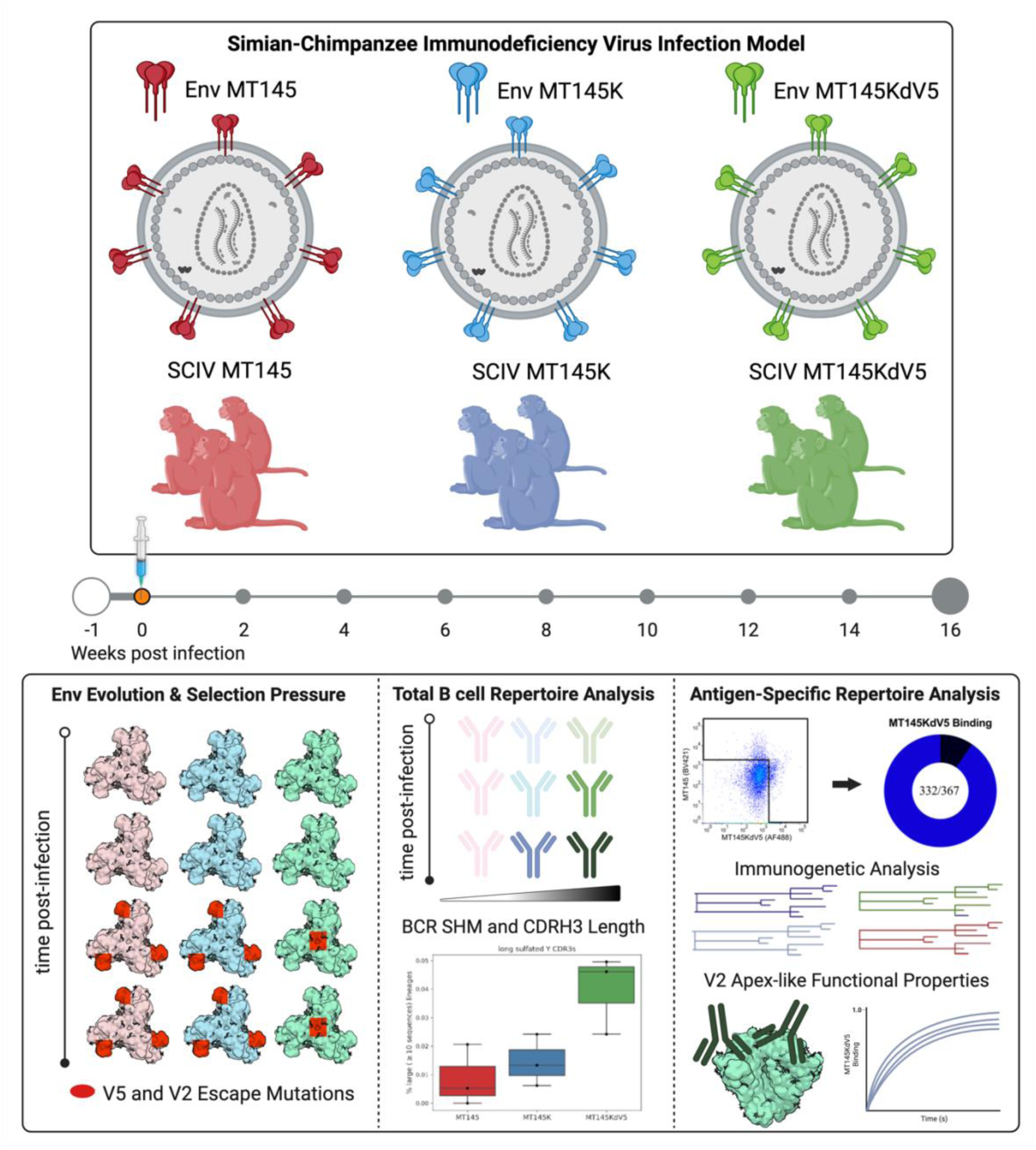

## Introduction

A major goal of HIV/AIDS vaccine research is to elicit broadly neutralizing antibodies (bnAbs), which represent a key immune defense as demonstrated by passive protection trials^1,2^. These bnAbs are grouped into six major classes based on their epitope specificities, which target V2-apex, V3-glycan, CD4 binding site (CD4bs), gp120/gp41 interface or fusion peptide (FP), the membrane proximal external region (MPER)^3,4^ and the silent face of gp120^5^. Although HIV-1 infected humans have the capacity to develop bnAbs, they do so only rarely^6–11^. This is because bnAbs are encoded by rare B cell precursors and require complex affinity maturation pathways, which makes their elicitation by vaccination particularly difficult^12–14^. In addition, each bnAb class presents unique challenges. For example, VRC01-class bnAb B cell precursors are present at relatively high frequencies (1 in ∼400,000 B cells) in humans, and germline-targeting approaches have been successful in activating them^15–19^. However, affinity maturing these responses requires extreme V-gene somatic hypermutation (SHM), which is difficult to achieve through vaccination. V2-apex and V3-glycan bnAbs exhibit comparably lower levels of SHM, but utilize long heavy-chain complementarity determining region 3 (HCDR3) loops that originate from rare VDJ recombination events^13,20–23^. BnAbs targeting the MPER region are extremely broad, but generally lack potency, require lipid binding, and are often polyreactive^24,25^.

To overcome these hurdles, HIV vaccine development efforts seek to engineer priming immunogens that stimulate the appropriate precursor B cells which are then followed by boosting immunogens that affinity mature these lineages along desired pathways. Germline-targeting immunogens have been developed for multiple bnAb specificities, including CD4bs, V3-glycan, and V2-apex, using molecular information from inferred bnAb precursors in reverse vaccine engineering approaches^17,18,21,26–30^. These immunogens have shown promise in preclinical animal models as well as humans since they generated the intended epitope-specific B cell responses^15,26,31–36^. Immunofocusing strategies, including glycan masking^37^ and protein resurface engineering^16,38–42^, have also been employed to direct B cell responses towards desired bnAb epitopes and away from off-target sites. Among various immunogen templates, native-like Env trimers that mimic the conformation of the pre-fusion spike on the virion surface^42–45^ have been shown to elicit autologous nAb responses^46–48^.

One promising target for vaccine strategies is the trimer apex^1,29,30,38,47–51^, which forms a quaternary epitope comprised of the lysine-rich V1V2 loops and surrounding glycans of three gp120 protomers^52–54^. In humans, five major V2-apex bnAb lineages have been identified: PG9, PGT145, CH01, CAP256, and PCT64^13,49,55–58^. These bnAb lineages share common immunogenetic features and utilize long anionic HCDR3 regions that allow them to penetrate the glycan shield and reach the protein surface underneath^50,52,54,57,59^. The most potent human V2-apex bnAbs utilize the IGHD3-3 gene, which encodes a germline “YYD” motif that makes contacts with basic residues within the Env C-strand^30^. These HCDR3s also contain sulfated tyrosines that are critical for epitope recognition in several V2-apex bnAb lineages^60,61^. Importantly, rhesus macaques develop V2-apex bnAbs that share very similar features, including anionic residues and sulfated tyrosines encoded by a long IGHD3-09 germline D-gene following infection with chimeric simian-human immunodeficiency viruses (SHIVs) that express the HIV-1 Env ectodomain^53,59^. Thus, SHIV infection of RMs recapitulates V2-apex bnAb development in an animal model that closely approximates HIV-1-infected humans ^53^.

Simian immunodeficiency viruses infecting chimpanzees (SIVcpz) are the zoonotic ancestors of HIV-1^62–64^ and share a last common ancestor at least 100 years ago^65^. HIV-1 and SIVcpz are thus phylogenetically closely related yet substantially more divergent than are different HIV-1 clades from each other. As a consequence, SIVcpz shares little antigenic cross-reactivity with HIV-1 in the canonical HIV-1 bnAb epitopes, except for the MPER and V2-apex epitopes, which are highly cross-reactive between SVcpz and HIV-1^66^. These findings suggest that SIVcpz Envs, if used as immunogens, might serve to immunofocus B cell responses to these highly conserved neutralizing epitopes. We previously generated a germline-targeting version of an SIVcpz Env, termed MT145, by introducing a Q171K mutation in the lysine-rich motif of strand C of the V2 loop^50^. This modified MT145K Env bound the inferred precursors of the human V2-apex bnAbs, PG9, CH01 and CAP256, and induced V2-apex directed neutralizing responses when used to immunize CH01 germline heavy chain knock-in mice^50^. To examine the germline-targeting properties of this Env in an outbred animal model and to test its bnAb-induction potential in the context of infection where viral Envs and antibody responses can coevolve, we generated simian-chimpanzee immunodeficiency viruses (SCIVs) that expressed wildtype MT145 and MT145K Envs, as well as a minimally modified derivative, MT145K.dV5, which has a shortened variable loop 5 (V5). We used these chimeric viruses to infect three groups of rhesus macaques. While all SCIVs established productive infections characterized by high peak and setpoint viral loads, animals infected with the germline-targeting SCIV_MT145K and SCIV_MT145K.dV5 exhibited a larger number of clonally expanded B cell lineages as well as more lineages that featured long (>24 residue) anionic HCDR3s than animals infected with wildtype SCIV_MT145. SCIV.MT145K.dV5 infected animals also exhibited sequence changes in the V2-apex bnAb core epitope suggestive of antibody mediated pressures as well as an enrichment of long HCDR3s and IGHD3-09 gene usage in antigen-specific memory B cells. These results suggest that germline-targeting SCIVs were able to activate and preferentially expand V2-apex bnAb-like precursors within the first 4 to 8 weeks post infection.

## Results

### SCIVs expressing wildtype and germline targeting MT145 Envs establish persistent infections and induce autologous neutralizing responses in rhesus macaques

SHIV infections provide insight into the immunogenicity of HIV-1 Env glycoproteins since these viruses replicate continuously over the course of the infection and express native trimers that co-evolve with germline and intermediate B cell receptors^53,67–69^. To investigate whether the germline-targeting MT145K Env can activate desirable B cell precursors in an outbred animal model, we generated SCIVs expressing both wildtype MT145 and MT145K Envs as well as a minimally modified MT145K.dV5 Env, which partially lacked immunodominant off-target epitopes (see below). These SCIVs were used to infect three groups of three rhesus macaques (Fig. 1A) after transient CD8+ T cell depletion^70^, which were then followed longitudinally for up to 88 weeks to assess in vivo viral replication, sequence evolution and nAb responses. Both SCIV_MT145 and SCIV_MT145K replicated efficiently in all RMs with high peak and setpoint viral loads, while SCIV_MT145K.dV5 had slightly lower setpoint viral loads (Fig. 1B). Two SCIV_MT145 infected RMs (6931, 6933) and one SCIV_MT145K infected RM (T924) with particularly high setpoint viral loads developed an AIDS-like illness and were euthanized at weeks 29, 18, and 22, respectively. The animal that died most rapidly (6933) failed to develop anti-SCIV antibodies, a feature previously described for SIVsm and SIVmac infected “rapid progressor” animals^71^.

**Figure 1.**
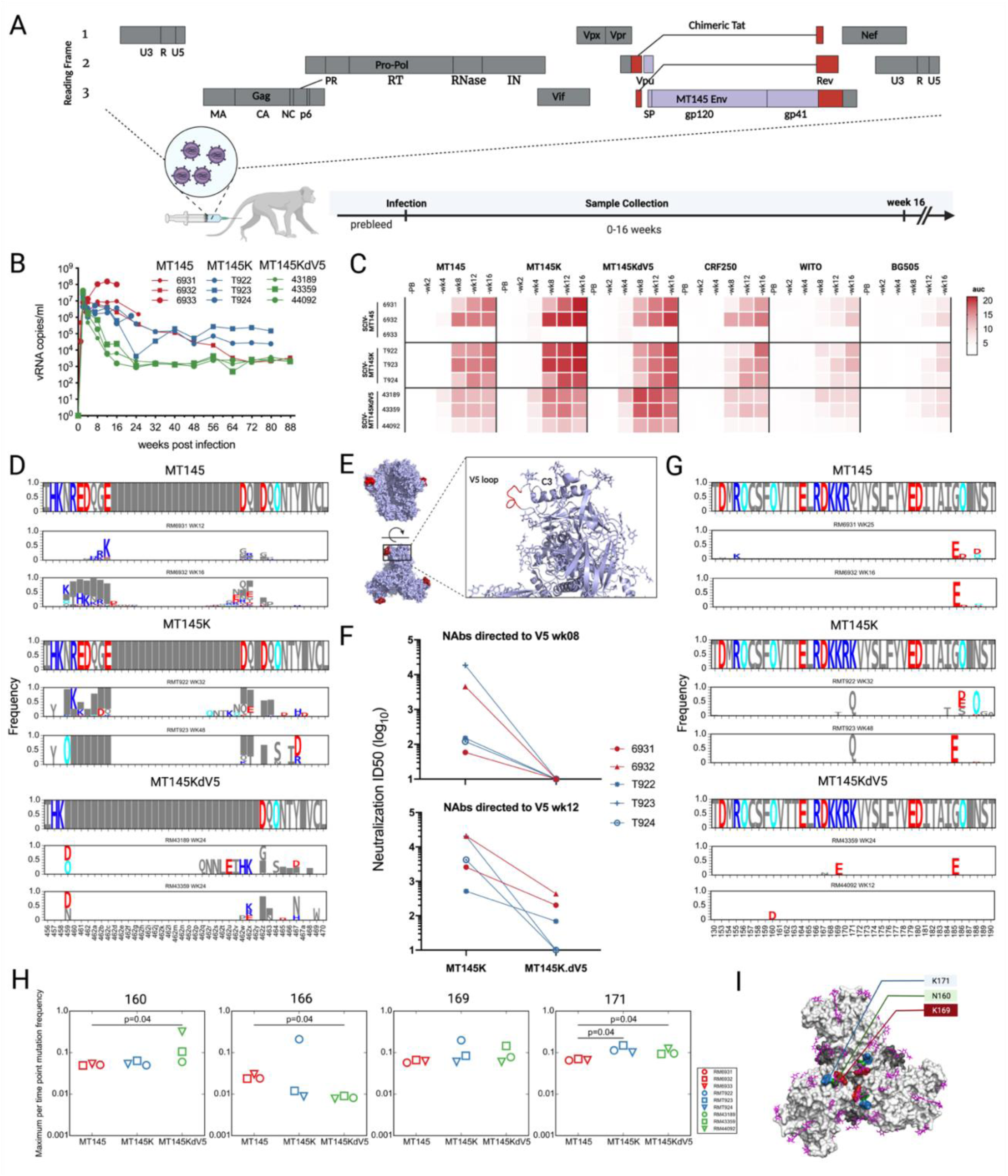
Resurfacing of SCIV expressed MT145 Envs redirects immune responses. (A) Generation of a SCIV construct that expresses SIVcpz Env ectodomains. The SIVcpz MT145 *vpu-env* region (blue) was cloned into an optimized SHIV vector (LI, JVI) consisting of a SIVmac766 proviral backbone (grey) and HIV-1 derived *tat* and *rev* genes (red). SCIVs expressing wildtype MT145 (n=3) as well as germline-targeting MT145K (n=3) and MT145K.dV5 (n=3) Envs were used to infect 9 naïve Indian rhesus macaques. (B) Plasma vRNA kinetics in group of RMs infected with SCIV.MT145 (red), SCIV.MT145K (blue) and SCIV_MT145K.dV5 (green). Blood samples were initially collected at -1 (pre-bleed) up to 96 wpi. (C) Serum ELISA binding titers in SCIV-infected animals against MT145K, MT145K.dV5, CRF250, WITO, and BG505 Envs during the first 16 weeks of infection. A heatmap indicates area under the curve (auc) values. (D) Single genome sequencing (SGS) alignment of the *env* V5 region gene sequences for 2/3 RMs in each group. Sequence logos show the per time-point frequency of mutations away from the TF sequence in the sequences from each RM. Red indicates negatively charged amino acids, blue for positively charge amino acids and cyan for Asn in potential N-linked glycans. Grey box indicates a gap. (E) Structure of Mt145K Env with the V5 loop shown in red (PDB: 6OHY). (F) Plasma neutralization from SCIV_MT145 and SCIV_MT145K infected animals at 8 wpi (left) and 12 wpi (right) against MT145K and MT145K.dV5 Env containing pseudoviruses. (G) SGS alignment of the *env* V2 region gene sequences for 2/3 RMs in each group. Same formatting as (D). (H) Maximum per-time point mutation frequency for each RM for residues 160, 166, 169, and 171 from NGS sequencing analysis. Statistics measured using Wilcoxon rank sum test, two-sided p value shown if p < 0.05 and are not corrected for multiple testing. (I) MT145K trimer (PDB: 6OHY) with mutations identified in the V2 region and C-strand colored green, red, and blue for N160, K169, and K171, respectively. gp120, gp41, and surface glycans shown in light grey, dark grey, and purple, respectively.

Analysis of plasma samples by enzyme linked immunosorbent assay (ELISA) showed that all SCIV infected animals, except for the rapid progressor animal 6933, developed antibodies that bound to both autologous and heterologous soluble Env trimers as early as 4-8 weeks post infection (Fig. 1C). All animals except for RM 6933 also developed high titer autologous neutralizing antibodies as well as antibodies that potently neutralized tier 1 HIV-1 strains. However, none of the animals developed significant heterologous neutralization breadth over the course of their infection (Fig. S1). Although plasma samples from four animals neutralized a handful of heterologous HIV-1 strains, all these responses were only weakly cross-neutralizing, with reciprocal 50% inhibitory dilutions (ID_50_) less than 1:100 with no increase in breadth and potency over time. Thus, none of the SCIV infected RMs developed bnAbs.

To characterize sequence changes from the infecting virus strains, we used limiting dilution PCR of plasma viral RNA/cDNA to generate longitudinal *env* gene sequences for each infected RM. This method, referred to as single genome sequencing or SGS ^72,73^, retains genetic linkage across the complete gp160 gene and eliminates PCR induced mutational artifacts from finished sequences. An alignment of these sequences revealed strong selection in variable loop 5 (V5) in both SCIV_MT145 and SCIV_MT145K infected RMs, as evidenced by a diverse set of mutations, including amino acid substitutions, insertions, and deletions, which led to changes in length, net charge, and number of glycans (Fig. 1D-F, Fig. S2). Among these mutations, the same nine amino acid V5 loop deletion was observed in multiple animals, suggesting a common pathway of escape from potent autologous neutralizing antibodies (Fig. S2). Longitudinal Antigenic Sequences and Sites from Intra-Host Evolution (LASSIE) analysis, which reveals sites under selection pressure using >50% transmitter/founder-loss at any time point, confirmed this selection in the V5 region as well as additional sites of selection elsewhere in gp160 (Figs. S3 and S4). These data indicated the presence of an immunodominant strain-specific neutralizing epitope in the V5 loop of the MT145 Env that elicited off-target responses. When we examined the primary, x-ray crystallographic and cryoEM structures of MT145 Env, we noted that the V5 loop was atypically long and disordered, likely contributing to is immunodominance. To remove this immunodominant epitope, we deleted the nine V5 residues that comprised the naturally-occurring truncation, thus generating SCIV_MT145K.dV5.

To confirm that SCIV_MT145K.dV5 retained its germline targeting potential, we tested its neutralization sensitivity to inferred germline (iGL) or reverted unmutated ancestors (RUAs) of the V2 apex bnAbs CH01^55^, PG9^21,74^, and PG16^74^, the inferred germline (iGL) of PCT64^56^ as well as the unmutated common ancestors (UCA) of VRC26^13^ and RHA1^53^. As expected, wildtype SCIV_MT145 was resistant to neutralization by all precursor antibodies (Fig. S5). However, both SCIV_MT145K and SCIV_MT145K.dV5 were neutralized by CH01_RUA and VRC26_UCA at IC_50_ values of 4.6-8.1 µg/ml and 124.3-202.7 µg/ml, respectively. While this analysis likely underestimated germline interactions, the results confirmed that the modified SCIVs engaged at least two human V2-apex precursors.

We next used both SGA of longitudinal Env sequences as well as next generation sequencing (NGS) of the V1V2 region to investigate within-host Env evolution across all three groups of SCIV-infected RMs. These analyses showed residual selection on the V5 region in MT145K.dV5 animals, despite epitope trimming, suggesting that resurfacing did not completely eliminate responses to this immunodominant off-target site (Fig. 1D, Fig. S2-4). This was confirmed by negative stain polyclonal electron microscopy mapping (EMPEM), which showed V5 binding of serum IgG Fabs for all animals except the rapid progressor RM 6933 (Fig. 2). LASSIE analysis also showed selection pressures in Env regions other than V5 for MT145K.dV5 as well as MT145 and MT145K circulating viruses (Fig. S4). Examining the V1V2 region in particular for mutations that altered the TF amino acid sequence, we found that both SCIV_MT145K and SCIV_MT145K.dV5, but not SCIV_MT145 infected RMs exhibited amino acid substitutions at positions 160, 166, 169 and 171 at one or more time points (Fig. 1G & 1H, Fig. S6 & S7); of these, only the site 171 was chosen for all three MT145K infected RMs, using the pre-specified LASSIE selection criterion of 50% or more mutation at any time point (Fig. S4). Statistical analyses of the corresponding mutation frequencies in the NGS dataset showed significantly higher mutation frequency for either SCIV_MT145K or SCIV_MT145K.dV5 groups as compared to WT SCIV_MT145 group at positions 160 and 171 (Fig. 1H, S7), while SCIV_MT145 group showed significantly higher frequency at 166 than SCIV_MT145K.dV5 group. However, despite selection in V1V2 and gp41 regions, no changes were observed at the canonical V2-apex bnAb site for any animal infected with SCIV_MT145K and SCIV_MT145K.dV5 from EMPEM analysis (Fig. 1I & 2). Together, these results suggest early B cell priming at the V2-apex bnAb epitope by germline-targeting MT145 Envs, but these responses did not contribute to the dominant serum antibody specificities.

**Figure 2.**
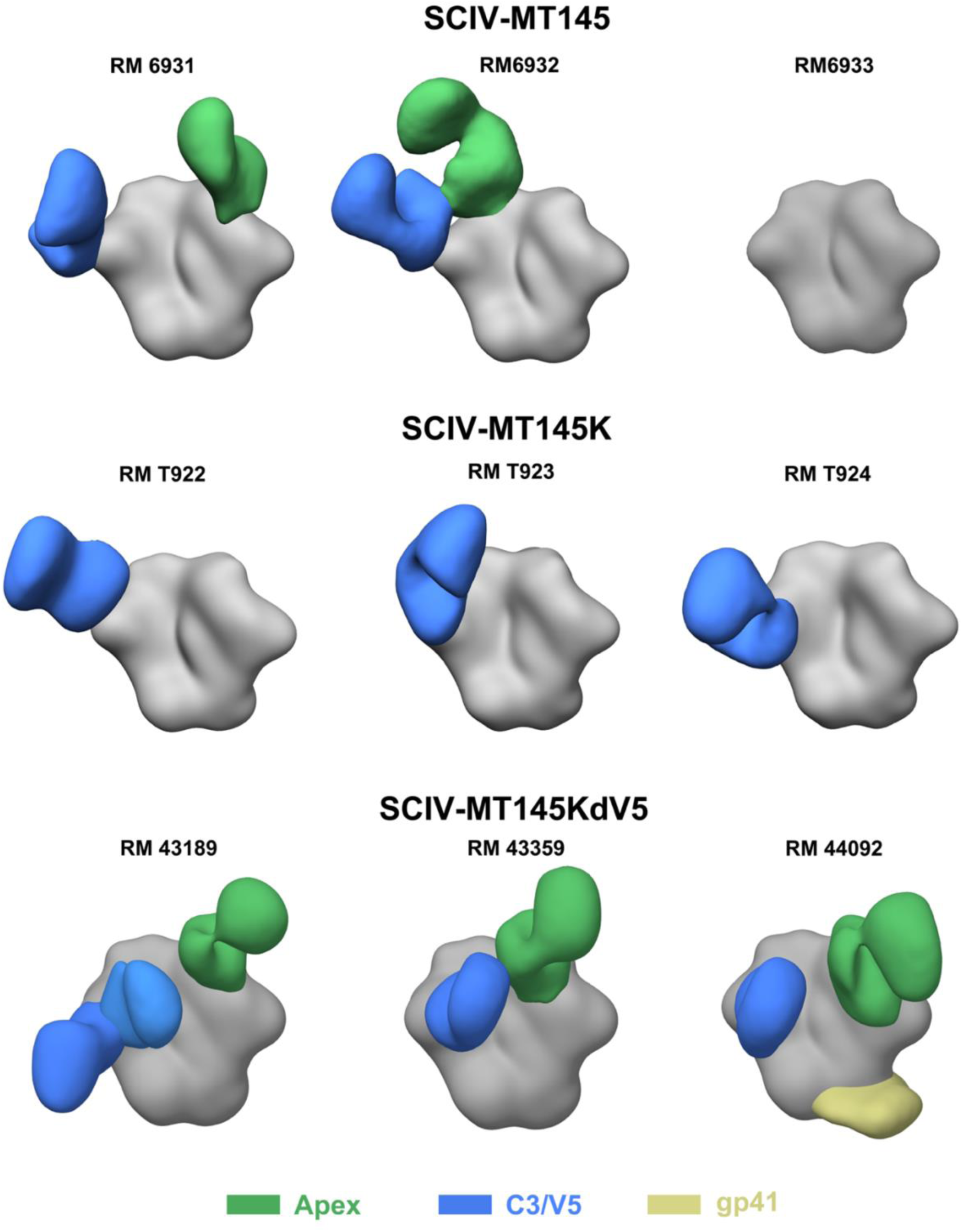
MT145, MT145K, and MT145K.dV5 trimers bind to digested Fabs from SCIV-infected animal sera. 3D reconstructions generated from negative stain electron microscopy epitope mapping (EMPEM) of serum Fabs from SCIV-infected rhesus macaques at week 12 time point. The Env corresponding to the SCIV Env was used for complexing. Fabs colored based on epitope, with V1/V2 (apex) in green, C3/V5 in blue, and gp41 in yellow. Top row shows SCIV_MT145 Envs bound to Fabs at the C3/V5 region (blue) and apex adjacent sites (green). Center row shows SCIV_MT145K Envs bound primarily to C3/V5 Fabs (blue). Bottom row shows SCIV_MT145K.dV5 Envs bound to apex adjacent (green), C3/V5 (blue), and gp41 (yellow) sites. Representative maps have been deposited to the Electron Microscopy Data Bank (see STAR Methods).

### Infection with SCIV_MT145K.dV5 results in early expansion of isotype-switched B cell lineages with long anionic HCDR3s

To investigate the immunogenetic features that define the earliest immune responses for each SCIV-infected RM group, we performed next-generation sequencing (NGS) of longitudinal peripheral blood B cells (up to week 16) to examine the bulk IgG and IgM repertoires (Fig. 3A). Prior to infection, animals from all three groups (SCIV_MT145, SCIV_MT145K and SCIV_MT145K.dV5) showed similar numbers of long (≥ 24 aa) HCDR3-bearing B cells in their IgG repertoires (Fig. 3B). Differences between the groups appeared as early as 2 wpi, with two animals in the SCIV_MT145K.dV5 group showing an increased number of long HCDR3 lineages, which peaked at 4 wpi (P=0.04, the Kruskal-Wallis test). All three animals in the SCIV_MT145K.dV5 group were substantially enriched in the number and fraction of lineages with long HCDR3s (P-values are 0.01 and 0.005, the linear mixed model) by week 4, which was maintained until 16 wpi (Fig. 3B-C). In contrast, no significant enrichment for long HCDR3 B cell lineages were observed in animals infected with wildtype SCIV_MT145. Long HCDR3s detected in SCIV_MT145K.dV5 infected animals at weeks 4 to 12 were also more anionic compared with SCIV_MT145 and SCIV_MT145K infected animals (P=0.008, the linear mixed model), containing DD, DE, ED, or EE residues as well as predicted sulfotyrosines (Fig. 3D). Overall, SCIV_MT145K.dV5 infected RMs had the highest percentage of clonally expanded long HCDR3s enriched for sulfated tyrosines across all time points, whereas SCIV_MT145 infected RMs had the lowest (Fig. 3E).

**Figure 3.**
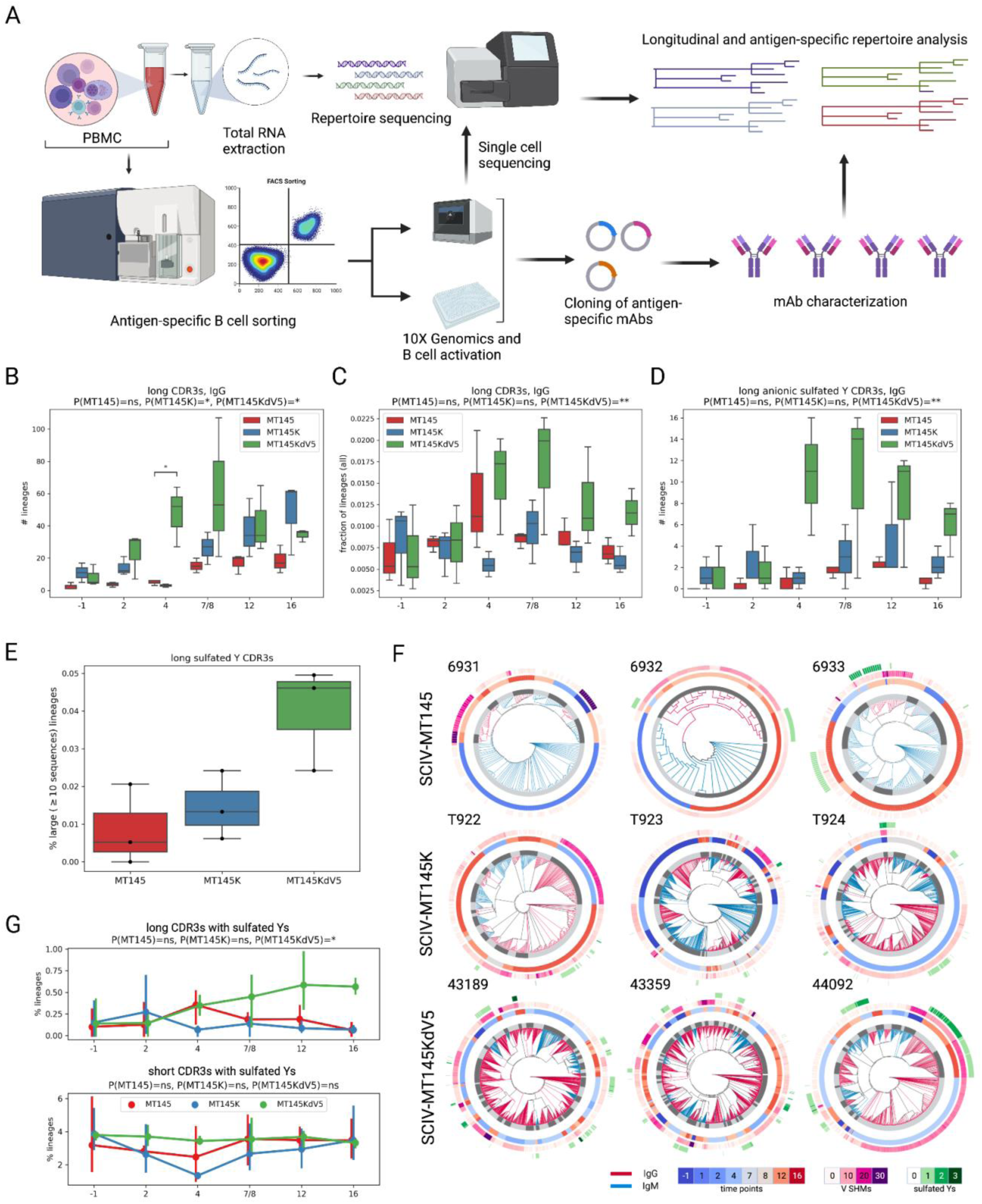
SCIV_MT145K.dV5 infection results in early expansion of isotype-switched lineages with long, anionic HCDR3s. (A) Experimental pipeline to determine repertoire and antigen-specific B cell responses as well as antibody characterization in rhesus macaques infected with SCIVs. (B) Number of lineages with long (≥24 aa) HCDR3s in isotype-switched IgG sequences at all time points for all 9 rhesus macaques. (C) Fraction of the total number of lineages with long HCDR3s in isotype-switched IgG sequences. (D) Total number of lineages with long HCDR3s that feature anionic, sulfated tyrosines in IgG sequences. Sulfated Ys were predicted using the Sulfinator prediction tool with a default E-value set to 30. (E) The percent of total lineages with HCDR3s that are long with predicted Y sulfations by group for all time points combined together. (F) Circular phylogenetic trees featuring all clonally expanded (defined as ≥10 sequences) lineages with long HCDR3 for all nine animals. Inner trees are colored red for IgG sequences and blue for IgM. The first inner circle represents unique lineage clusters. The second inner circle defines the time point in which the sequence was identified. The last two circles represent the V-region somatic hypermutation and sulfated Ys shown in gradients of purple and green, respectively. (G) Percent of lineages that feature HCDR3s with predicted Y sulfations for long (top) and short (bottom) HCDR3s at each time point.

SCIV_MT145K and SCIV_MT145K.dV5-infected animals featured significantly more clonally expanded (>10 sequences) long HCDR3 lineages compared with SCIV_MT145 infected RMs (P=0.02, the Kruskal-Wallis test). The three SCIV_MT145K-infected animals had 20, 32, and 46 unique expanded lineages, while the SCIV_MT145K.dV5 featured slightly higher numbers of 24, 50, and 59 lineages, respectively (Fig. 3F).

To illustrate differences in IgM and IgG repertoires during the first 16 weeks of SCIV infection, we constructed phylogenetic trees of all clonally expanded (>10 sequences) long HCDR3 lineages identified in longitudinal PBMC samples (Fig. 3F). Comparing the three groups, all animals had at least 3 clonally expanded lineages containing long HCDR3s. SCIV_MT145 infected animals had between 3 to 14 expanded lineages. For 50–90% of these lineages across three animals, IgM was a dominant isotype suggesting a memory B cell origin (Fig. 3F, upper row). Sulfotyrosines were predicted for 2 of the 3 animals in the SCIV_MT145 group (present in four clones) and were predominantly present in IgG lineages.

For animals in the SCIV_MT145K group, 76–95% of expanded long HCDR3 lineages were isotype switched relative to SCIV_MT145 animals (Fig. 3F, middle row). Most of the expanded IgG lineages in the SCIV_MT145K group were identified 4-12 wpi. Predicted sulfotyrosines were observed in all three animals in SCIV_MT145K group and were predominantly identified in IgG lineages.

SCIV_MT145K.dV5 animals had the 75–90% of isotype-switched IgG lineages with expanded long HCDR3s (Fig 3B, Fig S8). For animals in this group, most of the IgM lineages were identified prior to infection or less than 4 wpi, suggesting that most of the observed expansion was antigen-driven. Predicted sulfotyrosines were notably higher compared to the SCIV_MT145 and SCIV_MT145K groups and were identified in 27–32% of lineages across animals in the SCIV_MT145K.dV5 group. overall, clonally expanded lineages in MT145KdV5 animals have higher fractions of IgG sequences compared to MT145K and MT145 groups (P=0.0002, the Kruskal-Wallus test)

### SCIV_MT145K.dV5 infection elicited larger and more clonally expanded long HCDR3 lineages encoding germline D-genes with anionic residues and sulfated tyrosines

Unlike in animals infected with SCIV_MT145 and SCIV_MT145K, the percentage of lineages with long HCDR3s and sulfated tyrosines increased in SCIV_MT145K.dV5-infected animals over time (P=0.03, the linear mixed model) (Fig 3G, Fig. S8). Importantly, this effect was not observed for lineages with short HCDR3s (<24 aa) and sulfated tyrosines (Fig. 3G). While the fraction of large, expanded lineages remained stable in SCIV_MT145K and SCIV_MT145K.dV5 infected animals, this fraction decreased slightly in SCIV_MT145 infected animals (Fig. S9A). All groups showed increases in the average HCDR3 length over time, with the SCIV_MT145K.dV5 group showing the most pronounced increase (Fig. S9B).

We also observed differences in IGHD gene usage in expanded lineages with long HCDR3s among the three groups (Fig. S10 and S11). While SCIV_MT145-infected RMs used the IGHD6-25 gene more frequently than SCIV_ MT145K and SCIV_MT145K.dV5-infected animals (Fig. S9 and S10), SCIV_MT145K.dV5-infected animals exhibited the greatest enrichment of the IGHD3-09 gene, especially at later time points. The IGHD3-09 gene contains an “EDDY” motif that is highly conserved in V2-apex bnAbs isolated from macaques^53^ and likely contains germline-encoded features that are essential for activating the respective precursors. Moreover, these long HCDR3 germline D gene features are shared between human and macaque V2-apex bnAbs (Fig S13). In contrast, there was no statistically significant enrichment for D genes in short HCDR3 lineages (Fig. S11 and S12), although SCIV_MT145 infected animals used the IGHD3-9 gene less frequently (Fig. S11D).

We also analyzed V gene frequencies in IgM and IgG repertoires for all three groups. No enrichment of unique IGHV gene families or alleles were observed in any group, with most animals utilizing the IGHV3 and IGHV4 families (Fig. S14), consistent with gene usage in rhesus and cynomolgus macaques^75^.

Expanded long HCDR3 lineages for all animals contained moderately to highly mutated members with varying numbers of predicted sulfation sites. The most abundant lineages in SCIV.MT145-infected RMs were significantly smaller and less diverse than those found in SCIV_MT145K and SCIV_MT145K.dV5 infected animals (Fig. S16-18). Only three of five of the largest long HCDR3 lineages contained an ‘EDDY’ motif, and only one ‘EDDY’ lineage was IgG isotype-switched and contained significant SHM and predicted sulfotyrosine residues (Fig. S15). In contrast, the largest lineages in the SCIV_MT145K group were much more diverse and expanded. Eleven of 12 large long HCDR3 lineages from the SCIV_MT145K group contained ‘EDDY’, or ‘YY’, and six of 11 of these lineages were predominantly IgG (Fig. S16). For the SCIV_MT145K.dV5 group, all of 15 of the largest long HCDR3 lineages contained ‘EDDY’, ‘YY’, or ‘DDY’ motifs, and only two of these lineages were predominantly IgM (Fig. S17). Collectively these results suggest that the germline-targeting SCIV that was modified to limit V5 directed off target responses generated an antibody response that most closely resembled the activation of V2-directed bnAb precursors^50^.

### SCIV infection reshapes antigen-specific B cell repertoires toward V2-apex bnAb-like features

Given that IgM and IgG repertoires were significantly different between the three groups of SCIV infected RMs, we next asked whether these differences were also present in the antigen-specific repertoires. We sorted antigen-specific memory B cells from peripheral blood collected 8 and 12 wpi, since this was when the most substantial differences in the overall IgH repertoires were observed (Fig. 4A). For each animal, memory B cells were sorted using MT145, MT145K and MT145K.dV5 trimer probes to enrich BCRs with affinities for one or more of these antigens (Fig. 4B). The percentages of total antigen-specific IgG-positive B cells were similar across groups (Fig. 4B), with each group containing BCRs specific for all three trimers. There was a trend for higher numbers of antigen-specific B cell lineages as well as lineages with long HCDR3s in SCIV_MT145K and SCIV_MT145K.dV5-infected animals compared to SCIV_MT145-infected animals, but this did not reach statistical significance (Fig. 4C). HCDR3 lengths followed a normal distribution with a right shoulder towards longer HCDR3s and the peak fraction of lineages for both antigen-specific and repertoire sequences at 14 amino acids (Fig. 4D). Mutation levels in the V_H_ region were slightly higher for MT145K.dV5 antigen-specific sequences (3.96%) compared with MT145K (2.6%) and MT145 (2.1%) antigen-specific sequences. Similar to the IgM and IgG total repertoire frequencies, IGHV3 and IGHV4 families were mostly observed in longer HCDR3 B cells for all three groups.

**Figure 4.**
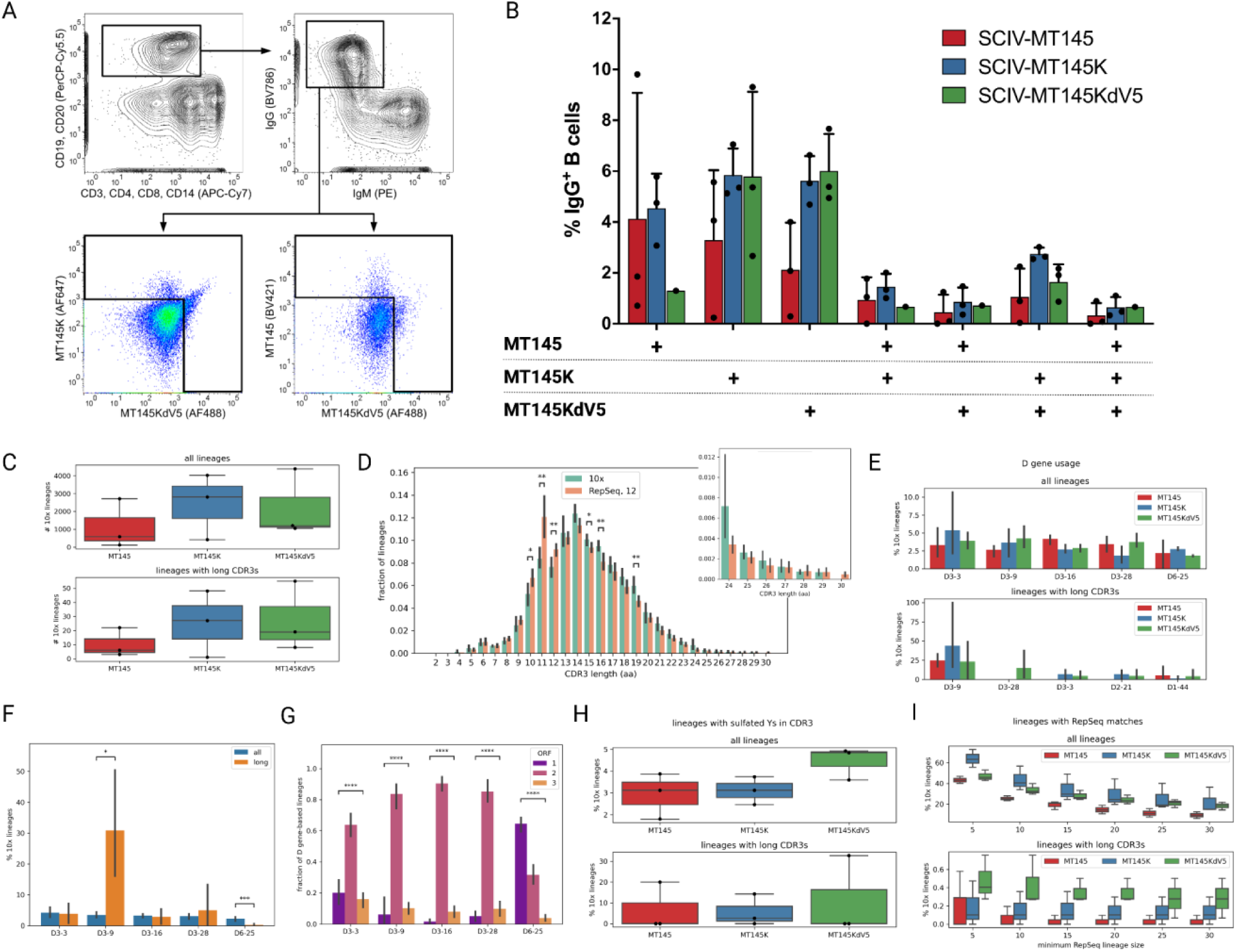
Antigen-specific IgG B cell lineages match features found in repertoire analysis. (A) MT145, MT145K, and MT145K.dV5 probes were used to sort all antigen-specific IgG (+) IgM (-) B cells from week 12 samples for single cell sequencing. (B) Sorting analysis showing the percent of total B cells from each group that is specific for one or more Env from each group. (C) The total number of lineages identified from single cell analysis per group (top) or the total number of lineages with long HCDR3s (bottom). (D) Distribution of HCDR3 lengths in the fraction of lineages from antigen-sorted B cells (green) and those from total repertoire analysis (orange). Long HCDR3s (residues 24 aa or longer) enlarged on the right panel. (E) Percent of all (top) lineages or lineages with long HCDR3s (bottom) that use certain IGHD genes by group. (F) Percent of all (blue) lineages or lineages with long HCDR3s (orange) that use certain IGHD genes from all nine rhesus macaques. (G) Reading frame used by IGHD genes in antigen-sorted B cell sequences from all animals. (H) Percent of lineages with sulfated Ys located in all (top) or long (bottom) HCDR3 sequences by group. (I) Percent of lineages for all (top) or long (bottom) HCDR3s that are also present in the overall repertoire by lineage size. P-values showing pairwise differences between percentages of lineages with long CDR3s across all time points. P-values were computed using the one-way ANOVA test. (*P < 0.05; **P < 0.01)

IGHD3 gene usage was similar among all three groups, with no significant differences between groups (Fig. 4E). The IGHD3-9 gene was overrepresented in all long HCDR3 lineages, while the IGHD6-25 gene was more abundant in short HCDR3s (Fig. 4F). The enrichment of specific IGHD3 gene families in total and antigen-specific repertoire sequences with long HCDR3s suggested that specific motifs were selected during clonal expansion and in the case of the IGHD3-09 gene, the germline-encoded “EDDY” motif was consistently observed. Since, during B cell development, nucleotide insertions and deletions in VDJ junctions could change germline-encoded motifs present in the D or J genes, we examined the open reading frame (ORF) across all common IGHD genes identified in long HCDR3 lineages. Interestingly, we observed very little variation in this ORF across the various IGHD3 genes, suggesting preservation of germline-encoded motifs important for V2 bnAb development (Fig. 4G). Analysis of the ORFs of IGHD genes showed that ORFs with ≥ 2 tyrosines had much higher usage in BCR sequences compared with ORFs with < 2 tyrosines (Fig. S19).

We next compared lineages from total IgM and IgG repertoires with antigen-specific single B cell sequences. Similar to the bulk repertoire sequencing data, the percent lineages with sulfated HCDR3 tyrosines tended to be higher for the SCIV_MT145K.dV5-infected group compared with the SCIV_MT145 or SCIV_MT145K-infected groups, although this did not reach statistical significance (Fig. 4H). All three groups showed similar numbers of matching lineages, with a slightly higher fraction in the SCIV_MT145K group among all sequences and the SCIV_MT145K.dV5 group among long HCDR3 sequences, independent of lineage size (Fig. 4I, S20A). Antigen-specific members from SCIV_MT145K and SCIV_MT145K.dV5-infected groups were mapped to large, expanded long HCDR3 lineages in IgG and IgM repertoires that utilized the IGHD3-09 gene, with more than one member identified in each lineage (Fig. S20B). Collectively these data suggested that the observed reshaping of the B cell repertoire following SCIV infection was driven at least in part by antigen-specific B cells.

### SCIV_MT145K.dV5-induced long HCDR3 antibodies with V2-apex bnAb features exhibit apex epitope binding properties

By minimizing responses to the immunodominant V5 region and introducing the germline-targeting Q171K substitution, the antigen-specific responses in SCIV_MT145K.dV5 infected animals exhibited features characteristic of V2-directed precursor activation. We thus characterized antigen-specific B cells from SCIV_MT145K.dV5-infected animals by performing a sort at week 16 and testing the recovered cells in a B cell activation method (Fig. 5A). A total of 1,271 antigen-specific B cells were isolated, resulting in the recovery of 367 IgG-secreting B cells after 14 days of culture (Fig 5B). Nearly all (332/367) secreted antibodies (mAb) bound the MT145K.dV5 SOSIP trimer, and 26.4% or 7.7% were able to neutralize MT145K.dV5 or CRF250 Env-containing pseudoviruses, respectively (Fig 5B, Supplementary Table 01). While almost all mAbs (94.5%) bound a version of the MT145K.dV5 trimer in which the N160 was removed (MT145K.dV5-N160K), a fraction (5.2%) was dependent on glycan N160. Only 19 mAbs bound the MT145K.dV5 trimer in a N160 glycan-dependent manner and were also able to neutralize CRF250 pseudoviruses. The most V gene-mutated mAbs (SHM level of 12.6%) were those that neutralized the MT145K.dV5 virus but maintained binding to the MT145K.dV5-N160K SOSIP. However, mAbs that were dependent on N160 were also moderately mutated (SHM level of 8.7%). These mAbs also had the second longest HCDR3s on average (19.5 aa).

**Figure 5.**
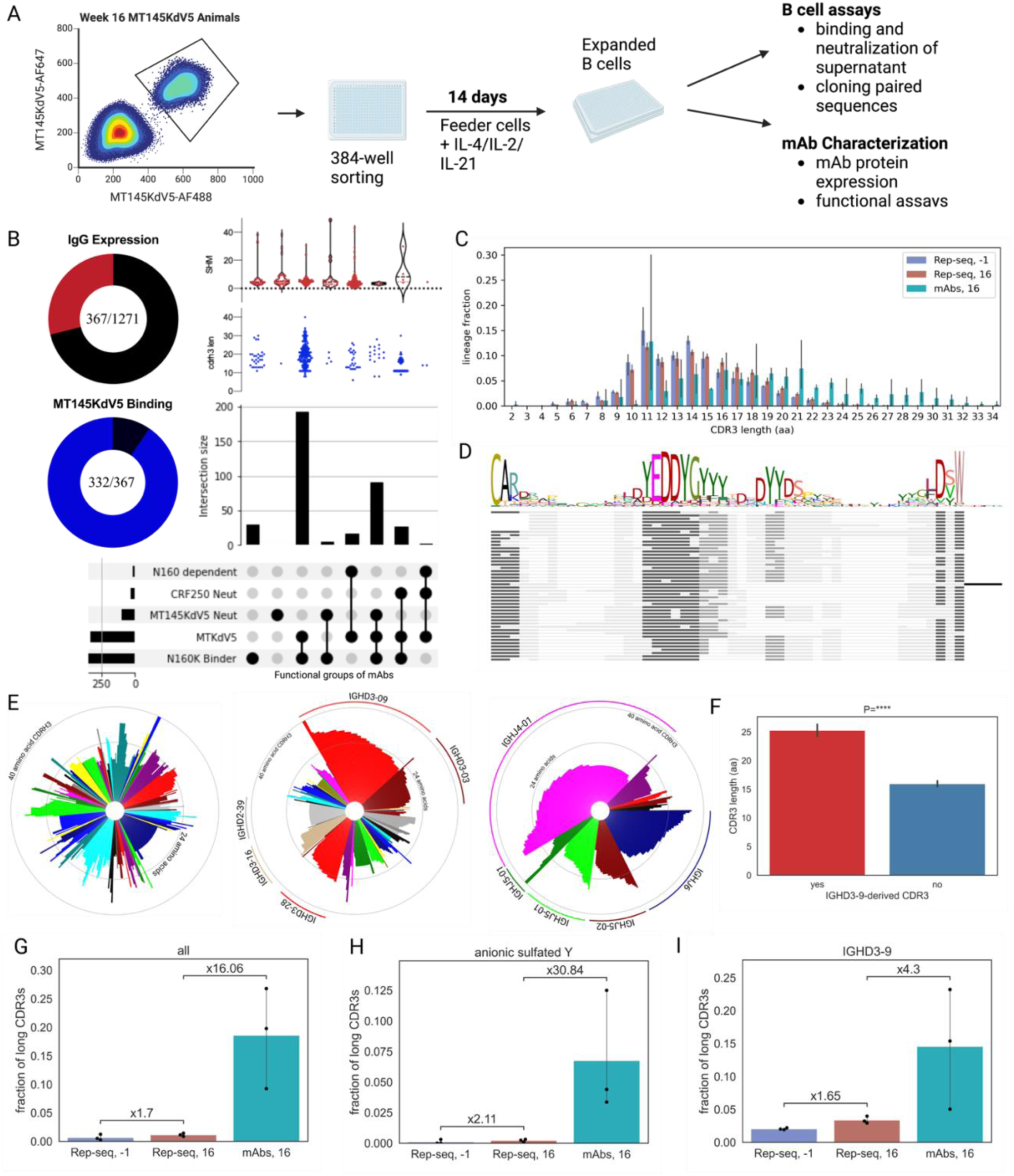
Binding mAbs from antigen-sorted B cells from SCIV_MT145K.dV5 animals are biased for long HCDR3 and IGHD3-09 gene usage. (A) Illustration of experimental design for isolating antigen-binding mAbs from B cells. Week 16 SCIV_MT145K.dV5 infected PBMC samples were sorted for antigen-specific IgG(+) B cells using MT145K.dV5 probes and cultured in 384-well plates for activation and mAb characterization. (B) Pie plots depicting total (red) and MT145K.dV5-specific (blue) mAbs isolated and characterized from cultured B cell supernatants. Upset plot showing intersecting sets of binding and neutralization properties as well as set count (histogram). HCDR3 lengths (red) and SHM levels (blue) are shown as categorical plots above each set. (C) Distribution of HCDR3 lengths in the fraction of lineages from antigen-sorted B cell cultures at week 16 (red) and those from total repertoire analysis preinfection (purple) and at week 16 (green). (D) Alignment of representative sequences from long HCDR3 mAbs that bind to MT145K.dV5 and use IGHD3-09 gene. (E) Circular bar plots representing HCDR3 lengths for IGHV, IGHD, and IGHJ gene sequences. Genes are color coded and enriched genes are individually labeled. (F) HCDR3 lengths from sequences that use or do not use IGHD3-09 gene from all three SCIV_MT145K.dV5 animals at week 16. (H) Fraction of all long HCDR3s present in the repertoire prior to and 16 wpi compared to antigen-sorted B cells at week 16. Fraction of long HCDR3s with anionic features and predicted Y sulfations in the repertoire prior to and 16 wpi compared to antigen-sorted B cells at week 16. (I) Fraction of long HCDR3s using the IGHD3-09 gene present in the repertoire prior to and 16 wpi compared to antigen-sorted B cells at week 16.

All antigen-specific mAbs isolated from B cell cultures were skewed towards longer HCDR3s (Fig. 5C), although there was one 11 aa HCDR3 expanded lineage from RM 44092 that was not observed in the pre-infection repertoire (Figs. 5C, 4D). Among all mAbs, 22% encoded the IGHD3-09 gene with the “EDDY” motif (Fig. 5E). However, for mAbs with long HCDR3s, IGHD3-09 was by far the most abundant and observed in 76% of all sequences. In contrast, IGHV or IGHJ genes were highly variable (Fig. 5F and G). Finally, while repertoire sequences in SCIV_MT145K.dV5 infected monkeys showed enrichment for long HCDR3s and anionic sulfated tyrosines, this fraction was significantly increased in the antigen-specific antibodies sorted from week 16 (Fig 5H-J).

Encouraged by the immunogenetic and functional findings of antigen-specific mAbs characterized for animals in the SCIV_MT145K.dV5 group, we next asked-what are the functional characteristics associated with mAbs expressed from antigen-specific V2-apex bnAb-like B cell precursors? Therefore, a subset of antigen-specific long HCDR3 mAbs encoding the IGHD3-09 that were isolated at week 12 and 16 from SCIV_MT145K.dV5 infected animals were cloned and expressed (Fig. 6). Although these mAbs were restricted by their IGHD gene usage, they all shared a similar anionic motif centered in the HCDR3 region (Fig. 6A). Eighteen mAbs (01, 03, 07, 15, 18, 23, 24, 27, 31, 38, 41, 51, 53, 55, 58, 59, 60, 70) exhibited significant binding to the MT145K.dV5 trimer as well as N160 glycan dependence. Out of this subset of N160 glycan dependent mAbs, nine (15,18, 23, 24, 27, 31, 41, 58, 70) were predicted to be trimer dependent based on their inability to bind MT145K.dV5 gp120 protein. These results suggest that, although we were enriching for V2 bnAb-like HCDR3 signatures, these mAbs showed surprising functional diversity. If a subset of these mAbs are true V2-apex bnAb precursors, we would expect them to show higher affinity to germline targeting MT145 trimer and lower affinity to trimer variants with mutations at specific V2 sites that interact with bnAbs. To test this, we compared the binding of these mAbs to MT145, MT145K and MT145K.dV5 SOSIP trimers as well as derivatives with mutations in the V2 epitope (Fig. 6B). The results showed less binding to MT145, MT145K-N160K and MT145K-N169E/N171E SOSIP trimers compared to MT145K or MT145K.dV5 SOSIP trimers. Since HCDR3 conformation or shape plays a significant role in V2-apex bnAb function, we predicted the structures for these antigen-specific mAb HCs with long HCDR3s using the Colabfold pipeline ^76–78^. V2-bnAb HCDR3s fall within a spectrum of conformations resembling a needle (i.e., PGT145) or hammerhead (i.e., PG9) (Fig. S21). Both hammerhead- and needle-like HCDR3s were identified in the antigen-specific mAbs (Fig. 6C). Of the 52 mAbs, 45 were identified to contain hammerhead-like HCDR3 conformation, while 7 showed a needle-like shape. Overall, antigen-specific long HCDR3 mAbs were highly diverse in their sequence and IGHV gene usage, but shared features characteristic of human and rhesus V2-like bnAbs and their precursors.

**Figure 6.**
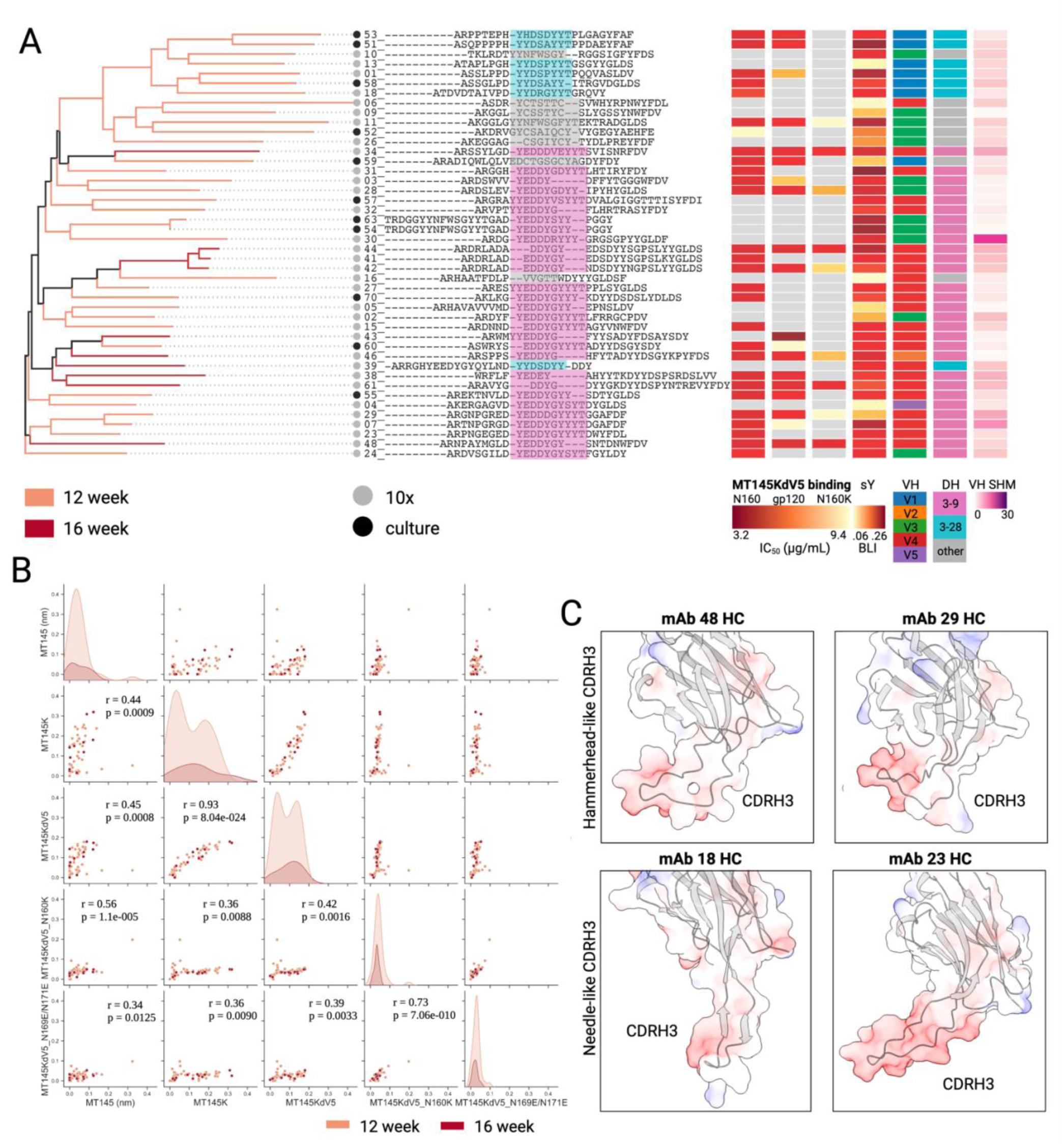
Select mAbs from antigen-sorted B cells from SCIV_MT145K.dV5 animals have V2 apex binding characteristics. (A) Phylogenetic tree for isolated antigen-binding mAbs expressed from B cells isolated at week 12 (tan) and 16 (maroon) time points for MT145K.dV5 animals. Grey and black circles at lineage terminals represent mAbs isolated using 10x Chromium or B cell culturing, respectively. HCDR3 sequences with IGHD regions highlighted in grey, pink, or aqua shown. IC_50_ values for mAbs against MT145K.dV5, MT145K.dV5-gp120, and MT145K.dV5-N160K shown in columns 1-3. Sulfotyrosine analysis using BLI (mean response) for each mAb shown in column 4. Column 5 and 6 show V gene and D gene usage, respectively. Last column shows a heatmap of V gene SHM levels per mAb. (B) Pairwise relationship across MT145 variants for all antigen-specific mAbs. The center diagonal subplots show histogram distributions for each MT145 variant, and the off-diagonal plots show binding correlation across different SOSIPs. All values obtained from BLI responses using mAb Fc capture biosensors and are from 12 and 16 wpi samples shown in tan and maroon, respectively. (C) Structural prediction of antigen-specific HC mAbs with hammerhead-like (top) and needle-like (bottom) HCDR3 conformations using ColabFold. Select mAb HCs 48 and 28 with hammerhead-like HCDR3s are shown in the top row and select mAb HCs 18 and 23 with needle-like HCDR3s are shown in the lower row.

## Discussion

Antigen-driven B cell selection and affinity maturation have been extensively studied for simple antigens but are less well-understood for complex antigens like the HIV-1 Env^79–81^. For example, haptens are often used to study clonal selection because they predominantly expand B cells with well-defined immunogenetic features and specificities to immunodominant epitopes^82–84^. However, for complex antigens such as the HIV-1 envelope glycoprotein, B cell responses emerge from intra-clonal and inter-clonal competition to multiple epitopes, and the dynamics of this competition is likely a key contributor to bnAb development^81,85^. Intra- and inter-clonal competition dynamics depend on BCR frequency in the repertoire, affinities, as well as avidities^19^. Here we show that B cell responses to germline targeting chimpanzee SIV MT145 Envelope glycoproteins can be enriched for desirable features present in the precursors of prototypical HIV-1 bnAbs. We show that unlike the wildtype chimpanzee MT145 Env, its germline-targeting trimer variants exhibit strikingly different immunogenetic properties across both total and antigen-specific repertoires by consistently eliciting long HCDR3 B cell lineages with paratope features characteristic for HIV-1 V2-apex bnAb precursors.

We previously showed that increasing the affinity of the MT145 Env to human V2-apex bnAb precursors resulted in epitope-specific nAbs following vaccination in a V2 apex bnAb precursor expressing knock-in (KI) mouse^50^. Here, we investigated the immunogenicity of three MT145 variants (MT145, MT145K, MT145.dV5) as replicating SCIVs in the rhesus macaque infection model. Although SCIV_MT145K elicited V2-targeted B cell responses, our data suggest that competition from the immunodominant V5 region may have reduced V2 apex directed responses. Partial trimming of the V5 region refocused these responses to the desired epitope although it did not completely eliminate V5 targeting. These findings suggest that eliminating or shielding immunodominant off-target epitopes may reduce interconal competition and focus the B cell response on more desirable epitopes. Indeed, germline-targeting and protein resurface engineering reduced off-target reactivities, favoring the engagement of rare V2-apex bnAb-like precursors with long HCDR3s. This was most pronounced for SCIV_MT145K.dV5 infected animals, where we observed a 10-fold increase in the fraction of long HCDR3 B cell lineages, some of which exhibited properties similar to prototypic human V2-apex bnAbs. Activation of rare long HCDR3 B cell precursors is a major bottleneck for inducing V2-apex bnAbs. Our study shows that this can be accomplished using both germline-targeting and immunogen resurfacing approaches.

In contrast to wildtype SCIV_MT145, infections with both MT145K and MT145K.dV5 SCIVs elicited selection in the V2 region and caused expansions of long HCDR3 antibody lineages with IGHD3-09 genes expressing the ‘EDDY’ motif. Both MT145K and MT145K.dV5 SCIVs expanded antibodies with V2-bnAb precursor-like features from the bulk repertoire, with more clonal expansion observed in the SCIV_MT145K.dV5 compared to the SCIV_MT145K infected group. These data suggest that while both germline-targeting Envs elicited responses to the V2 region, removal of the immunodominant V5 epitope increased the number of desired responses. It is possible that the precursor frequency and/or affinity of naive B cells targeting the MT145 V5 region was higher than that of B cells targeting the canonical V2 epitope, which would have given V5-targeting BCRs a competitive advantage over V2-targeting BCRs in the germinal centers. Although not statistically significant, there was a trend for improved V2-bnAb like responses in SCIV_MT145K.dV5 compared to SCIV_MT145K infected animals. Thus, it seems critical to eliminate off-target immunodominant B cell epitopes when designing germline-targeting immunogens.

The two germline-targeting SCIVs, but not their wildtype counterpart, appear to have primed V2 bnAb-like B cell precursors and elicited V2-directed antibody responses that placed weak, albeit transient, selection pressures on the C-strand. However, these responses failed to broaden since none of our SCIV infected macaques ultimately developed bnAbs. This failure may be due to the absence of the required viral variants in the germinal centers at the time of antibody maturation. A number of studies have shown that specific Ab-Env co-evolution pathways are needed for neutralization breadth to develop^9,53,86,87^. The lack of significant diversification within and around the core V2-apex epitope in our recovered Env sequences indicates that viral variants capable of driving affinity maturation down desirable pathways were lacking. The absence of key Env mutations in the majority of the circulating viral immunotypes may have impeded the maturation of nascent bnAb lineages. Alternatively, it is possible that the SCIV_MT145K and SCIV_MT145k.dV5 expanded B cell lineages, despite their similarity to known V2 apex bnAb UCAs, have additional features that prevent them from maturing to breadth even if the necessary boosting immunotypes are present. Future studies will need to differentiate between these scenarios.

The SCIV_MT145K.dV5 induced long HCDR3 mAbs exhibited a range of V2-apex epitope specific binding and neutralizing properties as well as immunogenetic features frequently found in V2 apex bnAbs. It is thus tempting to speculate that a germline-targeting immunogen that engages a large pool of long HCDR3 lineages with diverse paratope properties, including those with features similar to V2-apex bnAbs, would also prime desired B cell responses to this bnAb site. Once a sizable pool of long HCDR3 precursor B cells is engaged, boosting immunogens capable of immunofocusing and polishing are required. Rhesus macaque appears to represent a particularly suitable outbred animal model to test V2 apex bnAb induction strategies since their B cell repertoires possess precursors with paratope properties that closely resemble human V2-apex bnAb lineages^30^. In addition, the SHIV infection model allows for challenge/protection studies should the desired bnAbs be induced by vaccination^53,88^. Thus, rhesus macaques provide a rapid and reliable evaluation model to test proof-of-concept germline-targeting, immunofocusing, and polishing vaccine strategies for the HIV V2-apex bnAb site, which can inform HIV vaccine trials in humans.

In summary, our study illustrates that rare bnAb-like B cell precursors can be preferentially stimulated by germline targeting immunogens, which represents a first step in bnAb elicitation. Moreover, rational antigen design may allow to eliminate or shield off-target epitopes that would otherwise impede bnAb development. Although the SCIV infected RMs failed to develop bnAbs, these results provide new insights into strategies for HIV-1 vaccine development.

## Limitations of the study

Although we have demonstrated rare bnAb-like B cell precursors are expanded in germline targeting Envs using the SCIV infection model, protein immunization with the same immunogen might be different. Inducing similar responses in rhesus macaques by protein immunizations would be ideal for additional *in vivo* comparisons. An important variable for B cell expansion is

the concentration and retention of antigen in draining lymph nodes and the periphery. Although we see viral kinetics were consistent across animals for each group, slight differences across groups may differentially affect antigen availability and in turn influence B cell responses. Thus, evaluation of protein immunizations with fixed concentration should clarify the effect of antigen concentration on B cell responses. In this study, we used immunogenetic and biochemical signatures of rhesus macaque V2 bnAbs to categorize SCIV-elicited mAbs as precursor bnAbs. However, whether any and which of the expanded lineages have the potential to mature into real bnAbs is still unclear. Further studies using SHIVs and SCIVs to interrogate Envs that promote lineage maturation would help clarify the limitation of our study.

## ACKNOWLEDGEMENTS

We thank … for expert technical assistance; T. Denny, T. Demarco, and N. DeNaeyer and members of the Nonhuman Primate Virology Core Laboratory at the Duke Human Vaccine Institute for SCIV plasma vRNA measurements; the staff at Bioqual for exceptional care and assistance with nonhuman primates. This work was supported by National Institutes of Health grants R61 AI 161818 (R.A., G.M.S.), R01 AI 167716 (R.A.), R01 AI 050529 (B.H.H.), R37 AI 150590 (B.H.H., K.W., B.T.F), R01 AI 160607 (G.M.S.), P01 AI 131251 (G.M.S., K.W.), the University of Pennsylvania Center for AIDS Research (P30 AI 045008), the National Institute of Allergy and Infectious Diseases (NIAID) Consortium for HIV/AIDS Vaccine Development (CHAVD; UM1 AI144371, UM1AI144462) (A.B.W., D.S., B.B., and D.R.B.) .), the Bill and Melinda Gates Foundation through the Collaboration for AIDS Vaccine Discovery (INV-008352/ OPP1153692 and OPP1196345/INV-008813) (D.S., D.R.B., A.B.W., and R.A.), and (OPP1206647) (G.M.S., and R.A.). R.S.R. and R.M.R. were supported by a training grant (T32 AI 007632). R.M. was supported by a training grant (T32 AI 007244).

## Author Contributions

R.M., Y.S., G.S., R.S.R., G.M.S., and R.A. conceived and designed experiments; R.M., G.S., R.S.R., F.L., S.W., R.M.R., W.D., Y.L., J.R., A.I.M., E.L., C.Z., A.J.C., F.B., N.M., generated SCIV viruses, performed neutralization and binding assays; R.M., G.S., R.S.R., F.L., S.W., R.M.R., W.D., and Y.L. collected viral Env sequencing, repertoire, and single cell sequencing data; R.M., G.S., and P.Y. performed single B cell sorting and B cell culture experiments; R.M., G.S., P.Y., G.A., W.H., S.C., K.D., A.L.V., X.L., T.C., F.A., cloned, expressed, purified, and tested monoclonal antibodies and SOSIP trimers and probes; S.Z., W.L., G.O., and A.B.W. generated and analyzed EMPEM data; Y.S., R.M., J.H., and B.B. analyzed repertoire and single cell NGS data; B.T.F., K.W., Y.S., R.S.R., and R.M. analyzed viral envelope sequencing data; the experiments. N.M., and R.M. performed the structure prediction experiments. R.M., Y.S., E.L., B.H.H. and R.A. wrote the manuscript with input from all listed authors.

## Key Resource Table

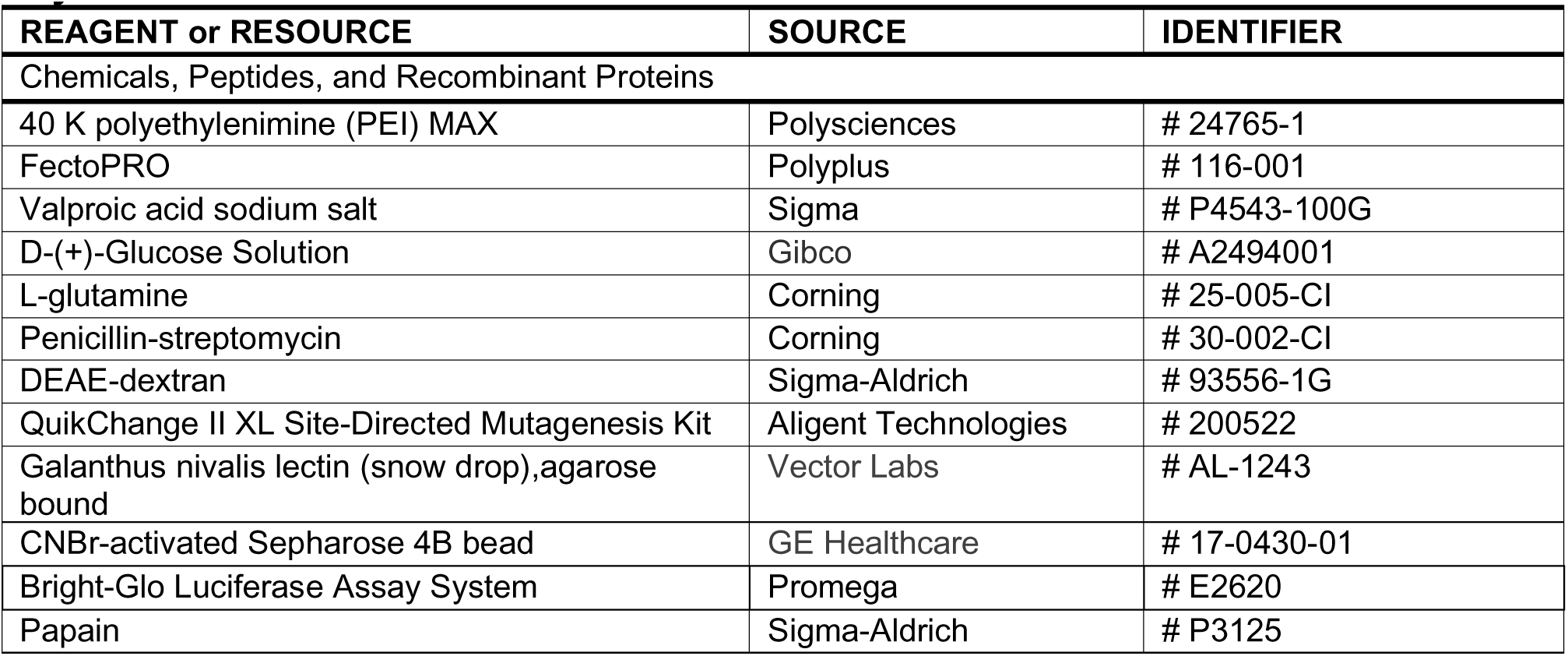
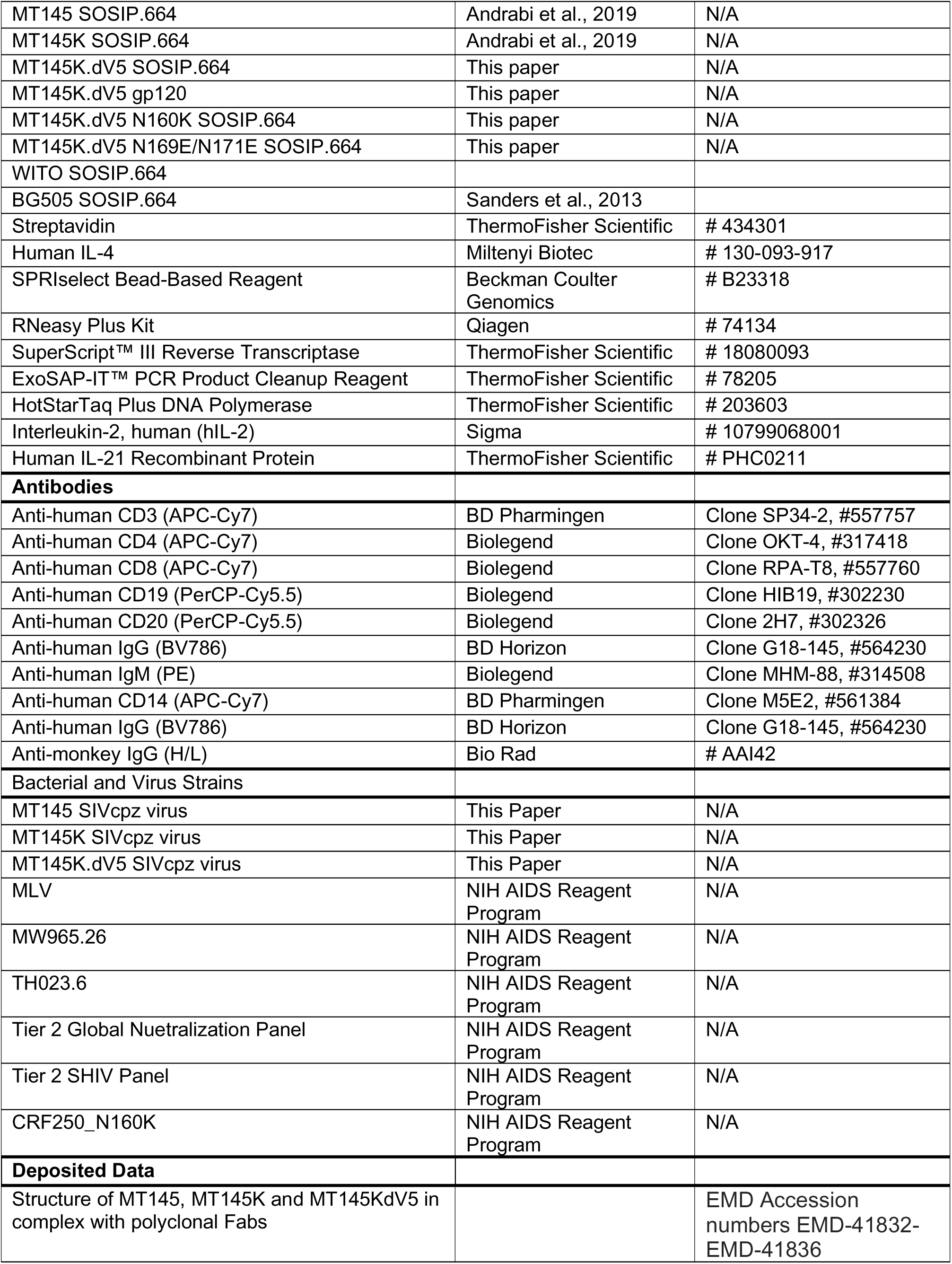
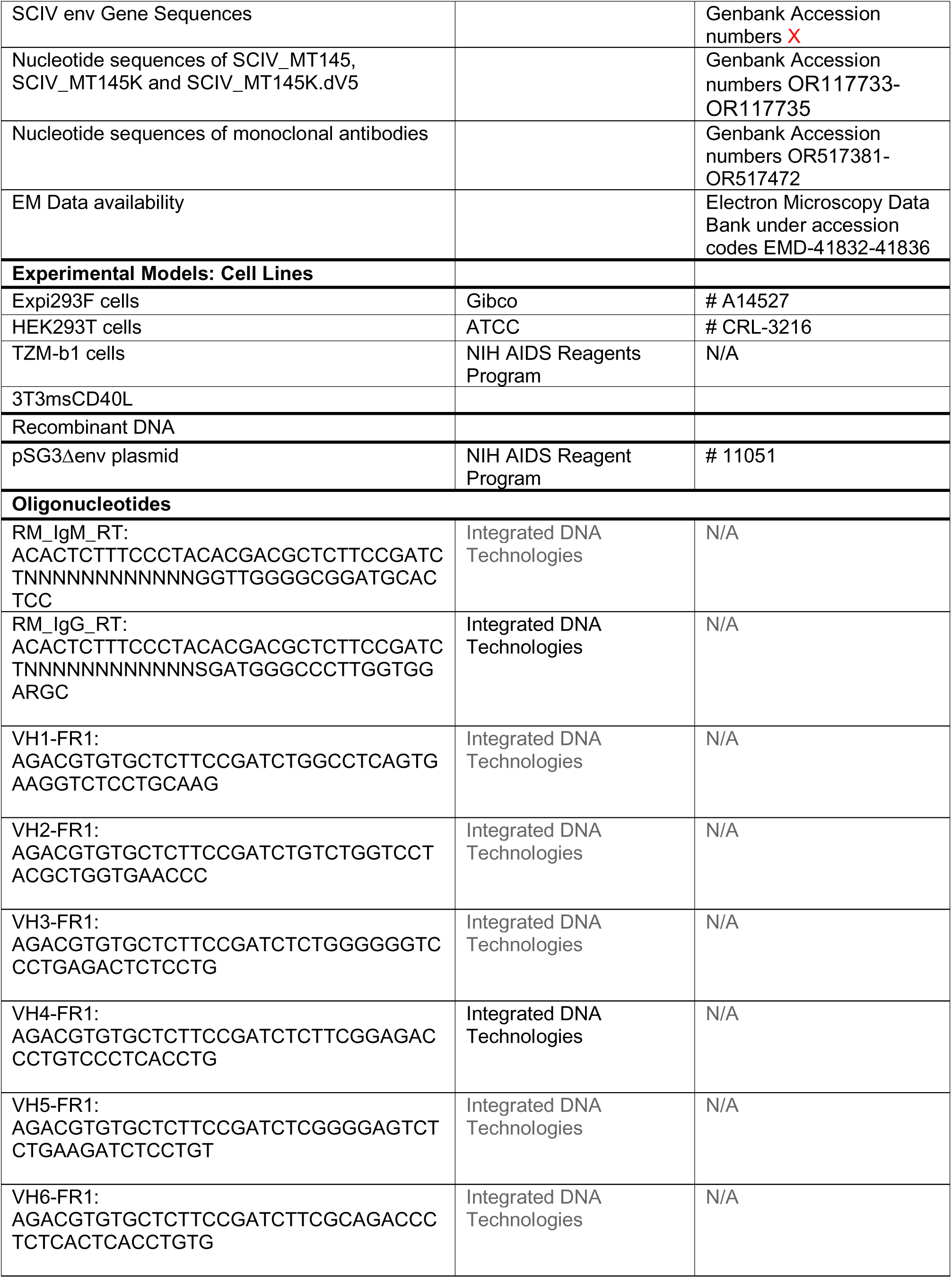
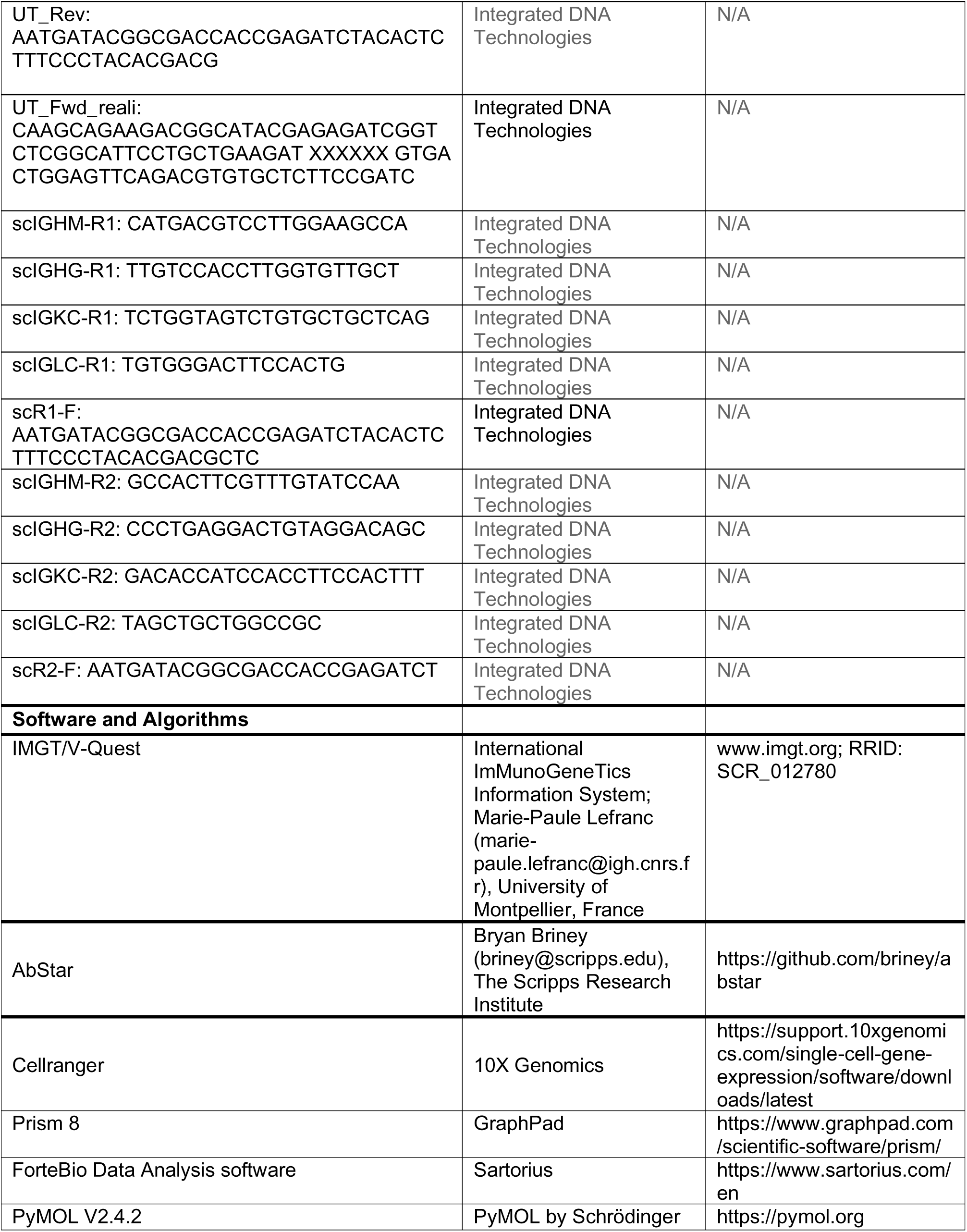
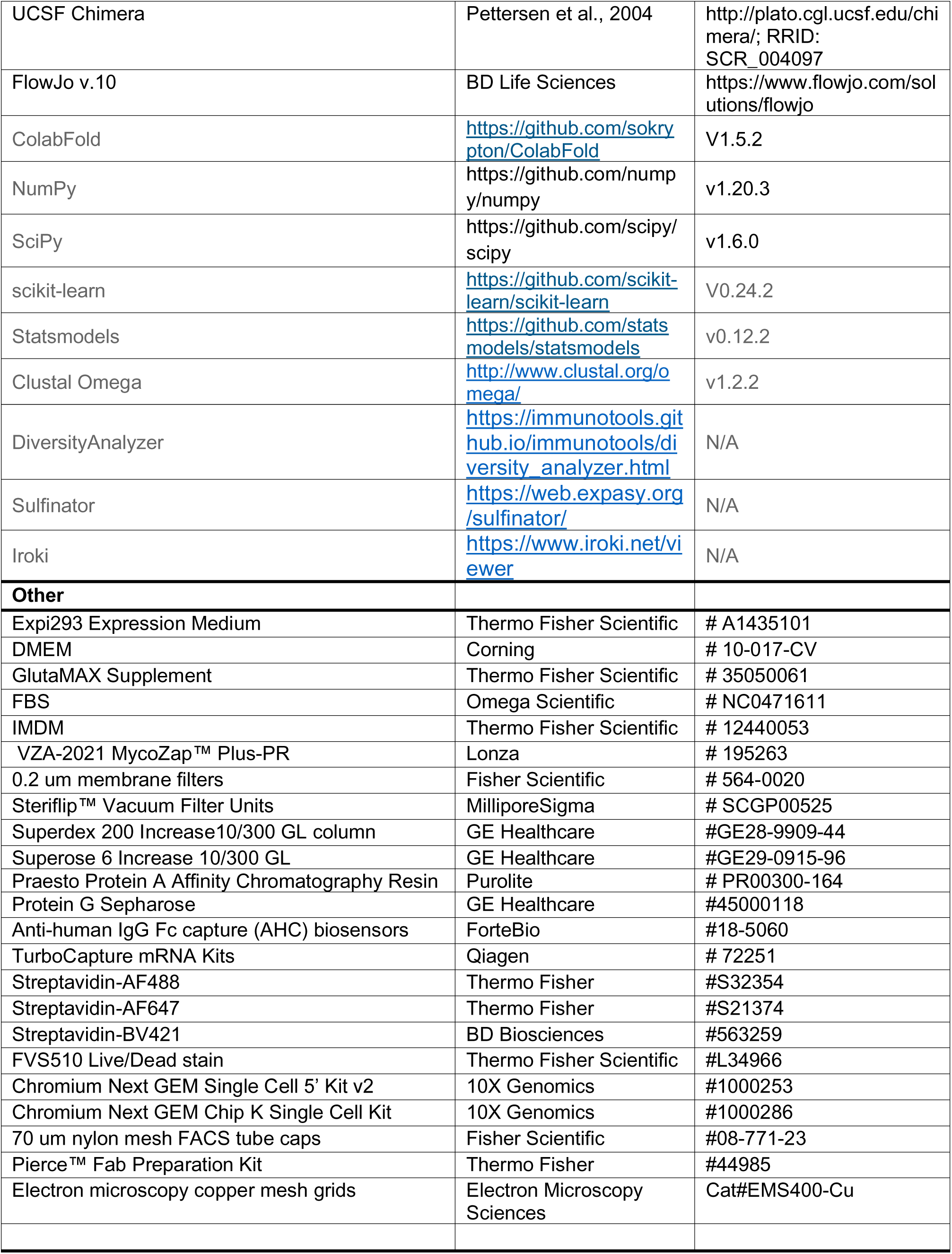

## EXPERIMENTAL MODEL AND SUBJECT DETAILS

### Animal studies

Nine outbred Indian rhesus macaques (Macaca mulatta) were used in this study. All animals were housed in Bioqual, Inc. (Rockville, MD) according to guidelines set by the Association for Assessment and Accreditation of Laboratory Animal Care International (AAALAC). All experiments were approved by the University of Pennsylvania and Bioqual Institutional Animal Care and Use Committees (IACUC). Animals were socially housed with a variety of recommended environmental enrichments and sedated for blood draws, anti-CD8 mAb infusions and SCIV inoculations. RM 6931 received 1 ml of an equal mixture of three SCIV constructs (50 ng of p27 antigen), all of which expressed the wildtype MT145 Env but differed in their SIVmac backbone. RM 6932 and RM 6933 were infected with a mixture of six allelic variants of SCIV_MT145, which differed at Env position 375 (50ng of p27 antigen each). RMs T922, T923, and T924 received 1 ml of SCIV_MT145K, which contained a glutamine (Q) to lysine (K) mutation at position 171 in the C strand as previously described ^89^. RMs 43189, 43359, and 44092 received 1 ml of SCIV_MT145K.dV5, which had a shortened V5 loop. All three SCIVs encoded the wildtype His at position 375 of the envelope glycoprotein. Plasma viral loads were determined as previously described^40,53^. All animals received an intravenous infusion of 25 mg/kg of the anti-CD8b mAb CD8beta255R1 at the time of SCIV inoculation. Infected RMs were followed for up to 88 weeks, with plasma and peripheral blood mononuclear cells (PBMCs) collected at weekly and bi-weekly (Fig 1B) and later at monthly and bi-monthly intervals (Fig S1). Animals 6931, 6933 and T924 progressed to AIDS within 29, 18 and 22 weeks, and were thus euthanized on day 203, 128 and 154, respectively.

### Cell Lines

Expi293F cells (Gibco Cat# A14527) were maintained in Expi293 Expression Medium (Gibco Cat# A1435101) at 37°C in a 8% CO_2_ atmosphere 150 rpm shaker. HEK293T (ATCC Cat# CRL-3216) cells were grown in Dulbecco’s Modified Eagle Medium (DMEM) (Corning Cat# 10-017-CV) with 10% heat-inactivated FBS (Omega Scientific Cat# NC0471611), 4mM L-Glutamine (Corning Cat# 25-005-CI) and 1% P/S (Corning Cat# 30-002-CI) in a 37°C, 5% CO_2_ incubator. Irradiated 3T3msCD40L cells were used in B cell culture assay and TZM-b1 cells (NIH AIDS Reagents Program) were used for the pseudovirus neutralization assay as previously described.

## METHOD DETAILS

### SCIV construction

To generate the wildtype SCIV_MT145 construct, we cloned a *vpu-env* fragment (*env* nucleotides 1 to 2153, HXB2 numbering) from the SIVcpz MT145 infectious molecular clone (JN835462) into first-generation^90^, “intermediate,” and second-generation SIVmac vectors^70^ and tested their relative replication potential by infecting RM 6931 with an equal mixture (based on p27 content) of these constructs. Sequence analysis 2 weeks post-infection identified the second-generation vector as the biologically most fit, which was thus selected for all subsequent constructions. Since the amino acid residue at position 375 of the HIV-1 Env determines how efficiently the corresponding SHIV replicates in rhesus CD4+ T cells, we next created isogenic mutants of SCIV_MT145 by changing the wildtype histidine (375H) to serine (375S), tyrosine (375Y), methionine (375M), tryptophan (375W) or phenylalanine (375F). After assessing their replication competence in rhesus CD4+ T cells *in vitro*, these constructs were used as equal mixtures (based on p27 content) to infect RM 6932 and RM 6933.

Single genome sequencing of plasma viral RNA 14 and 28 days post-infection identified the 375H variant as the predominant strain in both animals. The 375H construct was then used to generate the germline-targeting SCIV_MT145K by replacing a glutamine (Q) at position 171 in the C strand of SCIV_MT145 with a lysine (K) residue as previously described^50^. The second germline-targeting SCIV_MT145K.dV5 was generated by deleting nine amino acids (NREDQGEDQ) from the V5 loop of SCIV_MT145K, which was identified as a strain-specific immunodominant off-target epitope. The germline-targeting potential of the SCIV-expressed MT145K and MT145K.dV5 Envs was confirmed by testing their sensitivity to neutralization by inferred human V2 apex bnAb precursors (Fig. S5). The nucleotide sequences of SCIV_MT145, SCIV_MT145K and SCIV_MT145K.dV5 are available under GenBank accession numbers OR117733, OR117734and OR117735, respectively.

### Neutralizing antibody assay

The neutralization capacity of rhesus macaque plasma was assessed using the TZM-bl assay as described^91^. Briefly, serial 5-fold dilutions of RM plasma (1:20, 1;100, 1:2,500, 1:12,500, 1:62,500, 1:312,500) were incubated with transfection-derived virus at a multiplicity of infection of 0.3 in a total volume of 100 ml in the presence of DEAE-dextran (Signma-Aldrich) (40 mg/ml) for 1 h at 37°C, and this mixture was then added to TZM-bl cells. After 48 h, TZM-bl cells were analyzed for luciferase expression with uninfected cells used to correct for background luciferase activity. The infectivity of each virus without plasma was set at 100% and the plasma dilution that reduced the relative light units (RLUs) by 50% compared with the no plasma control wells were calculated by using the variable slope (four parameters) function in Prism software (v8.0). Viral stocks were generated by transfection of 293T cells, using 4.5 μg of and Env-minus HIV-1 (SG3Δenv) (NIH AIDS Reagent Program) backbone and 30 ng of codon optimized HIV-1 Env plasmids, or 6 μg of SCIV or SHIV construct DNA. B cell culture supernatants and mAbs were incubated with Env-encoded pseudoviruses in 384-well plates (Greiner Bio-One) for 1 h at 37°C. TZM-bl cells were added at 5,000 cells/well in 50 *μ*L of complete DMEM and incubated for an additional 48 h at 37°C. After incubation, culture media was removed, cells were lysed, and luciferase activity was read after Bright-Glo (Promega) addition.

### Env SOSIP Expression and Purification

SOSIP Envs were expressed and purified as previously described^50^. Briefly, plasmids encoding for HIV Env SOSIP trimers were cotransfected in HEK293F cells using PEI-MAX 4000 transfection reagent (Polysciences, Inc.). After transfected cells were incubated for four days, supernatants containing expressed trimers were placed in an agarose-bound Gallanthus Nivalis Lectin (GNL) or CNBr-activated Sepharose 4B bead (GE Healthcare) bound PGT145 bnAb antibody affinity columns for purification. Size exclusion chromatography (SEC) purification in a Superdex 200 10/300 GL column (GE Healthcare) in PBS/TBS was performed for further purification if needed.

Introduction of specific mutations in the SOSIP Envs was performed using QuikChange site-directed mutagenesis kit (Agilent Technologies, USA) according to manufacturer’s instructions and verified by sequencing analysis (Eton Bioscience, San Diego, CA).

### Monoclonal Antibody Expression and Purification

Plasmids containing HC and LC sequences were expressed in Expi293F cells. Briefly, FectoPRO (Polyplus) transfection reagent was used to cotransfect both HC and LC plasmids according to manufacturer’s instructions. Transfected cells were incubated at 37°C, 5% CO_2_ with 150 rpm shaking. One day after transfection, cells were given 300 mM valproic acid and 40% glucose (Gibco). After 5 days of incubation, supernatant was filtered through a 0.22 mm Steriflip (EMD Millipore) to remove cells. To purify the antibodies, the filtered supernatant ran through a protein A and protein G affinity column (GE Healthcare) and eluted with 0.2 M citric acid at pH 3.0 and neutralized in 2 M Tris-base. Antibodies were buffer-exchanged into phosphate-buffered saline (PBS). The nucleotide sequences of mAb HCs and LCs are available under GenBank accession numbers OR517427-OR517472 and OR517381-OR517426, respectively.

### B Cell Sorting

For antigen-specific sorting, Avi-tagged biotinylated MT145, MT145K, and MT145K.dV5 trimers were conjugated with SA-labeled fluorophores at RT for 30 min. Cryopreserved PBMC samples were thawed and added into RPMI medium (Thermo Fisher) supplemented with 50% FBS. Cells were washed with FACS buffer (PBS with 2% FBS) and stained using anti-CD3, CD4, CD8, CD14, CD19, CD20, IgG, and IgM fluorescent antibodies for 15 min at RT in the dark. After incubation, conjugated antigens were added to stained cells and incubated for an additional 30 min. A 1:300 dilution of FVS510 LIVE/DEAD cell stain (Thermo Fisher Scientific) was added to the sample and incubated for an additional 15 min. Prior to sorting, cells were washed with FACS buffer and filtered through a cell strainer in a 5 mL round bottom tube (Corning). Sorting was performed on a BD FACSMelody and cells were either sorted in tubes for single cell sequencing or incubated in 384-well plates for B cell culturing.

### BioLayer Interferometry (BLI) Binding Assay

Binding by BLI was performed using an Octet K2 system (Sartorius) using anti-human IgG-Fc biosensors (AHC: ForteBio). To set up the assay, 10 *μ*g/mL of IgGs and 200 *μ*M of SOSIP was used for loading and capture, respectively. Biosensors were first dipped into PBST and loaded with IgGs for 60 s. The loaded biosensors were then incubated with Env for 120 s. To measure off rates, biosensors with captured protein was placed in PBST for 240 s. Analysis was performed using the ForteBio Data Analysis Software 10.0 and plotted using Prism 8.

### Antibody-Env ELISA Binding Assay

ELISA assays were performed using biotinylated proteins on streptavidin coated plates as previously described^50^. Briefly, 2 *μ*g/mL streptavidin (Thermo Fisher Scientific) was used to coat 96-well half-area clear plates (Corning, Thermo Fisher Scientific) overnight. Plates were washed 3 times with PBST and blocked with 3% BSA in PBS for 1 h. Biotinylated proteins were added at 2 *μ*g/mL in 1% BSA in PBST and incubated for 1.5 h at RT. After protein incubation, plates were washed three times and diluted mAbs were added for an additional 1.5 h. Secondary antibodies conjugated to alkaline phosphatase (Jackson ImmunoResearch Laboratories) was added following staining with alkaline phosphatase substrate pNPP (Thermo Fisher Scientific). Absorbance was measured after 20 minutes at 405 nm using a VersaMax microplate reader (Molecular Devices) and plotted using Prism 8.

### Negative Stain Electron Microscopy

The negative stain EMPEM method was described previously^92^. Briefly, 15 µg of trimer MT145, MT145K or MT145K.dV5 was incubated overnight with 0.5 mg polyclonal Fab (generated using papain digestion) and purified the next day using a Superdex 200 Increase 10/300 GL gel filtration column (Cytiva). After concentrating, purified complexes were diluted to 0.03 mg/mL and deposited on glow-discharged carbon coated copper mesh grids, followed by staining with NanoW (Nanoprobes). Imaging was performed on an FEI Tecnai Spirit T12 equipped with an FEI Eagle 4k x 4k CCD camera (120 keV, 2.06 Å/pixel), an FEI TF20 equipped with a TVIPS TemCam F416 CMOS 4k x 4k camera (200 keV, 1.77 Å/pixel) and an FEI Talos Arctica equipped with an FEI Ceta (4k x 4k) camera (120 keV, 1.98 Å/pixel). All data were processed using Relion 3.0 (PMID: 30412051) using standard 2D and 3D classification procedures. Composite maps were generated using UCSF Chimera (PMID: 15264254). Representative maps have been deposited to the Electron Microscopy Data Bank (see STAR Methods).

### Single Genome Amplification and Env Evolution Analysis

Single genome amplification of viral RNA was performed as described previously^53^. Briefly, ∼20,000 copies of viral RNA were extracted from plasma using QIAamp Viral RNA kit (Qiagen) and reverse transcribed using SuperScript III Reverse Transcriptase (Invitrogen). Viral cDNA was then endpoint diluted and 3’ half genomes or viral *env* genes were amplified using nested PCR with primers and conditions as previously reported (refs). Geneious software was used for alignments and sequence analysis (see Table X for GenBank accession numbers of longitudinal *env* gene sequences).

### Bulk Repertoire Sequencing

Cryopreserved PBMC samples were thawed at 37°C and added into RPMI medium (Thermo Fisher) supplemented with 5% FBS. Cells were centrifuged at 400 x g for 5 min, and the supernatant was discarded. Cells were lysed and RNA was extracted using the RNeasy Plus Mini Kit (Qiagen). Extracted RNA was used to synthesize cDNA using IgM and IgG reverse transcription primers and SuperScript III enzyme (Thermo Fisher Scientific). cDNA products were cleaned using ExoSAP-IT (Thermo Fisher Scientific) following a 2-step VDJ amplification using HotStarTaq Plus (Qiagen). PCR products were enzymatically cleaned again and illumina adapters and indexes were introduced via PCR. The final DNA libraries were SPRI-cleaned (SPRIselect, Beckman Coulter Genomics) and quantified by concentration (Qubit, Thermo Fisher Scientific) and size (Bioanalyzer, Agilent 2100). Libraries were loaded on a Illumina MiSeq system using 2 x 300 bp read length.

### Single B Cell Sequencing

Antigen-enriched B cell repertoires were sequenced using the 10x 5’V2 Single Cell Immune Profiling kit per manufacturer’s instructions (10x Genomics) with the modification of custom NHP primers for V(D)J amplification steps. Libraries were loaded on a Illumina MiSeq system using 2 x 300 bp read length and sequences were identified and analyzed using Cell Ranger.

### B Cell Activation Assay

Antigen-specific B cells were sorted in 384-well plates for B cell culturing as previously described (Zhao et al. 2022). Briefly, B cells were cultured with Iscove’s modified Dulbecco’s medium (IMDM) supplemented with GlutaMAX (Gibco) and 10% FBS, 1× MycoZap Plus-PR (Lonza), 100 U/mL human IL-2 (Roche), 50 ng/mL human IL-21 (Thermo Fisher Scientific), 50 ng/mL human IL-4 (Miltenyi), 0.1 *μ*g/ mL anti-rhesus IgG (H + L) (BioRad), and irradiated 3T3msCD40L feeder cells. After 14 days of incubation at 37°C, the supernatant was transferred to a new, sterile 384-well plate for binding and neutralization experiments. Remaining B cells were lysed and mRNA was extracted using TurboCapture plates (Qiagen) according to manufacturer’s instructions. RT-PCR was performed using a Superscript IV reaction and IgH, IgK, and IgL primers. Paired HC and LC sequences were amplified using nested PCR reactions and analyzed by 2% 96 E-gels (Thermo Fisher Scientific) followed by sequencing (Eton Bioscience, San Diego, CA).

### Immunogenomics analysis

Repertoire sequencing reads were merged and processed using the DiversityAnalyzer^93^tool. The macaque database of germline immunoglobulin genes reported by^75^ was used for alignment of repertoire sequencing reads and CDR labeling. VDJ sequences that have identical V and J gene matches and HCDR3s of the same lengths with at least 90% similarity were combined into clonal lineages. Posttranslationally modified tyrosine sulfation sites were predicted using the Sulfinator tool^94^with the maximum E-value threshold equal to 30. Phylogenetic trees of the clonal lineages were computed using the Clustal Omega tool^95^ and visualized using the Iroki tool^96^). Statistical tests were performed using the following Python packages: NumPy (v1.20.3), SciPy (v1.6.0), scikit-learn (v0.24.2), statsmodels (v0.12.2).

### Structure Prediction and Visualization

The structure for heavy chain sequences of all expressed mAbs from infected RMs, PGT145, and PG9 were predicted with ColabFold^76^. The alignment was prepared using MMseqs2^78^ PDB100 was used as the template database. The structure prediction was carried out with alphafold2_ptm as the model with 3 recycles and max MSA of 512:1024. The structures were visualized in UCSF ChimeraX^77^.

### Env SGS, NGS, and LASSIE analysis

SGA sequence alignment and analyses using LASSIE and hypervariable loop characteristics was performed as previously^53^. Briefly, Gene Cutter (https://www.hiv.lanl.gov/content/sequence/GENE_CUTTER/cutter.html) from the Los Alamos HIV Database was used to isolate the Env genes and an automated codon-alignment spanning all time points from all RMs was obtained, which was with further refined by manual curation (particularly in the hypervariable loop regions). LASSIE ^97^ was used to identify Env sites under putative selection pressure using a pre-specified criterion of 50% or more mutation away from the TF sequence at any time point in each RM. Hypervariable V5 loop characteristics for each time point from each RM were calculated using Variable Region Characteristics webtool from the Los Alamos HIV Database (https://www.hiv.lanl.gov/content/sequence/VAR_REG_CHAR/index.html) by assuming that hypervariable V5 loop spans from HXB2 site 460 up to 465. Sequence logos were generated using Logomaker ^98^ (https://logomaker.readthedocs.io/en/latest/).

Given the volume of NGS sequences, it was not possible to use standard multiple alignment tools even on sequences from the same RM and the same time point, and thus, the following modified strategy was used. MACSEv2 ^99^ (https://www.agap-ge2pop.org/macse/) was used to individually align each NGS sequence to the TF V1V2 region, using a custom Python script that both initiated these runs and analyzed the results of each alignment. The vast majority of NGS sequences had the same length as the TF or only showed deletions, both of which led to straightforward alignment of the NGS sequences to TF and to each other. For a minority of sequences (around 300-800 sequences out of ∼44,000-136,000 of week 12 sequences from MT145 group), there were sizeable insertions (including frameshift mutations) to rule out straightforward alignment; these were isolated and subjected to baseline alignment using MACSEv2 followed by manual refinement. In these sequence alignments, around 10-30 sequences per RM were found to have no homology to MT145 sequence, and were found to map to other organisms using BLAST ^100^; these reads were removed. The simpler alignment (without any insertions) and the more complex alignments (with insertions) were then merged together by matching the reference TF sequence from each alignment to obtain the full alignment for all NGS sequences from the same time point for the same RM. These alignments were subjected to a custom Python script to calculate per time point mutation frequency at each site in the V1V2 region for each RM, and mutation frequencies at select V2 apex sites were compared using two sided Wilcoxon rank sum test as implemented in Scipy (v0.18.0)(www.scipy.org) and plotted using Matplotlib (v.1.4.2)(www.matplotlib.org).

### EM Data availability

The structures presented in this manuscript can be found in the Electron Microscopy Data Bank under accession codes EMD-41832, EMD-41833, EMD-41834, EMD-41835, and EMD-41836. Heavy chain antibody repertoire sequencing reads were deposited to NCBI under accession number PRJNA1014130.

---

**Supplementary Figure 1.**
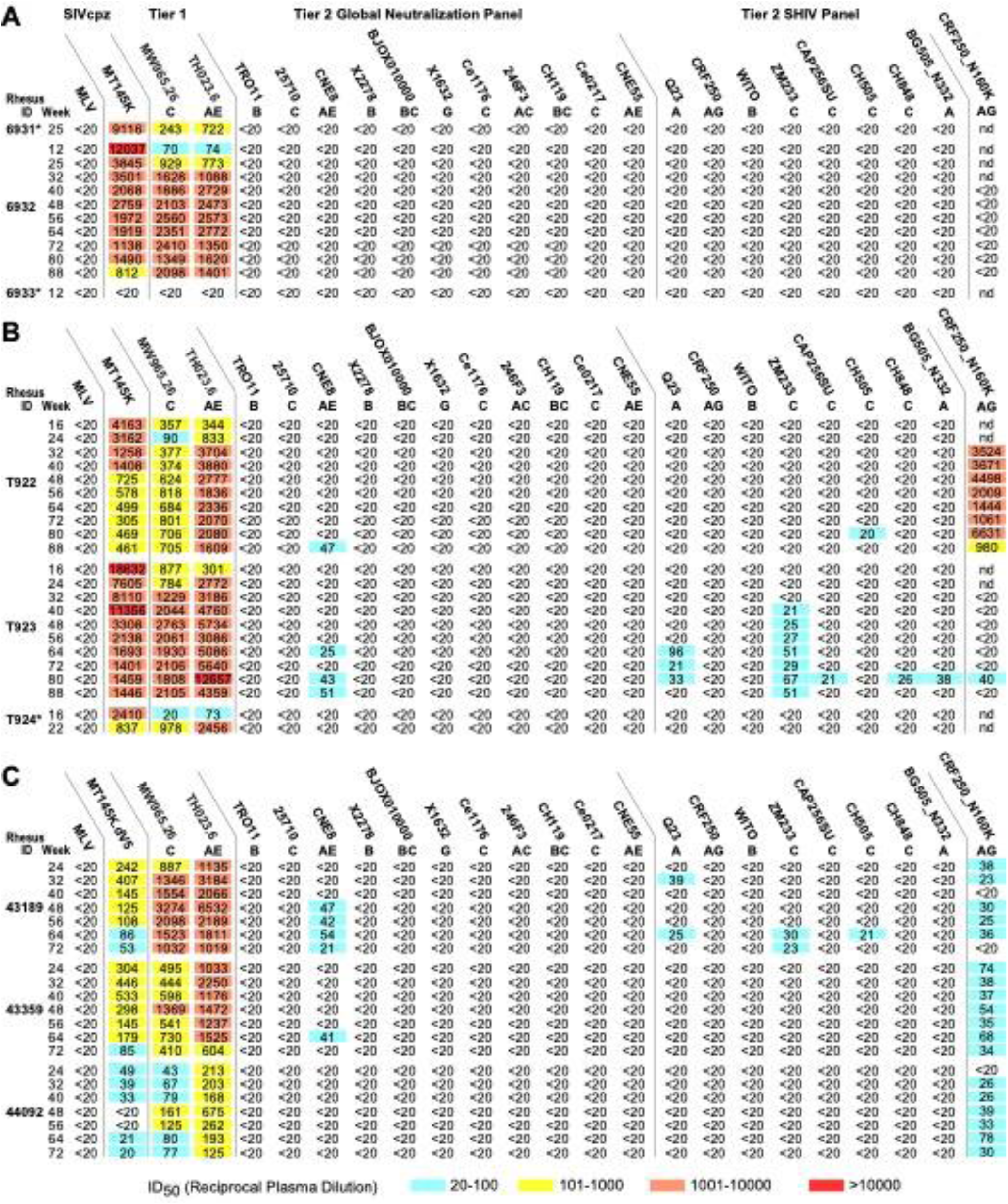
Autologous and heterologous neutralization of longitudinal plasma samples from RMs infected with (A) SCIV_MT145, (B) SCIV_MT145K and (C) SCIV_MT145K.dV5. Reciprocal 50% inhibitory dilutions (ID_50_) are shown for autologous (SCIV_MT145K and SCIV_MT145K.dV5) and heterologous (tier 1 and tier 2) viruses representing different HIV-1 subtypes (A, AG, AE, AC, B, C, BC, G; indicated below virus name), with no reactivity observed against a murine leukemia virus (MLV) Env control (all ID_50_ <1:20). One N160K mutant was also tested (CRF250_N160K). Coloring indicates relative neutralization potency. Both Env pseudoviruses (tier 1 and tier 2 global panel) and replication competent SHIV strains were tested. Asterisks denote rapid progressor animals that were euthanized prior to week 88.

**Supplementary Figure 2.**
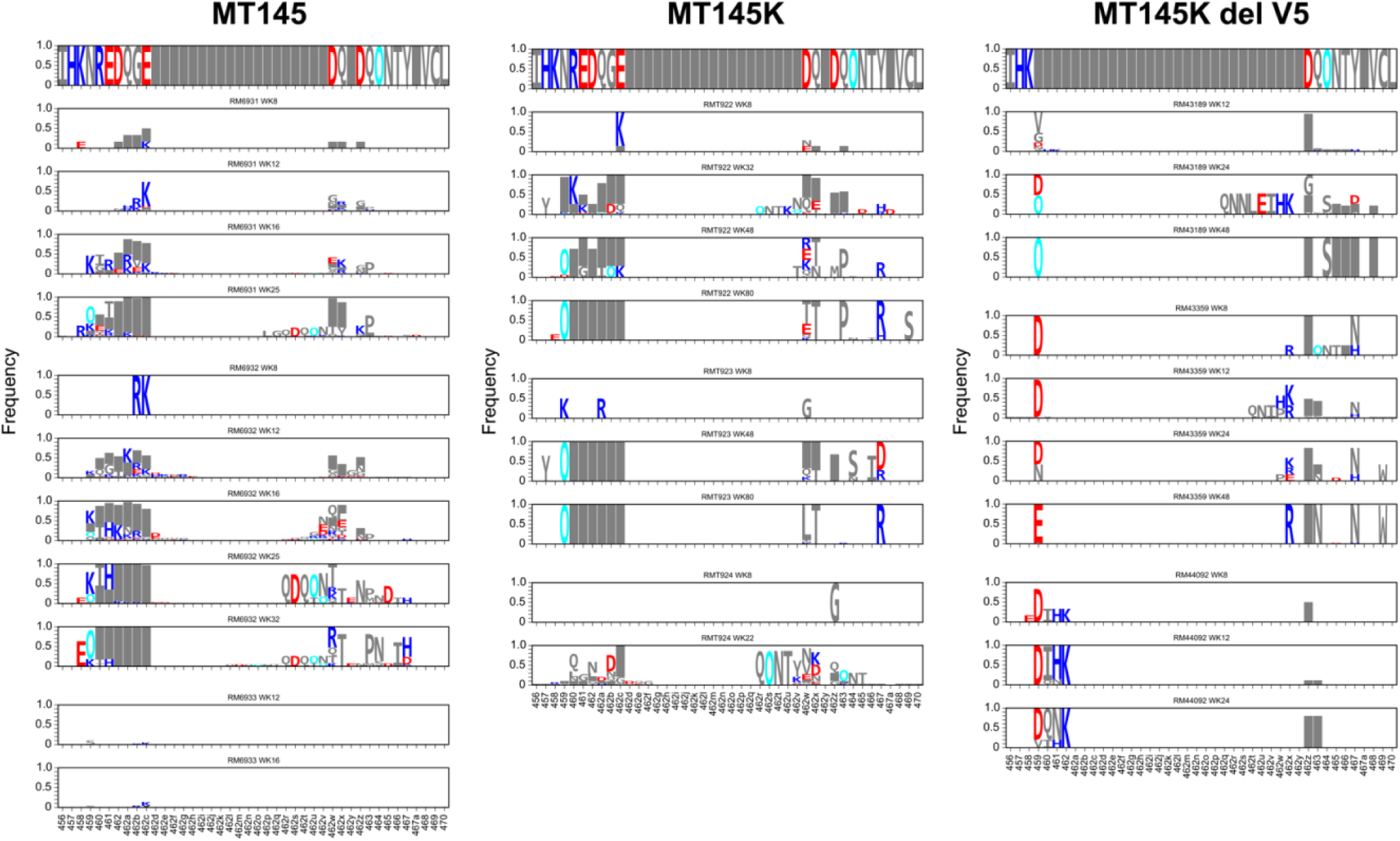
Env evolution in the V5 region in MT145 (left), MT145K (center), and MT145K.dV5 (right) SCIV infected rhesus macaques. Sequence logos show the per time-point frequency of Env mutations away from the TF sequence using the SGS sequences from each RM. The horizontal axis labels are HXB2 numbers of sites, with gaps relative to HXB2 indicated by letters appended to the immediately preceding HXB2 number (e.g. 462a). O=glycan. Colors: O=cyan, DE=red, HRK=blue.

**Supplementary Figure 3.**
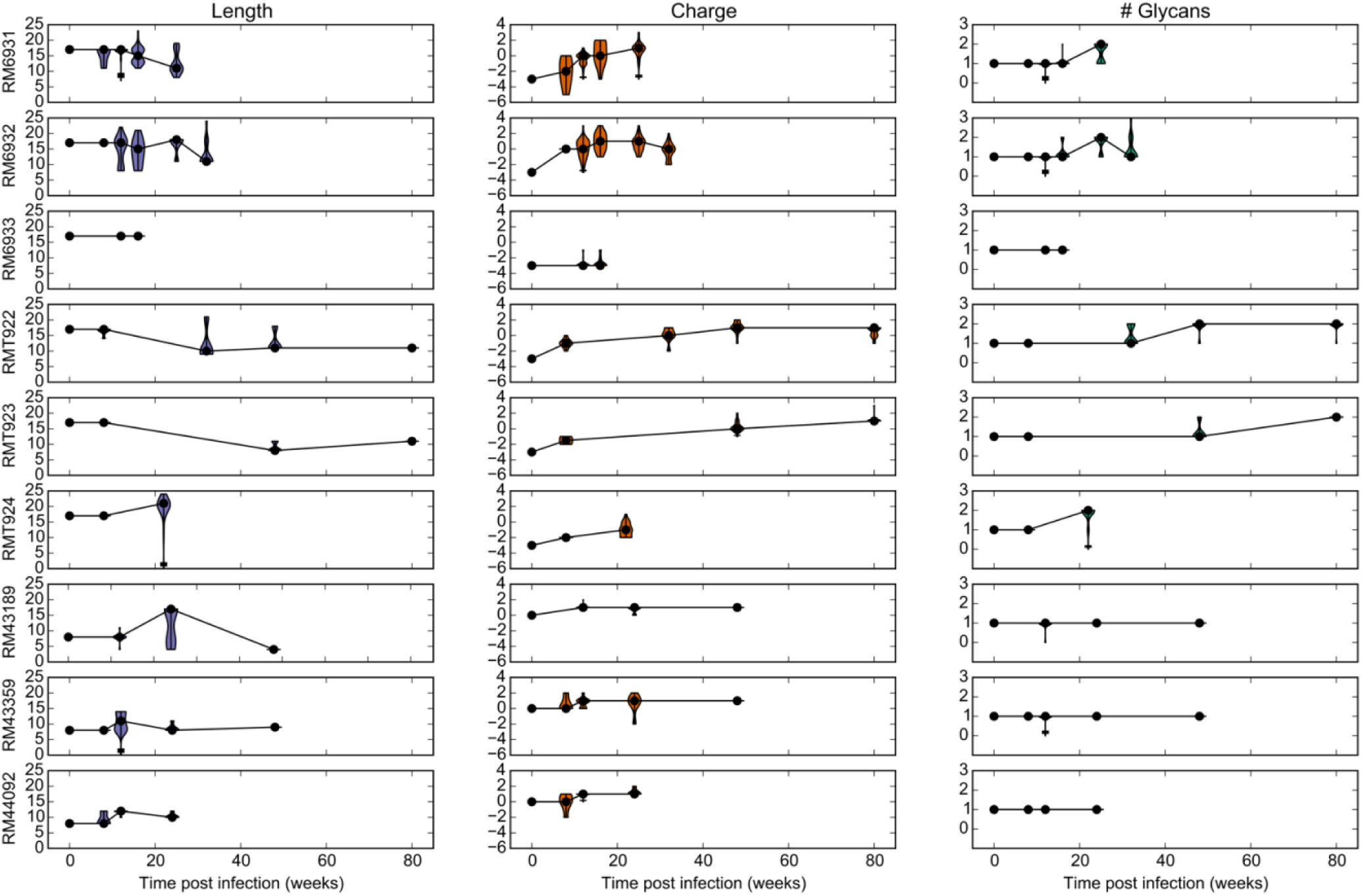
Longitudinal alignment-free hypervariable V5 loop characteristics for all three groups. Changes in V5 length (purple, left), charge (orange, center), and glycans (green, right) in the hypervariable V5 loop were determined by SGA sequence analysis. The distributions of each characteristic for sequences from each time point and each RM are shown using violin plots with black dots indicating medians, the vertical lines indicate the inter-quartile range (25^th^ to 75^th^ percentile). Median points are connected with black lines.

**Supplementary Figure 4.**
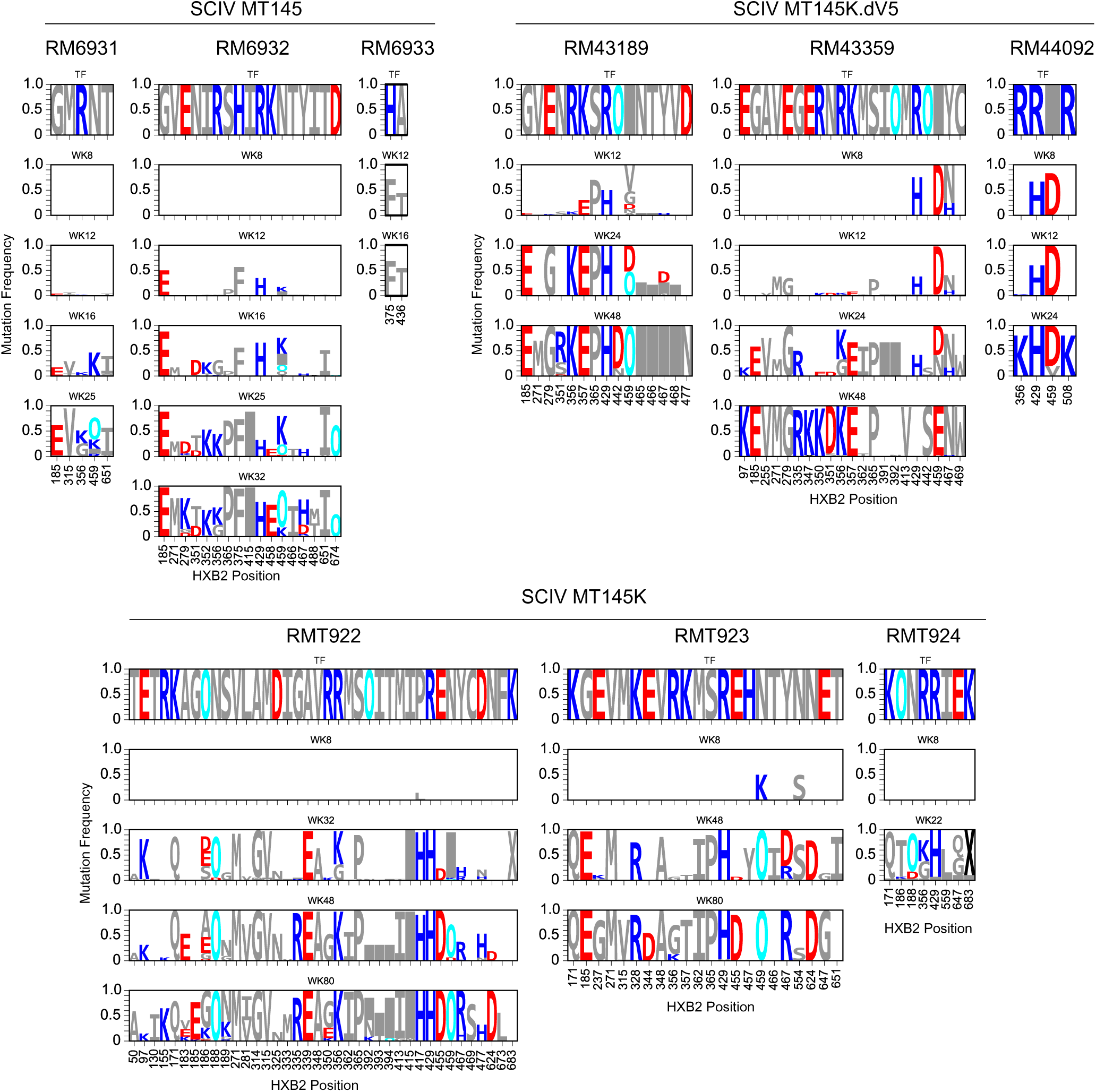
Longitudinal Antigenic Sequences and Sites from Intra-Host Evolution (LASSIE) selected sites for each RM. LASSIE using the TF loss criterion of 50% or more was applied to SGS data from each RM. Logo plots are shown for identified sites per RM, with the transmitter/founder (TF) amino acid sequence shown in the first row. O=glycan. Colors: O=cyan, DE=red, HRK=blue.

**Supplementary Figure 5.**
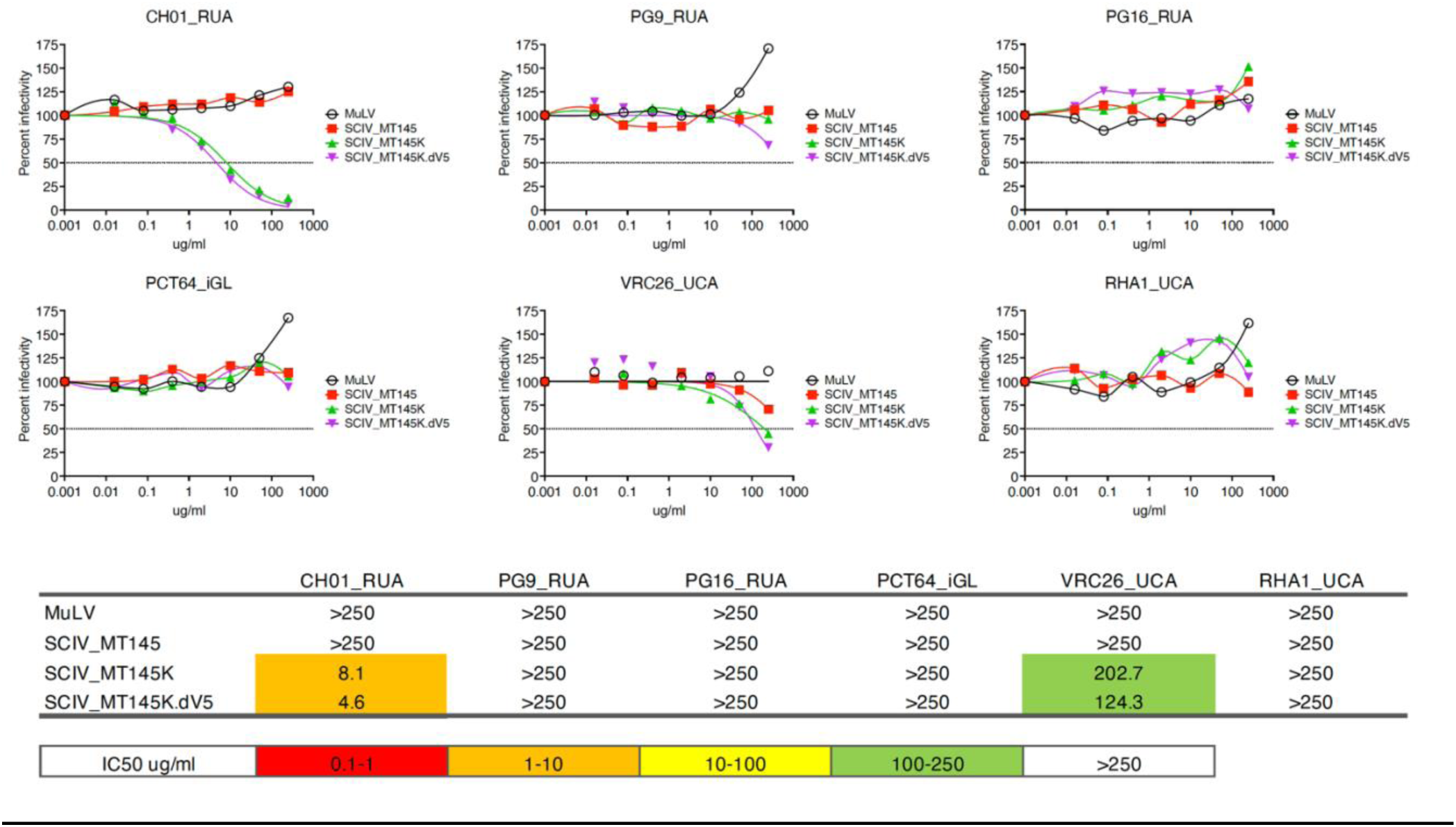
Germline-binding potential of SCIV_MT145, SCIV_MT145K and SCIV_MT145K.dV5. Neutralization curves depicting the sensitivity of SCIV_MT145 (red squares), SCIV_MT145K (green triangles), SCIV_MT145K.dV5 (purple triangles) and MLV (open circles, for control) to the RUA or iGL of the human V2-apex bnAbs CH01^55^, PG9^21,74^, PG16^74^, PCT64^56^, and VRC26^13^ and the rhesus V2-apex bnAb RHA1^53^. Dashed lines indicate 50% reduction in virus infectivity (the antibody concentration is shown on the x-axis in mg/ml). RUA, reverted unmutated ancestor; iGL, inferred germline; UCA, unmutated common ancestor.

**Supplementary Figure 6.**
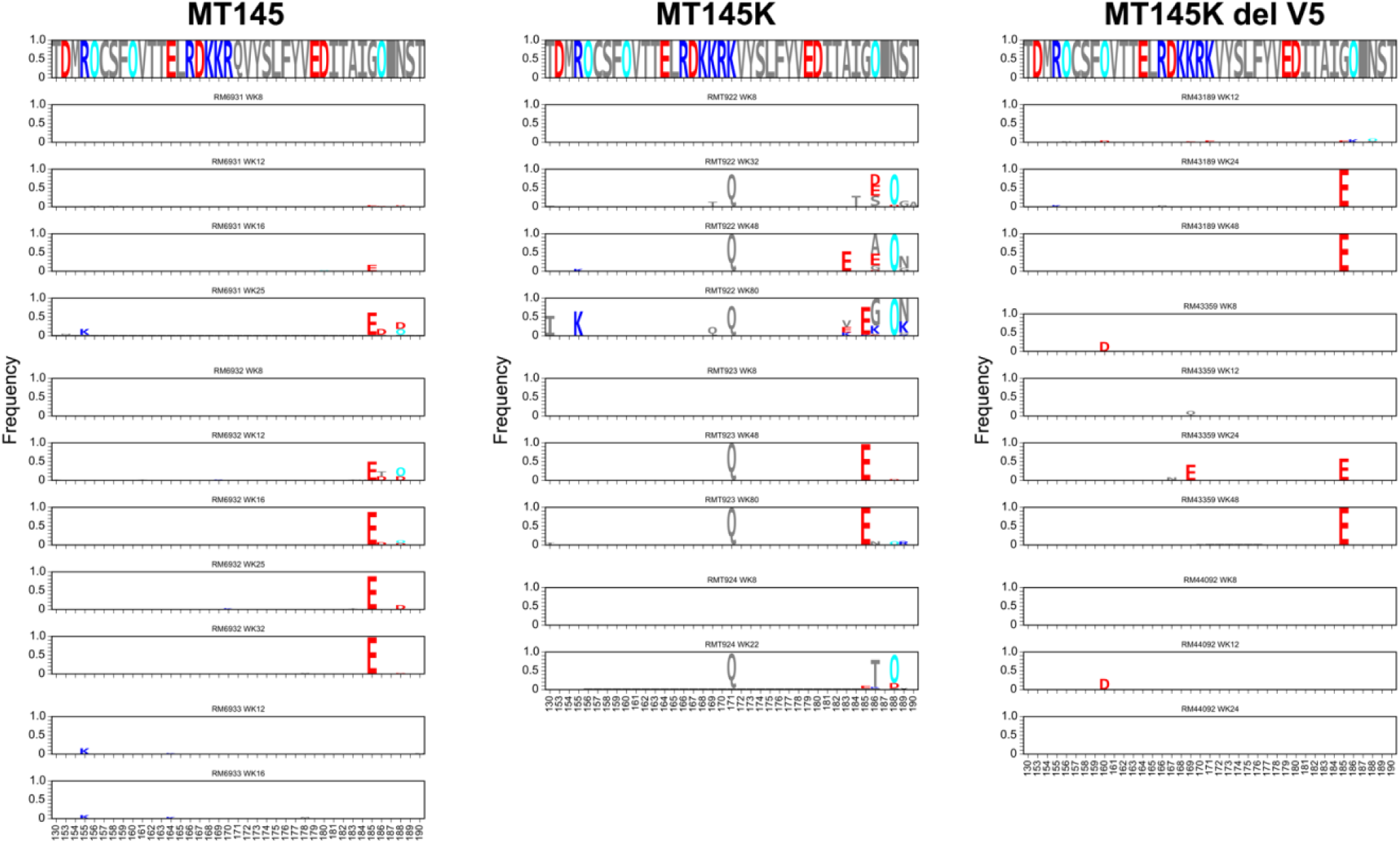
Env V2 region evolution for MT145 (left), MT145K (center), and MT145K.dV5 (right) groups. Sequence logos show the per time-point frequency of Env mutations away from the TF sequence using the single genome amplification sequences (SGA) from each RM. The horizontal axis labels are HXB2 numbers of sites. O=glycan. Colors: O=cyan, D,E=red, H,R,K=blue.

**Supplementary Figure 7.**
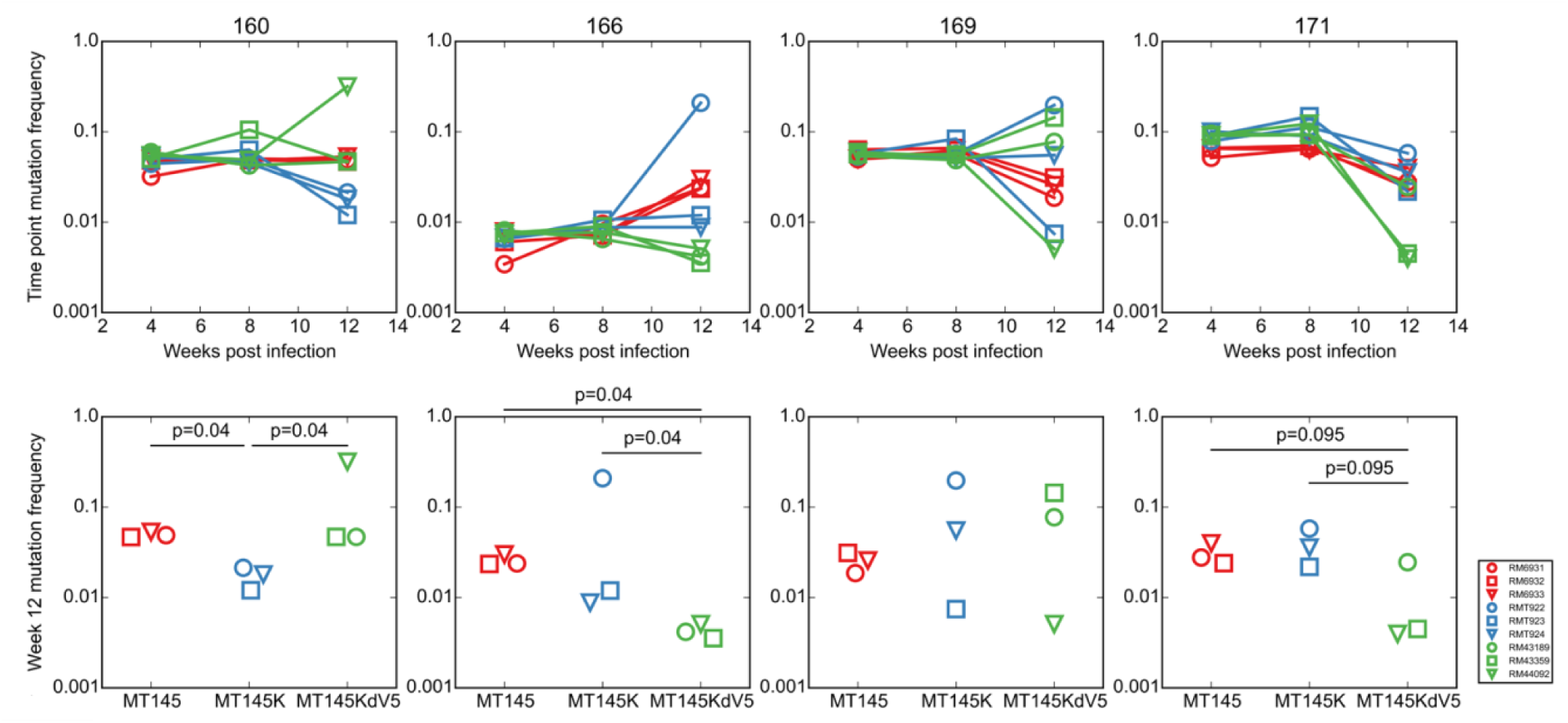
Mutation frequency at key V2 apex sites using NGS sequencing data. Top row: time course of mutation frequency for each RM at each site. Bottom row: comparison of mutation frequency at each site at week 8 across RMs. Each color is based on the SCIV group (red=MT145, blue=MT145K, and green=MT145K.dV5) and each line representing an individual animal is shown with a unique symbol. Significance of difference in frequency in bottom row calculated using Wilcoxon rank sum test, with two-sided uncorrected p < 0.05 shown.

**Supplementary Figure 8.**
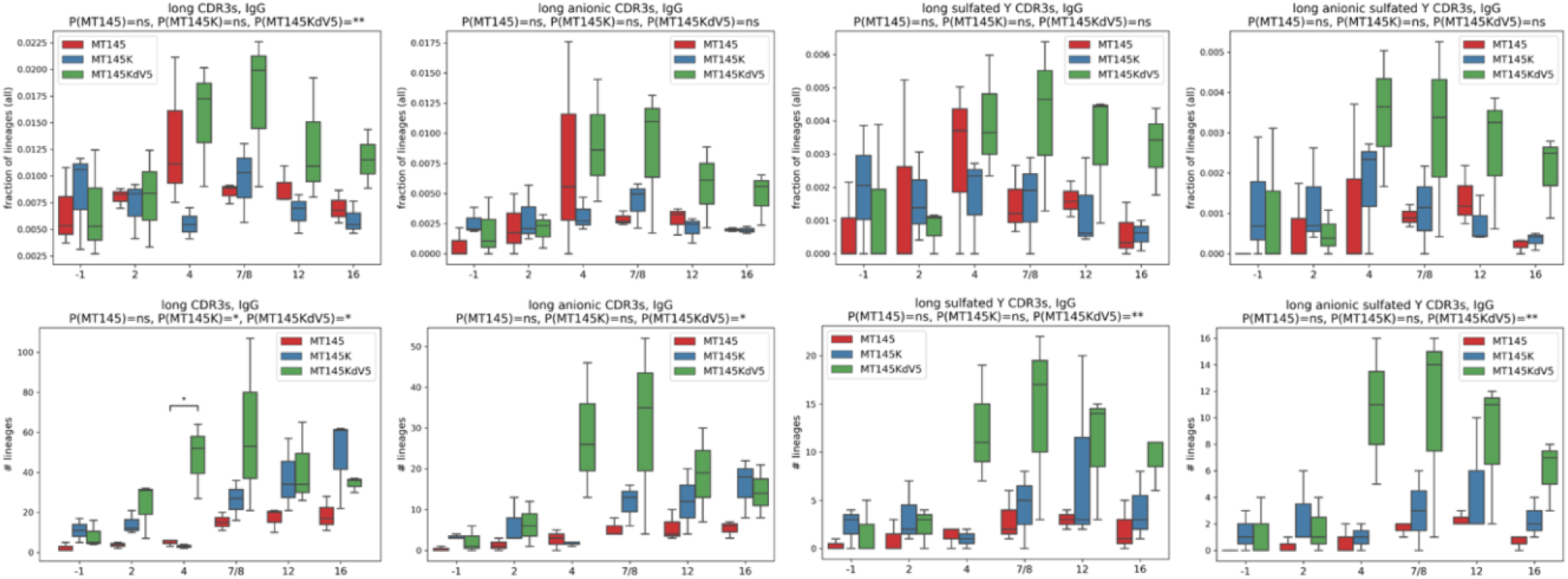
Longitudinal analysis of long HCDR3 lineages across MT145, MT145K, and MT145K.dV5 groups. The fraction and number of lineages containing long HCDR3, long anionic HCDR3, long sY HCDR3, and long anionic sY HCDR3s are shown in the top and bottom rows, respectively. P-values showing pairwise differences between percentages of lineages with long CDR3s across all time points. P-values were computed using the linear mixed model. (*P < 0.05; **P < 0.01)

**Supplementary Figure 9.**
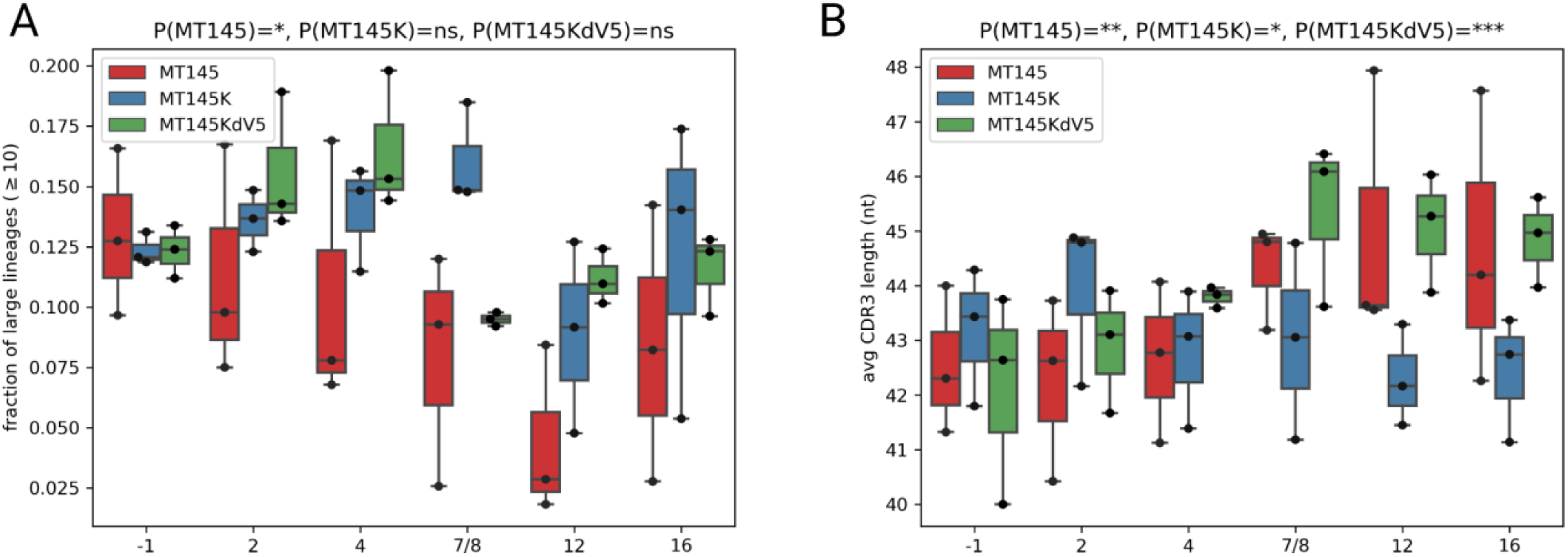
Longitudinal analysis of percentage of expanded lineages derived from IGHD3-9 (A) and HCDR3 length (B) across groups. P-values showing changes across time points were computed using the linear mixed model. (*P < 0.05;**P < 0.01;***P < 0.001).

**Supplementary Figure 10.**
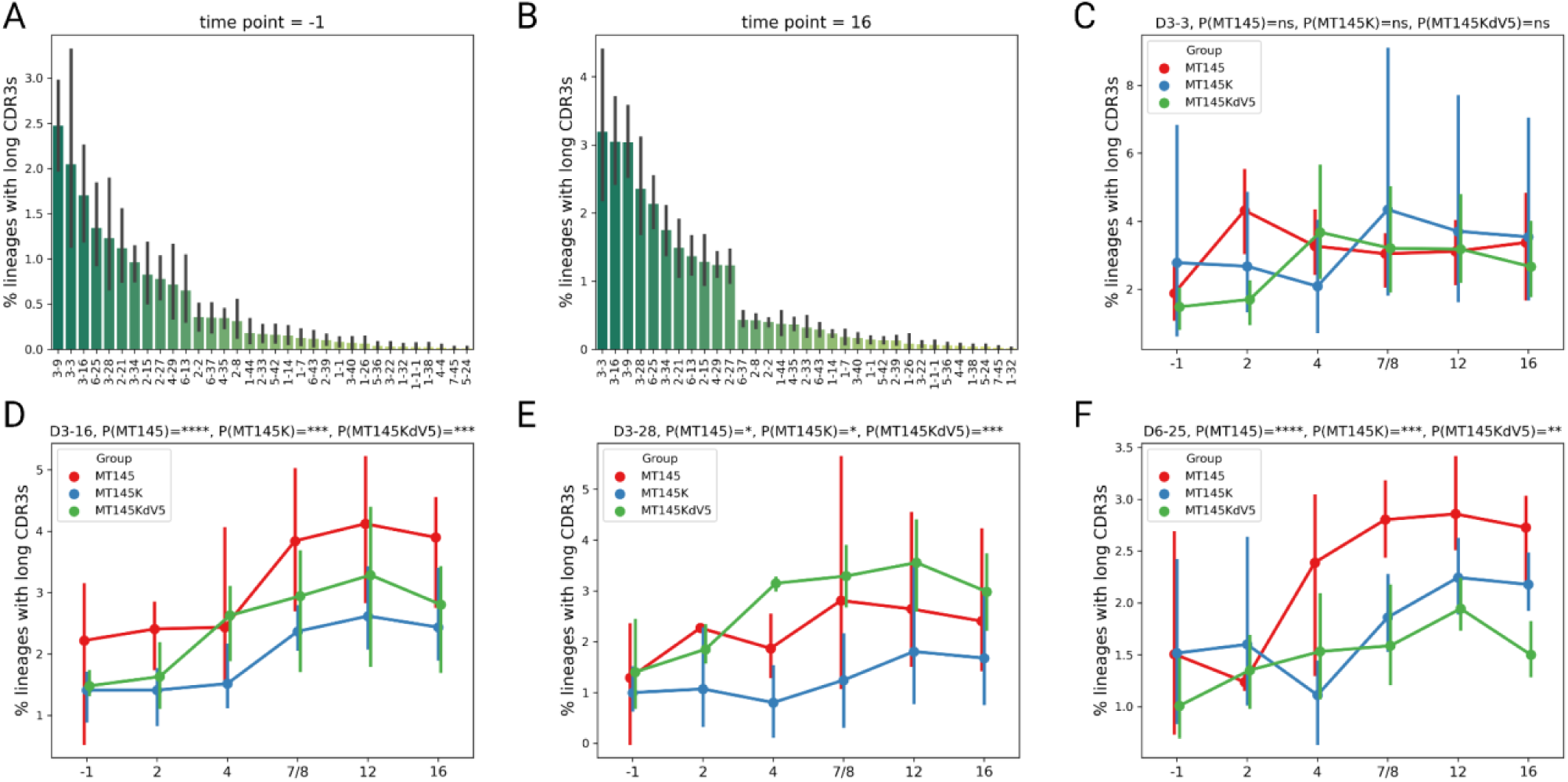
Repertoire analysis of longitudinal IGHD gene usages across groups. Percent of lineages with long HCDR3s containing specific IGHD genes for all animals in MT145, MT145K, and MT145K.dV5 groups (A) prior to infection and (B) 16 wpi. (C) Percent of lineages in long HCDR3s across groups for all time points containing (C) IGHD3-3, (D) IGHD3-16, (E) IGHD3-28, and (F) IGHD6-25. P-values showing pairwise differences between percentages of lineages with long CDR3s across all time points. P-values were computed using the lineage mixed model. (*P < 0.05; **P < 0.01;***P < 0.001;****P < 0.0001).

**Supplementary Figure 11.**
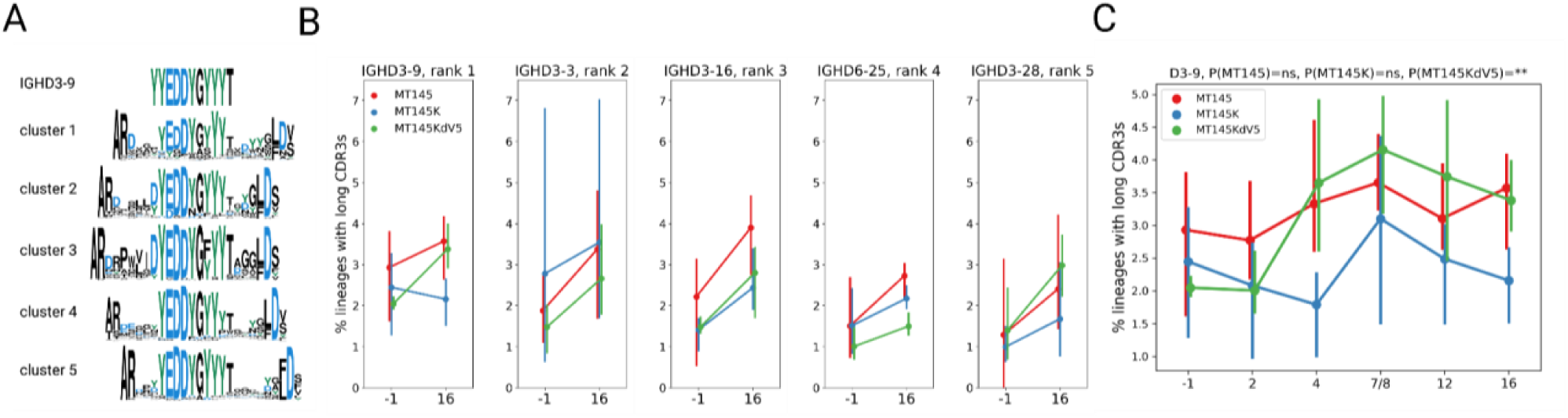
Enrichment of ‘EDDY’ motif encoding IGHD3-9 in MT145K.dV5 group animals (A) HCDR3s that use the IGHD3-09 gene that are clustered based on common amino acid motifs are reparented as sequence logo plots. The EDDY is conserved in the top 5 clusters identified from the repertoire of SCIV_MT145K.dV5 infected rhesus macaques. (B) Change in the percent of lineages with long HCDR3s for the top 5 D genes identified. (C) Change in the percent of long HCDR3s that use the D3-09 gene over all time points. P-values showing pairwise differences between percentages of lineages with long CDR3s across all time points. P-values were computed using the linear mixed model. (**P < 0.01)

**Supplementary Figure 12.**
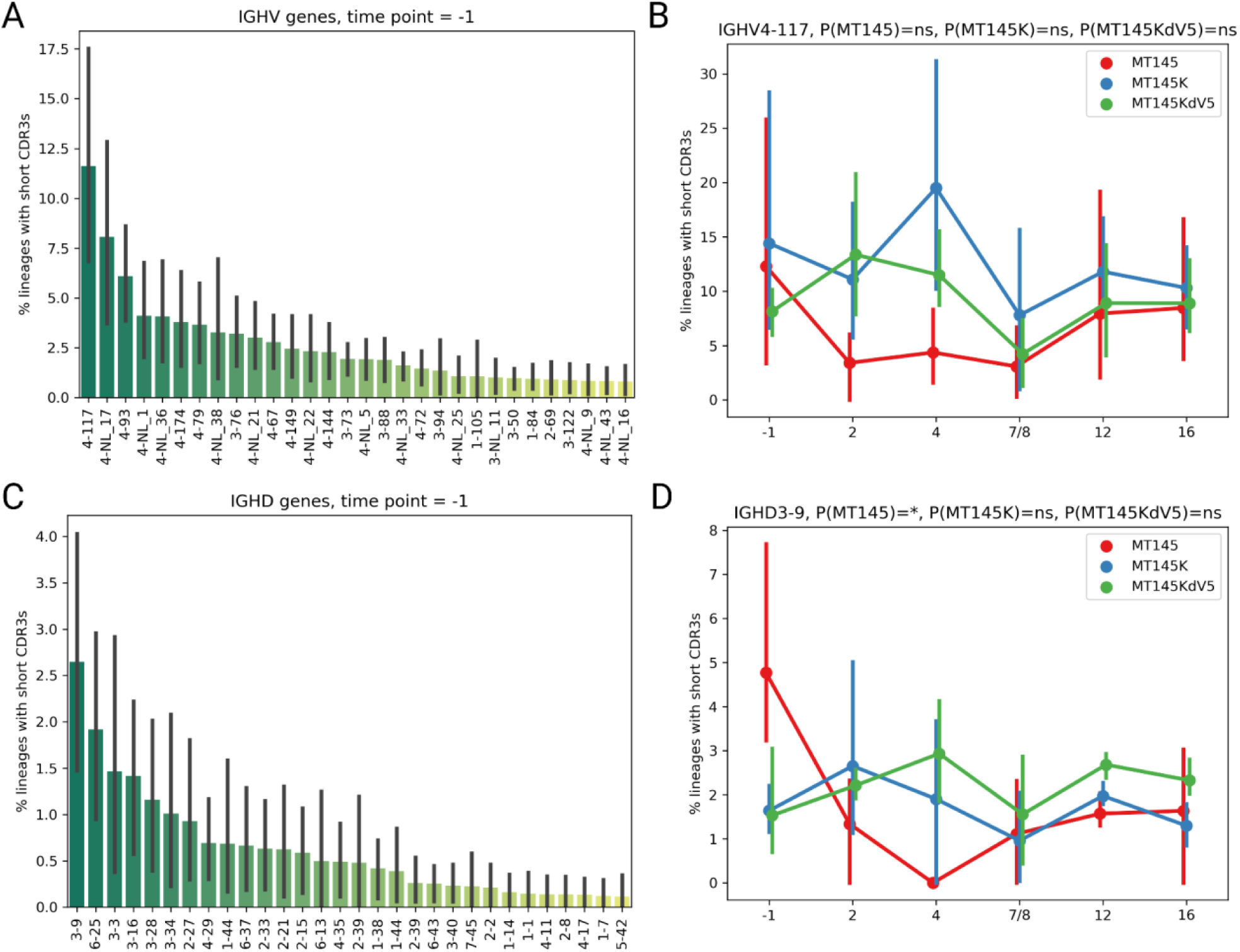
Frequencies of most prevalent V and D genes in short HCDR3 lineages present in pre-infection repertoires do not change after infection. (A) Percent of lineages with short HCDR3s that contain specific IGHV genes prior to infection. (B) Percent of short HCDR3 lineages with IGHV4-117 across groups over time. (C) Percent of lineages with short HCDR3s that contain specific IGHD genes prior to infection. (D) Percent of short HCDR3 lineages with IGHD3-9 across groups over time. P-values showing pairwise differences between percentages of lineages with long HCDR3s across all time points. P-values were computed using the linear mixed model. (*P < 0.05).

**Supplementary Figure 13.**
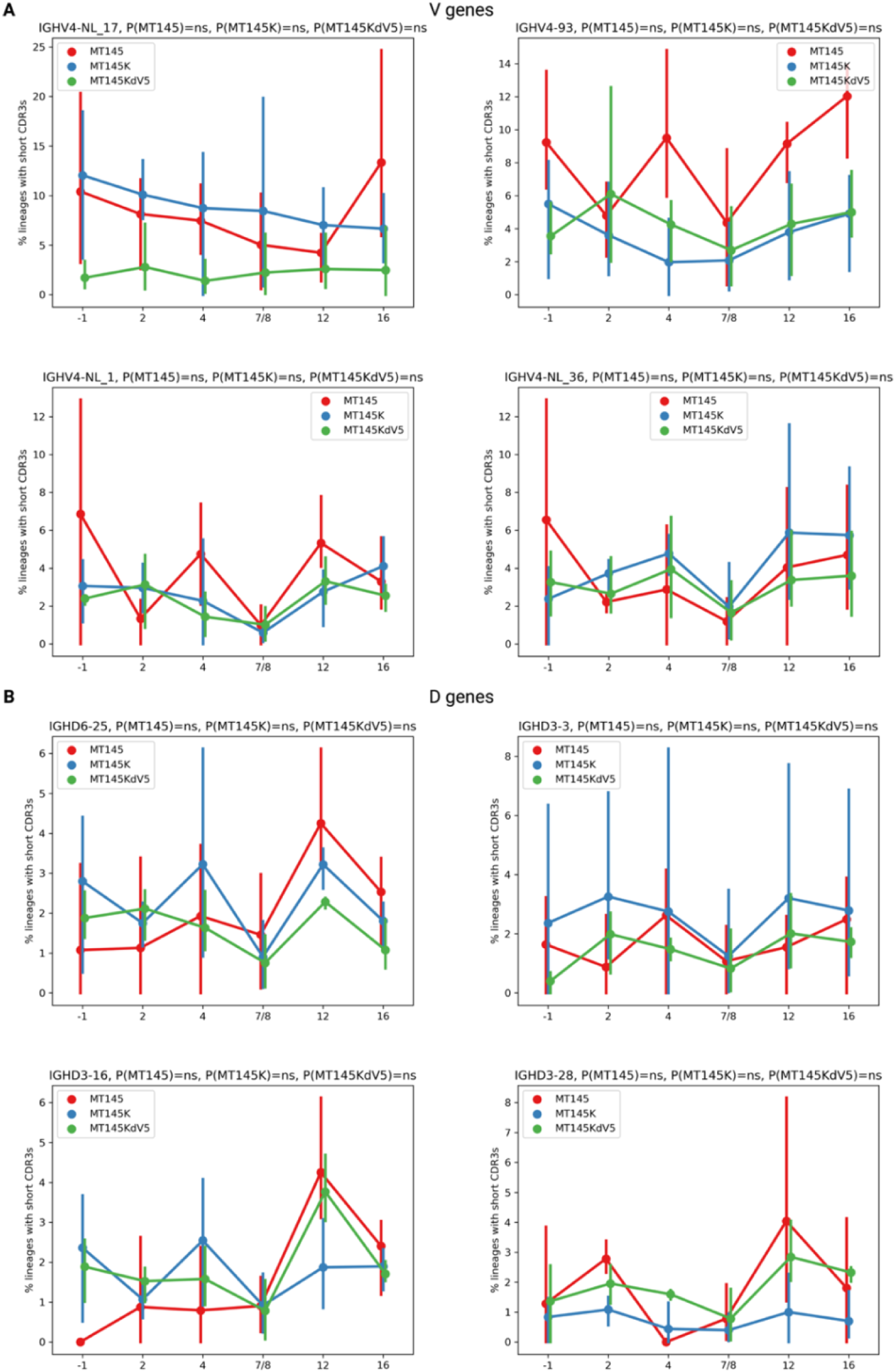
No significant difference in longitudinal V or D gene frequencies in short HCDR3s across groups. Percent of lineages with short HCDR3s containing common V genes (A) or D genes (B) prior to infection across groups over time. P-values were computed using the linear mixed model.

**Supplementary Figure 14.**
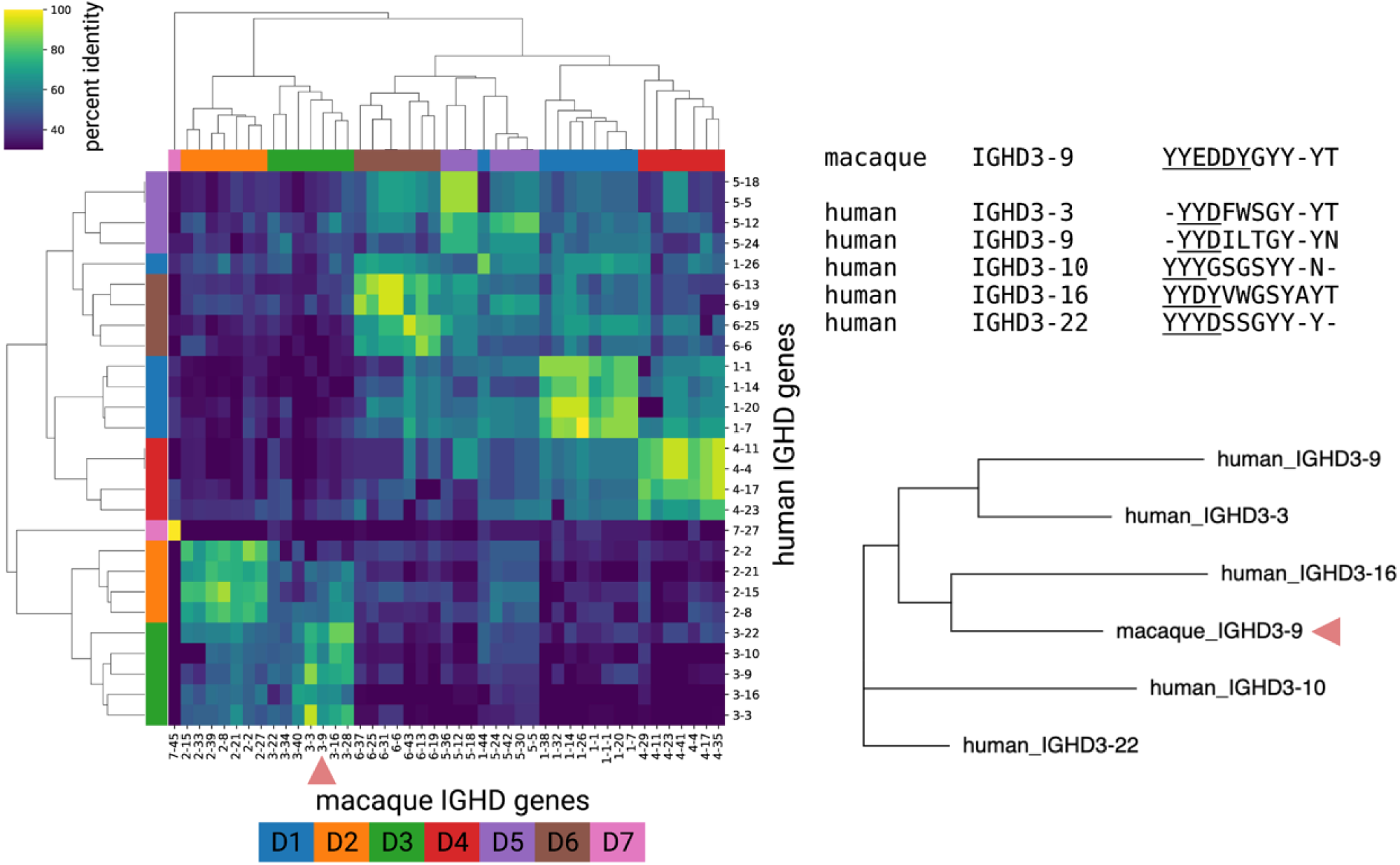
Germline IGHD gene similarities across humans and macaques. (Left) Phylogenetic heatmap of germline macaque IGHD genes on the horizontal axis against human germline IGHD genes on the vertical axis. Clusters of related genes are shown as a percent identifying color gradient. Red arrow indicates the RM IGHD3-09 gene. (Right) Human IGHD genes with similar anionic motifs characteristic of human and macaque V2 bnAbs.

**Supplementary Figure 15.**
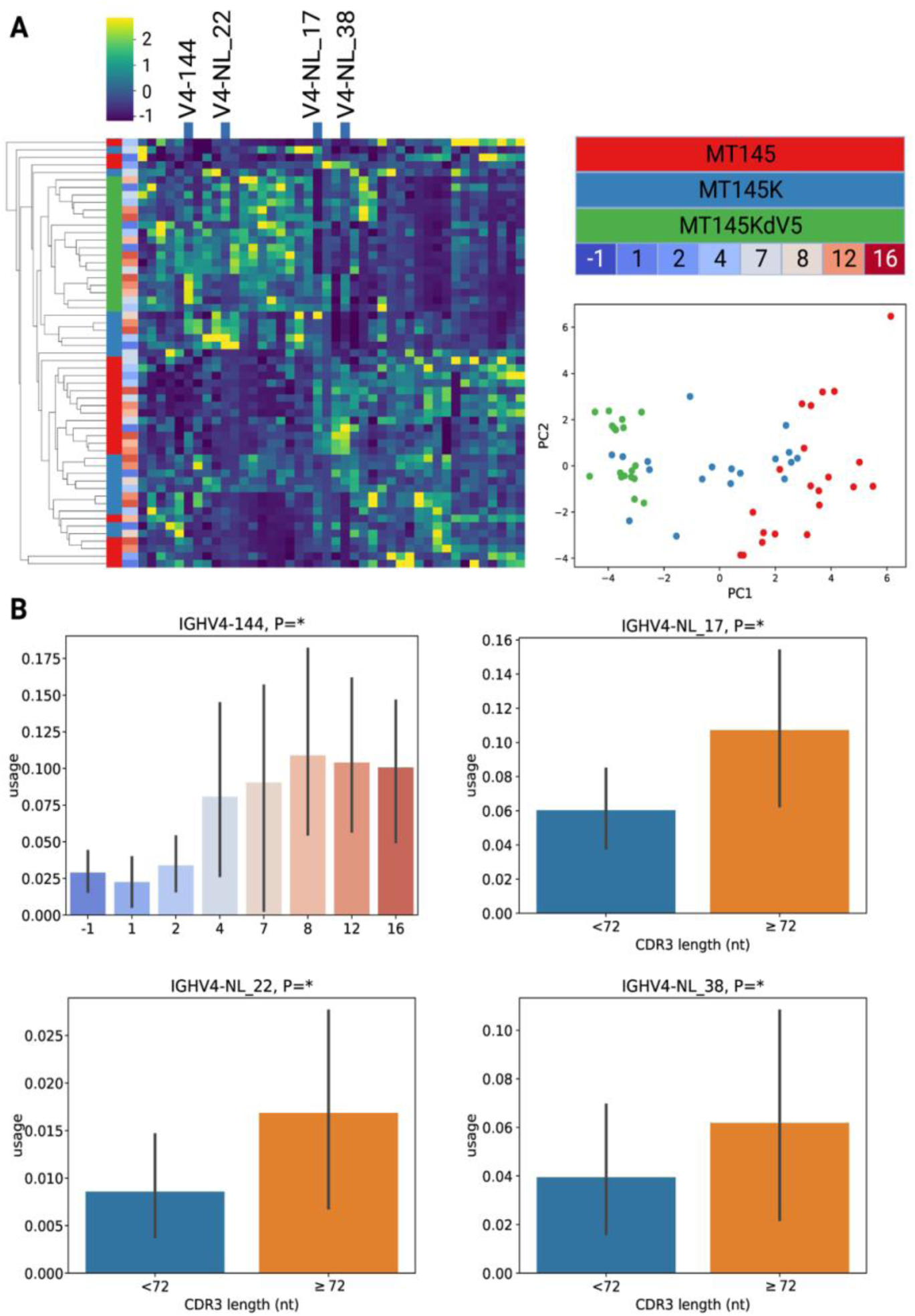
Longitudinal repertoire analysis of IGHV gene usages across groups reveals no enrichment of unique gene families or alleles. (A) A heatmap of germline macaque IGHV gene usages across animals and time points. Rows showing samples and columns showing IGHV genes of the heatmap are rearranged using hierarchical clustering. Four IGHV genes with statistically significant associations with usages across time usages or usages in short/long HCDR3s are shown on the top of the heatmap. The scatterplot on the right shows the principal component analysis of the usage matrix shown on the left. Principal components 1 and 2 are shown are x- and y-axes. Each point represents a sample and is colored across the group. (B) enrichment of IGHV4-144 gene across time and IGHV4-NL_17, IGHV4-NL_22, and IGHV4-NL_38 gene usage in short (<72 nt) and long (≥72 nt) HCDR3s in all animals. P-values showing pairwise differences between groups. P-values were computed using the linear mixed model and the Kruskal-Wallis test (*P < 0.05).

**Supplementary Figure 16.**
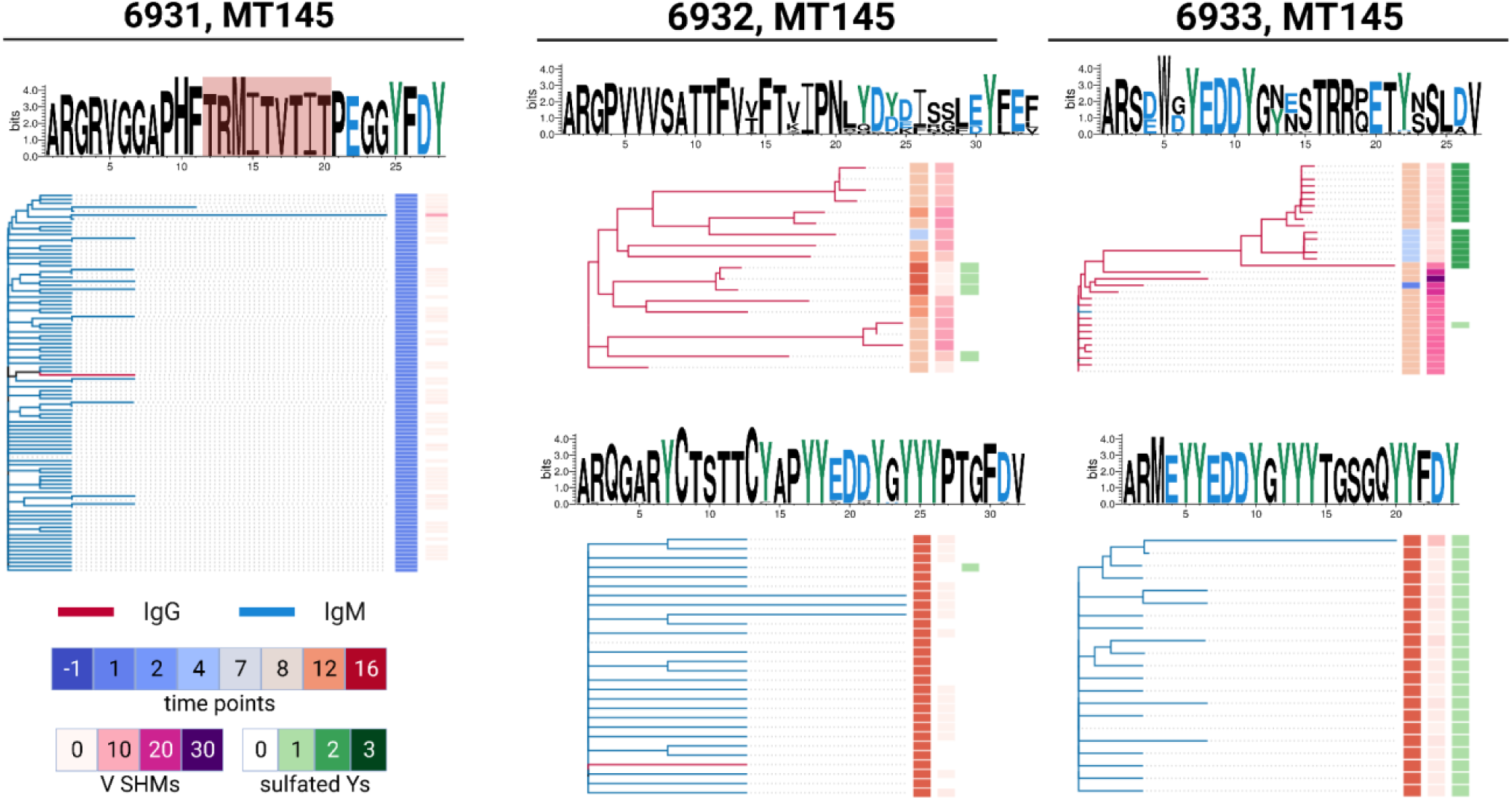
Top lineages from SCIV.MT145 infected animals containing long HCDR3s. Isotype, timepoint, SHM and Y sulfation levels are shown on the right columns of each tree. HCDR3 sequences are shown above each phylogenetic tree.

**Supplementary Figure 17.**
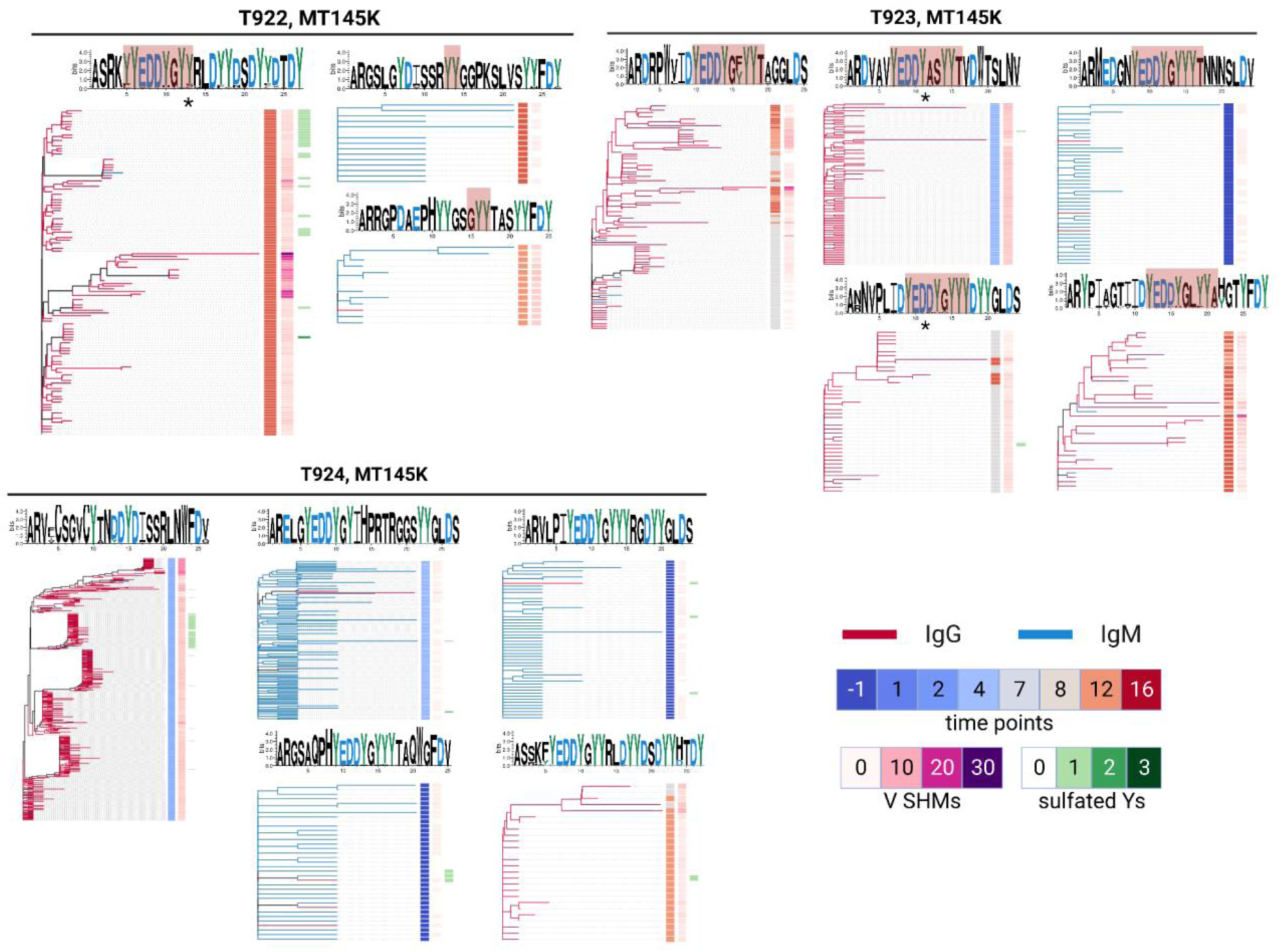
Top lineages from SCIV.MT145K infected animals containing long HCDR3s. Isotype, timepoint, SHM and Y sulfation levels are shown on the right columns of each tree. HCDR3 sequences are shown above each phylogenetic tree.

**Supplementary Figure 18.**
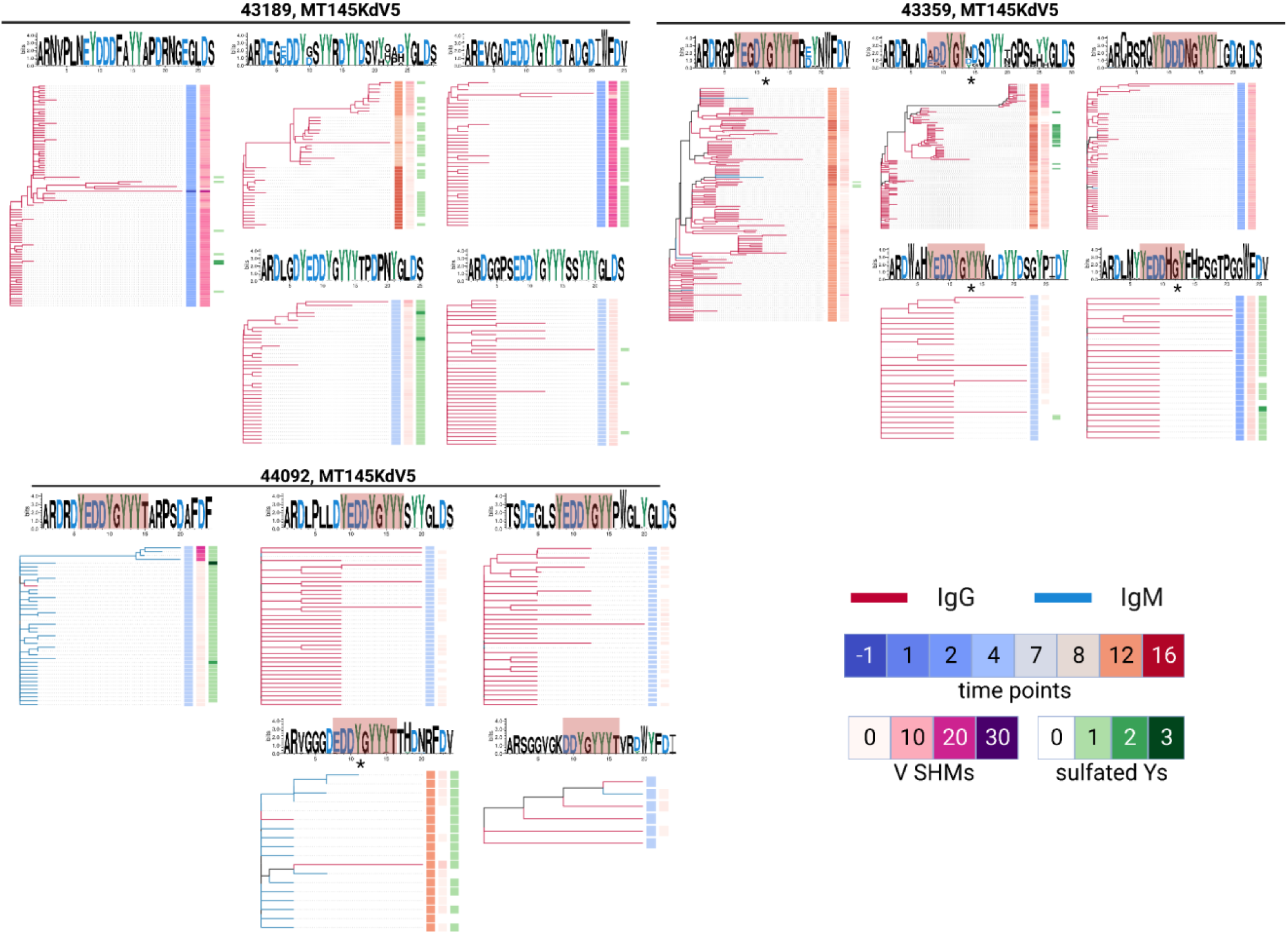
Top lineages from SCIV.MT145K.dV5 infected animals containing long HCDR3s. Isotype, timepoint, SHM and Y sulfation levels are shown on the right columns of each tree. HCDR3 sequences are shown above each phylogenetic tree.

**Supplementary Figure 19.**
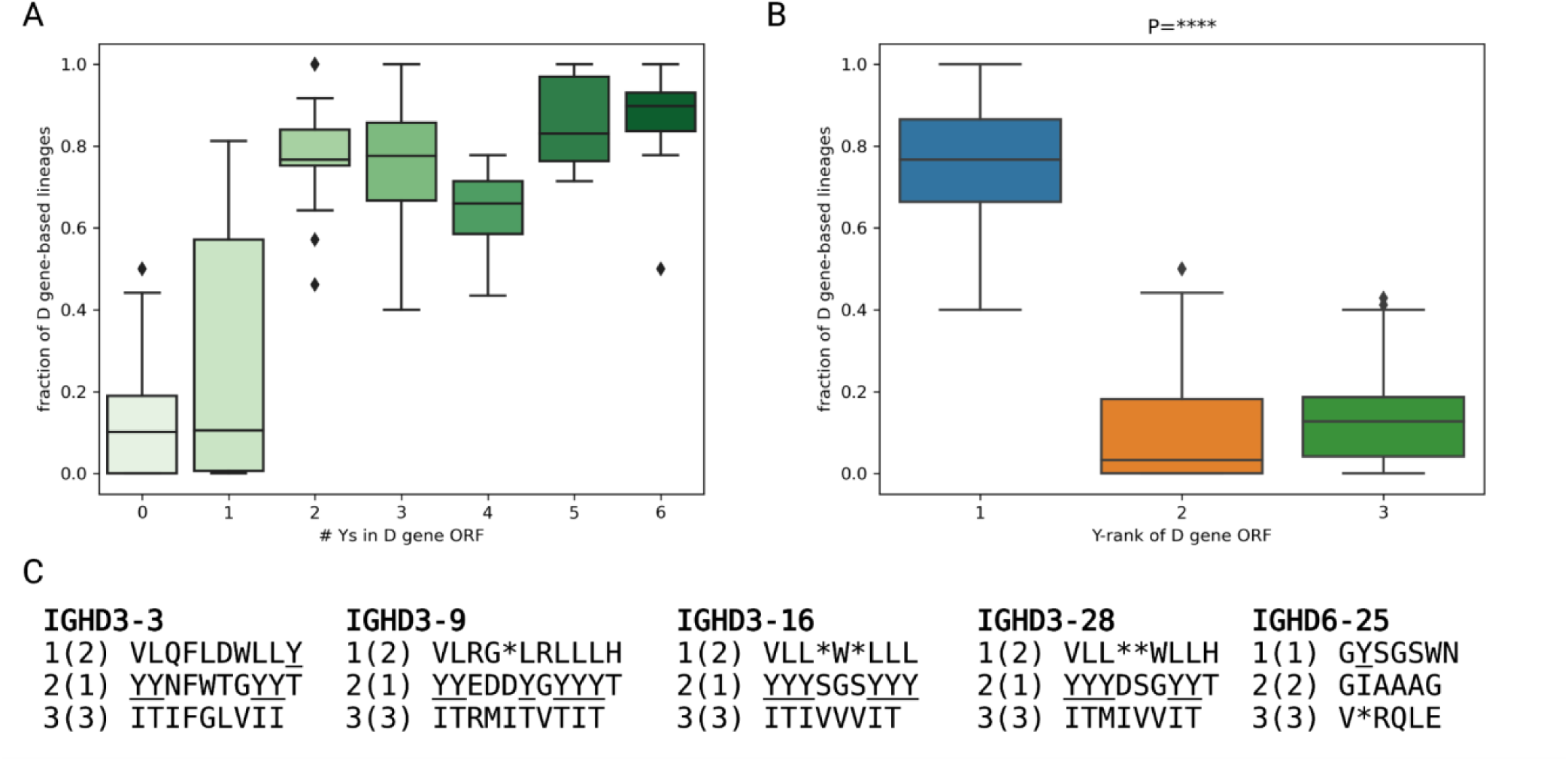
Antigen-specific repertoires are skewed for D gene ORFs enriched in tyrosine residues. (A) The fraction of lineages corresponding to D gene ORFs with 0, 1, …, 6 tyrosines. Each fraction was computed with respect to the number of lineages derived from a given D gene. (B) Three D gene ORFs were sorted in the descending order of the number of tyrosines in them, and ranks 1, 2, 3 were assigned to the ORFs with highest, medium and lowest numbers of tyrosines, respectively. The plot shows the fractions of lineages for D gene ORFs with ranks 1–3. P-value was computed using the Kruskal-Wallis test. (****P<0.0001) (C) ORFs of D genes with top five usages. The numbers before and in the parenthesis show the ORF number and the ORF rank, respectively.

**Supplementary Figure 20.**
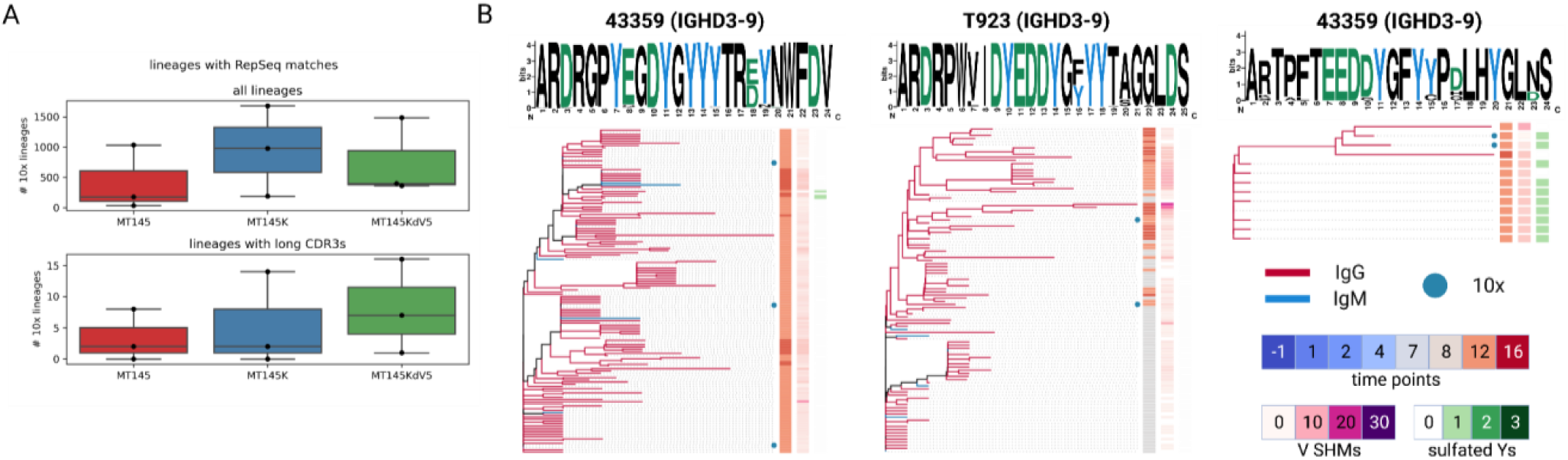
Antigen-specific IgG B cell lineages match features found in repertoire analysis. (A) Total number of lineages from antigen-sorted B cells that match sequences in the bulk repertoire for all (top) or long (bottom) HCDR3. (B) Phylogenetic trees for expanded lineages in SCIV_MT145K.dV5 and SCIV_MT145K animals with repertoire sequence matches. Isotype, timepoint, and SHM and Y sulfation levels are shown on the right columns of each tree. HCDR3 sequences are shown above each phylogenetic tree.

**Supplementary Figure 21.**
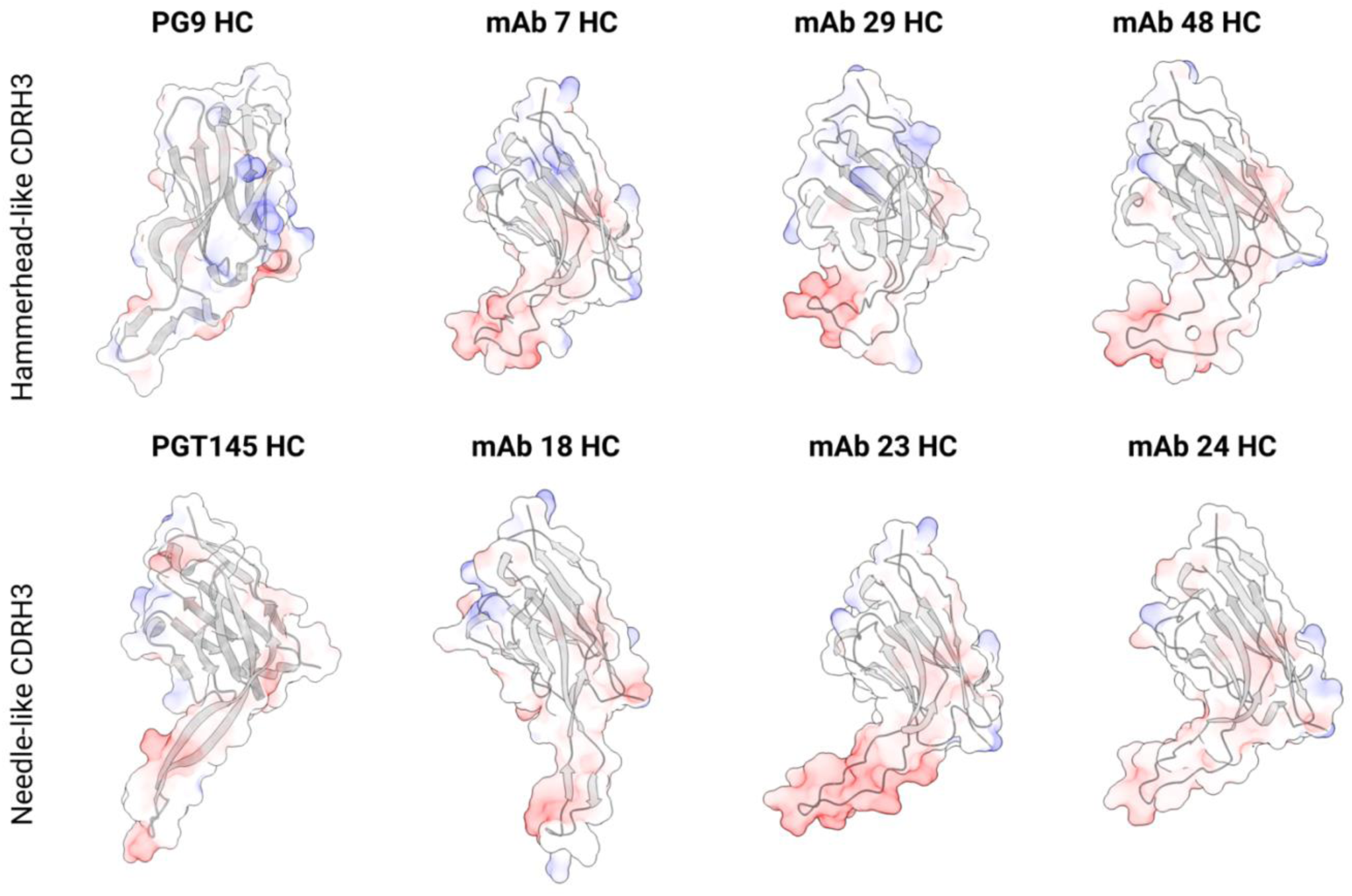
Structural prediction of antigen-specific mAb HCs. Antibodies with hammerhead-like (top) and needle-like (bottom) HCDR3s are shown. PG9 (3U2S) and PGT145 (3U1S) HC structures are shown on the left for comparison.

## References

1. Haynes, B.F., Wiehe, K., Borrow, P., Saunders, K.O., Korber, B., Wagh, K., McMichael, A.J., Kelsoe, G., Hahn, B.H., Alt, F., and Shaw, G.M. (2023). Strategies for HIV-1 vaccines that induce broadly neutralizing antibodies(vol 23, pg 142, 2023). Nature Reviews Immunology 23, 265–265. 10.1038/s41577-023-00854-0.

2. Burton, D.R., and Hangartner, L. (2016). Broadly Neutralizing Antibodies to HIV and Their Role in Vaccine Design. Annu Rev Immunol 34, 635–659. 10.1146/annurev-immunol-041015-055515.

3. 3. Derking, R., Ozorowski, G., Sliepen, K., Yasmeen, A., Cupo, A., Torres, J.L., Julien, J.P., Lee, J.H., van Montfort, T., de Taeye, S.W., et al. (2015). Comprehensive antigenic map of a cleaved soluble HIV-1 envelope trimer. PLoS Pathog 11, e1004767. 10.1371/journal.ppat.1004767.

4. Zhao, F., Joyce, C., Burns, A., Nogal, B., Cottrell, C.A., Ramos, A., Biddle, T., Pauthner, M., Nedellec, R., Qureshi, H., et al. (2020). Mapping Neutralizing Antibody Epitope Specificities to an HIV Env Trimer in Immunized and in Infected Rhesus Macaques. Cell Rep 32, 108122. 10.1016/j.celrep.2020.108122.

5. Zhou, T., Zheng, A., Baxa, U., Chuang, G.Y., Georgiev, I.S., Kong, R., O’Dell, S., Shahzad-Ul-Hussan, S., Shen, C.H., Tsybovsky, Y., et al. (2018). A Neutralizing Antibody Recognizing Primarily N-Linked Glycan Targets the Silent Face of the HIV Envelope. Immunity 48, 500–513 e506. 10.1016/j.immuni.2018.02.013.

6. Hraber, P., Seaman, M.S., Bailer, R.T., Mascola, J.R., Montefiori, D.C., and Korber, B.T. (2014). Prevalence of broadly neutralizing antibody responses during chronic HIV-1 infection. Aids 28, 163–169. 10.1097/QAD.0000000000000106.

7. Gray, E.S., Madiga, M.C., Hermanus, T., Moore, P.L., Wibmer, C.K., Tumba, N.L., Werner, L., Mlisana, K., Sibeko, S., Williamson, C., et al. (2011). The neutralization breadth of HIV-1 develops incrementally over four years and is associated with CD4+ T cell decline and high viral load during acute infection. J Virol 85, 4828–4840. 10.1128/JVI.00198-11.

8. Bonsignori, M., Liao, H.X., Gao, F., Williams, W.B., Alam, S.M., Montefiori, D.C., and Haynes, B.F. (2017). Antibody-virus co-evolution in HIV infection: paths for HIV vaccine development. Immunol Rev 275, 145–160. 10.1111/imr.12509.

9. Liao, H.X., Lynch, R., Zhou, T., Gao, F., Alam, S.M., Boyd, S.D., Fire, A.Z., Roskin, K.M., Schramm, C.A., Zhang, Z., et al. (2013). Co-evolution of a broadly neutralizing HIV-1 antibody and founder virus. Nature 496, 469–476. 10.1038/nature12053.

10. Burton, D.R., and Mascola, J.R. (2015). Antibody responses to envelope glycoproteins in HIV-1 infection. Nature immunology 16, 571–576. 10.1038/ni.3158.

11. Landais, E., Huang, X., Havenar-Daughton, C., Murrell, B., Price, M.A., Wickramasinghe, L., Ramos, A., Bian, C.B., Simek, M., Allen, S., et al. (2016). Broadly Neutralizing Antibody Responses in a Large Longitudinal Sub-Saharan HIV Primary Infection Cohort. PLoS pathogens 12, e1005369. 10.1371/journal.ppat.1005369.

12. Liao, H.X., Lynch, R., Zhou, T., Gao, F., Alam, S.M., Boyd, S.D., Fire, A.Z., Roskin, K.M., Schramm, C.A., Zhang, Z., et al. (2013). Co-evolution of a broadly neutralizing HIV-1 antibody and founder virus. Nature 496, 469–476. 10.1038/nature12053.

13. Doria-Rose, N.A., Schramm, C.A., Gorman, J., Moore, P.L., Bhiman, J.N., DeKosky, B.J., Ernandes, M.J., Georgiev, I.S., Kim, H.J., Pancera, M., et al. (2014). Developmental pathway for potent V1V2-directed HIV-neutralizing antibodies. Nature 509, 55–62. 10.1038/nature13036.

14. MacLeod, D.T., Choi, N.M., Briney, B., Garces, F., Ver, L.S., Landais, E., Murrell, B., Wrin, T., Kilembe, W., Liang, C.H., et al. (2016). Early Antibody Lineage Diversification and Independent Limb Maturation Lead to Broad HIV-1 Neutralization Targeting the Env High-Mannose Patch. Immunity 44, 1215–1226. 10.1016/j.immuni.2016.04.016.

15. Leggat, D.J., Cohen, K.W., Willis, J.R., Fulp, W.J., deCamp, A.C., Kalyuzhniy, O., Cottrell, C.A., Menis, S., Finak, G., Ballweber-Fleming, L., et al. (2022). Vaccination induces HIV broadly neutralizing antibody precursors in humans. Science 378, eadd6502. 10.1126/science.add6502.

16. Wu, X., Yang, Z.Y., Li, Y., Hogerkorp, C.M., Schief, W.R., Seaman, M.S., Zhou, T., Schmidt, S.D., Wu, L., Xu, L., et al. (2010). Rational design of envelope identifies broadly neutralizing human monoclonal antibodies to HIV-1. Science 329, 856–861. 10.1126/science.1187659.

17. Jardine, J.G., Kulp, D.W., Havenar-Daughton, C., Sarkar, A., Briney, B., Sok, D., Sesterhenn, F., Ereno-Orbea, J., Kalyuzhniy, O., Deresa, I., et al. (2016). HIV-1 broadly neutralizing antibody precursor B cells revealed by germline-targeting immunogene. Science 351, 1458–1463. 10.1126/science.aad9195.

18. Sok, D., Briney, B., Jardine, J.G., Kulp, D.W., Menis, S., Pauthner, M., Wood, A., Lee, E.C., Le, K.M., Jones, M., et al. (2016). Priming HIV-1 broadly neutralizing antibody precursors in human Ig loci transgenic mice. Science 353, 1557–1560. 10.1126/science.aah3945.

19. Abbott, R.K., Lee, J.H., Menis, S., Skog, P., Rossi, M., Ota, T., Kulp, D.W., Bhullar, D., Kalyuzhniy, O., Havenar-Daughton, C., et al. (2018). Precursor Frequency and Affinity Determine B Cell Competitive Fitness in Germinal Centers, Tested with Germline-Targeting HIV Vaccine Immunogens. Immunity 48, 133-+. 10.1016/j.immuni.2017.11.023.

20. Andrabi, R., Voss, J.E., Liang, C.H., Briney, B., Mccoy, L.E., Wu, C.Y., Wong, C.H., Poignard, P., and Burton, D.R. (2015). Identification of Common Features in Prototype Broadly Neutralizing Antibodies to HIV Envelope V2 Apex to Facilitate Vaccine Design. Immunity 43, 959–973. 10.1016/j.immuni.2015.10.014.

21. Willis, J.R., Berndsen, Z.T., Ma, K.M., Steichen, J.M., Schiffner, T., Landais, E., Liguori, A., Kalyuzhniy, O., Allen, J.D., Baboo, S., et al. (2022). Human immunoglobulin repertoire analysis guides design of vaccine priming immunogens targeting HIV V2-apex broadly neutralizing antibody precursors. Immunity 55, 2149–2167 e2149. 10.1016/j.immuni.2022.09.001.

22. Rantalainen, K., Berndsen, Z.T., Murrell, S., Cao, L., Omorodion, O., Torres, J.L., Wu, M., Umotoy, J., Copps, J., Poignard, P., et al. (2018). Co-evolution of HIV Envelope and Apex-Targeting Neutralizing Antibody Lineage Provides Benchmarks for Vaccine Design. Cell reports 23, 3249–3261. 10.1016/j.celrep.2018.05.046.

23. Briney, B.S., Willis, J.R., and Crowe, J.E., Jr. (2012). Human peripheral blood antibodies with long HCDR3s are established primarily at original recombination using a limited subset of germline genes. PloS one 7, e36750. 10.1371/journal.pone.0036750.

24. Pinto, D., Fenwick, C., Caillat, C., Silacci, C., Guseva, S., Dehez, F., Chipot, C., Barbieri, S., Minola, A., Jarrossay, D., et al. (2019). Structural Basis for Broad HIV-1 Neutralization by the MPER-Specific Human Broadly Neutralizing Antibody LN01. Cell Host Microbe 26, 623–637 e628. 10.1016/j.chom.2019.09.016.

25. Williams, L.D., Ofek, G., Schatzle, S., McDaniel, J.R., Lu, X., Nicely, N.I., Wu, L., Lougheed, C.S., Bradley, T., Louder, M.K., et al. (2017). Potent and broad HIV-neutralizing antibodies in memory B cells and plasma. Sci Immunol 2. 10.1126/sciimmunol.aal2200.

26. Steichen, J.M., Lin, Y.C., Havenar-Daughton, C., Pecetta, S., Ozorowski, G., Willis, J.R., Toy, L., Sok, D., Liguori, A., Kratochvil, S., et al. (2019). A generalized HIV vaccine design strategy for priming of broadly neutralizing antibody responses. Science 366. 10.1126/science.aax4380.

27. Steichen, J.M., Kulp, D.W., Tokatlian, T., Escolano, A., Dosenovic, P., Stanfield, R.L., McCoy, L.E., Ozorowski, G., Hu, X.Z., Kalyuzhniy, O., et al. (2016). HIV Vaccine Design to Target Germline Precursors of Glycan-Dependent Broadly Neutralizing Antibodies. Immunity 45, 483–496. 10.1016/j.immuni.2016.08.016.

28. Caniels, T.G., Medina-Ramirez, M., Zhang, J., Sarkar, A., Kumar, S., LaBranche, A., Derking, R., Allen, J.D., Snitselaar, J.L., Capella-Pujol, J., et al. (2023). Germline-targeting HIV-1 Env vaccination induces VRC01-class antibodies with rare insertions. Cell Rep Med 4, 101003. 10.1016/j.xcrm.2023.101003.

29. Voss, J.E., Andrabi, R., McCoy, L.E., de Val, N., Fuller, R.P., Messmer, T., Su, C.Y., Sok, D., Khan, S.N., Garces, F., et al. (2017). Elicitation of Neutralizing Antibodies Targeting the V2 Apex of the HIV Envelope Trimer in a Wild-Type Animal Model. Cell reports 21, 222–235. 10.1016/j.celrep.2017.09.024.

30. Andrabi, R., Voss, J.E., Liang, C.H., Briney, B., McCoy, L.E., Wu, C.Y., Wong, C.H., Poignard, P., and Burton, D.R. (2015). Identification of Common Features in Prototype Broadly Neutralizing Antibodies to HIV Envelope V2 Apex to Facilitate Vaccine Design. Immunity 43, 959–973. 10.1016/j.immuni.2015.10.014.

31. McGuire, A.T., Hoot, S., Dreyer, A.M., Lippy, A., Stuart, A., Cohen, K.W., Jardine, J., Menis, S., Scheid, J.F., West, A.P., et al. (2013). Engineering HIV envelope protein to activate germline B cell receptors of broadly neutralizing anti-CD4 binding site antibodies. The Journal of experimental medicine 210, 655–663. 10.1084/jem.20122824.

32. Jardine, J.G., Ota, T., Sok, D., Pauthner, M., Kulp, D.W., Kalyuzhniy, O., Skog, P.D., Thinnes, T.C., Bhullar, D., Briney, B., et al. (2015). HIV-1 VACCINES. Priming a broadly neutralizing antibody response to HIV-1 using a germline-targeting immunogen. Science 349, 156–161. 10.1126/science.aac5894.

33. Escolano, A., Steichen, J.M., Dosenovic, P., Kulp, D.W., Golijanin, J., Sok, D., Freund, N.T., Gitlin, A.D., Oliveira, T., Araki, T., et al. (2016). Sequential Immunization Elicits Broadly Neutralizing Anti-HIV-1 Antibodies in Ig Knockin Mice. Cell 166, 1445–1458 e1412. 10.1016/j.cell.2016.07.030.

34. Briney, B., Sok, D., Jardine, J.G., Kulp, D.W., Skog, P., Menis, S., Jacak, R., Kalyuzhniy, O., de Val, N., Sesterhenn, F., et al. (2016). Tailored Immunogens Direct Affinity Maturation toward HIV Neutralizing Antibodies. Cell 166, 1459–1470 e1411. 10.1016/j.cell.2016.08.005.

35. Sok, D., Briney, B., Jardine, J.G., Kulp, D.W., Menis, S., Pauthner, M., Wood, A., Lee, E.C., Le, K.M., Jones, M., et al. (2016). Priming HIV-1 broadly neutralizing antibody precursors in human Ig loci transgenic mice. Science. 10.1126/science.aah3945.

36. Tian, M., Cheng, C., Chen, X., Duan, H., Cheng, H.L., Dao, M., Sheng, Z., Kimble, M., Wang, L., Lin, S., et al. (2016). Induction of HIV Neutralizing Antibody Lineages in Mice with Diverse Precursor Repertoires. Cell 166, 1471–1484 e1418. 10.1016/j.cell.2016.07.029.

37. Martina, C.E., Crowe, J.E., Jr., and Meiler, J. (2023). Glycan masking in vaccine design: Targets, immunogens and applications. Front Immunol 14, 1126034. 10.3389/fimmu.2023.1126034.

38. Crooks, E.T., Almanza, F., D’Addabbo, A., Duggan, E., Zhang, J., Wagh, K., Mou, H., Allen, J.D., Thomas, A., Osawa, K., et al. (2021). Engineering well-expressed, V2-immunofocusing HIV-1 envelope glycoprotein membrane trimers for use in heterologous prime-boost vaccine regimens. PLoS Pathog 17, e1009807. 10.1371/journal.ppat.1009807.

39. Brouwer, P.J.M., Antanasijevic, A., de Gast, M., Allen, J.D., Bijl, T.P.L., Yasmeen, A., Ravichandran, R., Burger, J.A., Ozorowski, G., Torres, J.L., et al. (2021). Immunofocusing and enhancing autologous Tier-2 HIV-1 neutralization by displaying Env trimers on two-component protein nanoparticles. Npj Vaccines 6, 24. 10.1038/s41541-021-00285-9.

40. Bibollet-Ruche, F., Russell, R.M., Ding, W., Liu, W., Li, Y., Wagh, K., Wrapp, D., Habib, R., Skelly, A.N., Roark, R.S., et al. (2023). A Germline-Targeting Chimpanzee SIV Envelope Glycoprotein Elicits a New Class of V2-Apex Directed Cross-Neutralizing Antibodies. mBio 14, e0337022. 10.1128/mbio.03370-22.

41. Moyer, T.J., Kato, Y., Abraham, W., Chang, J.Y.H., Kulp, D.W., Watson, N., Turner, H.L., Menis, S., Abbott, R.K., Bhiman, J.N., et al. (2020). Engineered immunogen binding to alum adjuvant enhances humoral immunity. Nature medicine 26, 430–440. 10.1038/s41591-020-0753-3.

42. Kulp, D.W., Steichen, J.M., Pauthner, M., Hu, X., Schiffner, T., Liguori, A., Cottrell, C.A., Havenar-Daughton, C., Ozorowski, G., Georgeson, E., et al. (2017). Structure-based design of native-like HIV-1 envelope trimers to silence non-neutralizing epitopes and eliminate CD4 binding. Nature communications 8, 1655. 10.1038/s41467-017-01549-6.

43. Sanders, R.W., and Moore, J.P. (2017). Native-like Env trimers as a platform for HIV-1 vaccine design. Immunological Reviews 275, 161–182. 10.1111/imr.12481.

44. Ringe, R.P., Yasmeen, A., Ozorowski, G., Go, E.P., Pritchard, L.K., Guttman, M., Ketas, T.A., Cottrell, C.A., Wilson, I.A., Sanders, R.W., et al. (2015). Influences on the Design and Purification of Soluble, Recombinant Native-Like HIV-1 Envelope Glycoprotein Trimers. Journal of virology 89, 12189–12210. 10.1128/JVI.01768-15.

45. Kumar, R., Deshpande, S., Sewall, L.M., Ozorowski, G., Cottrell, C.A., Lee, W.H., Holden, L.G., Richey, S.T., Chandrawacar, A.S., Dhiman, K., et al. (2021). Elicitation of potent serum neutralizing antibody responses in rabbits by immunization with an HIV-1 clade C trimeric Env derived from an Indian elite neutralizer. PLoS Pathog 17, e1008977. 10.1371/journal.ppat.1008977.

46. Sanders, R.W., van Gils, M.J., Derking, R., Sok, D., Ketas, T.J., Burger, J.A., Ozorowski, G., Cupo, A., Simonich, C., Goo, L., et al. (2015). HIV-1 VACCINES. HIV-1 neutralizing antibodies induced by native-like envelope trimers. Science 349, aac4223. 10.1126/science.aac4223.

47. Sahoo, A., Hodge, E.A., LaBranche, C.C., Styles, T.M., Shen, X., Cheedarla, N., Shiferaw, A., Ozorowski, G., Lee, W.H., Ward, A.B., et al. (2022). Structure-guided changes at the V2 apex of HIV-1 clade C trimer enhance elicitation of autologous neutralizing and broad V1V2-scaffold antibodies. Cell Rep 38, 110436. 10.1016/j.celrep.2022.110436.

48. de Taeye, S.W., Go, E.P., Sliepen, K., de la Pena, A.T., Badal, K., Medina-Ramirez, M., Lee, W.H., Desaire, H., Wilson, I.A., Moore, J.P., et al. (2019). Stabilization of the V2 loop improves the presentation of V2 loop-associated broadly neutralizing antibody epitopes on HIV-1 envelope trimers. J Biol Chem 294, 5616–5631. 10.1074/jbc.RA118.005396.

49. Walker, L.M., Phogat, S.K., Chan-Hui, P.Y., Wagner, D., Phung, P., Goss, J.L., Wrin, T., Simek, M.D., Fling, S., Mitcham, J.L., et al. (2009). Broad and potent neutralizing antibodies from an African donor reveal a new HIV-1 vaccine target. Science 326, 285–289. 10.1126/science.1178746.

50. Andrabi, R., Pallesen, J., Allen, J.D., Song, G., Zhang, J., de Val, N., Gegg, G., Porter, K., Su, C.Y., Pauthner, M., et al. (2019). The Chimpanzee SIV Envelope Trimer: Structure and Deployment as an HIV Vaccine Template. Cell reports 27, 2426–2441 e2426. 10.1016/j.celrep.2019.04.082.

51. Gorman, J., Soto, C., Yang, M.M., Davenport, T.M., Guttman, M., Bailer, R.T., Chambers, M., Chuang, G.Y., DeKosky, B.J., Doria-Rose, N.A., et al. (2016). Structures of HIV-1 Env V1V2 with broadly neutralizing antibodies reveal commonalities that enable vaccine design. Nature structural & molecular biology 23, 81–90. 10.1038/nsmb.3144.

52. McLellan, J.S., Pancera, M., Carrico, C., Gorman, J., Julien, J.P., Khayat, R., Louder, R., Pejchal, R., Sastry, M., Dai, K., et al. (2011). Structure of HIV-1 gp120 V1/V2 domain with broadly neutralizing antibody PG9. Nature 480, 336–343. 10.1038/nature10696.

53. Roark, R.S., Li, H., Williams, W.B., Chug, H., Mason, R.D., Gorman, J., Wang, S., Lee, F.H., Rando, J., Bonsignori, M., et al. (2021). Recapitulation of HIV-1 Env-antibody coevolution in macaques leading to neutralization breadth. Science 371. 10.1126/science.abd2638.

54. Gorman, J., Chuang, G.Y., Lai, Y.T., Shen, C.H., Boyington, J.C., Druz, A., Geng, H., Louder, M.K., McKee, K., Rawi, R., et al. (2020). Structure of Super-Potent Antibody CAP256-VRC26.25 in Complex with HIV-1 Envelope Reveals a Combined Mode of Trimer-Apex Recognition. Cell reports 31, 107488. 10.1016/j.celrep.2020.03.052.

55. Bonsignori, M., Hwang, K.K., Chen, X., Tsao, C.Y., Morris, L., Gray, E., Marshall, D.J., Crump, J.A., Kapiga, S.H., Sam, N.E., et al. (2011). Analysis of a clonal lineage of HIV-1 envelope V2/V3 conformational epitope-specific broadly neutralizing antibodies and their inferred unmutated common ancestors. J Virol 85, 9998–10009. 10.1128/JVI.05045-11.

56. Landais, E., Murrell, B., Briney, B., Murrell, S., Rantalainen, K., Berndsen, Z.T., Ramos, A., Wickramasinghe, L., Smith, M.L., Eren, K., et al. (2017). HIV Envelope Glycoform Heterogeneity and Localized Diversity Govern the Initiation and Maturation of a V2 Apex Broadly Neutralizing Antibody Lineage. Immunity 47, 990–1003 e1009. 10.1016/j.immuni.2017.11.002.

57. Doria-Rose, N.A., Bhiman, J.N., Roark, R.S., Schramm, C.A., Gorman, J., Chuang, G.Y., Pancera, M., Cale, E.M., Ernandes, M.J., Louder, M.K., et al. (2016). New Member of the V1V2-Directed CAP256-VRC26 Lineage That Shows Increased Breadth and Exceptional Potency. J Virol 90, 76–91. 10.1128/JVI.01791-15.

58. Walker, L.M., Huber, M., Doores, K.J., Falkowska, E., Pejchal, R., Julien, J.P., Wang, S.K., Ramos, A., Chan-Hui, P.Y., Moyle, M., et al. (2011). Broad neutralization coverage of HIV by multiple highly potent antibodies. Nature 477, 466–470. 10.1038/nature10373.

59. Lee, J.H., Andrabi, R., Su, C.Y., Yasmeen, A., Julien, J.P., Kong, L., Wu, N.C., McBride, R., Sok, D., Pauthner, M., et al. (2017). A Broadly Neutralizing Antibody Targets the Dynamic HIV Envelope Trimer Apex via a Long, Rigidified, and Anionic beta-Hairpin Structure. Immunity 46, 690–702. 10.1016/j.immuni.2017.03.017.

60. Cai, C.X., Doria-Rose, N.A., Schneck, N.A., Ivleva, V.B., Tippett, B., Shadrick, W.R., O’Connell, S., Cooper, J.W., Schneiderman, Z., Zhang, B., et al. (2022). Tyrosine O-sulfation proteoforms affect HIV-1 monoclonal antibody potency. Sci Rep 12, 8433. 10.1038/s41598-022-12423-x.

61. Pejchal, R., Walker, L.M., Stanfield, R.L., Phogat, S.K., Koff, W.C., Poignard, P., Burton, D.R., and Wilson, I.A. (2010). Structure and function of broadly reactive antibody PG16 reveal an H3 subdomain that mediates potent neutralization of HIV-1. Proceedings of the National Academy of Sciences of the United States of America 107, 11483–11488. 10.1073/pnas.1004600107.

62. Gao, F., Bailes, E., Robertson, D.L., Chen, Y., Rodenburg, C.M., Michael, S.F., Cummins, L.B., Arthur, L.O., Peeters, M., Shaw, G.M., et al. (1999). Origin of HIV-1 in the chimpanzee Pan troglodytes troglodytes. Nature 397, 436–441. 10.1038/17130.

63. Keele, B.F., Van Heuverswyn, F., Li, Y., Bailes, E., Takehisa, J., Santiago, M.L., Bibollet-Ruche, F., Chen, Y., Wain, L.V., Liegeois, F., et al. (2006). Chimpanzee reservoirs of pandemic and nonpandemic HIV-1. Science 313, 523–526. 10.1126/science.1126531.

64. Sharp, P.M., and Hahn, B.H. (2011). Origins of HIV and the AIDS pandemic. Cold Spring Harb Perspect Med 1, a006841. 10.1101/cshperspect.a006841.

65. Korber, B., Muldoon, M., Theiler, J., Gao, F., Gupta, R., Lapedes, A., Hahn, B.H., Wolinsky, S., and Bhattacharya, T. (2000). Timing the ancestor of the HIV-1 pandemic strains. Science 288, 1789–1796.

66. Barbian, H.J., Decker, J.M., Bibollet-Ruche, F., Galimidi, R.P., West, A.P., Jr., Learn, G.H., Parrish, N.F., Iyer, S.S., Li, Y., Pace, C.S., et al. (2015). Neutralization properties of simian immunodeficiency viruses infecting chimpanzees and gorillas. mBio 6. 10.1128/mBio.00296-15.

67. Li, H., Wang, S., Lee, F.H., Roark, R.S., Murphy, A.I., Smith, J., Zhao, C., Rando, J., Chohan, N., Ding, Y., et al. (2021). New SHIVs and Improved Design Strategy for Modeling HIV-1 Transmission, Immunopathogenesis, Prevention and Cure. J Virol. 10.1128/JVI.00071-21.

68. van Schooten, J., van Haaren, M.M., Li, H., McCoy, L.E., Havenar-Daughton, C., Cottrell, C.A., Burger, J.A., van der Woude, P., Helgers, L.C., Tomris, I., et al. (2021). Antibody responses induced by SHIV infection are more focused than those induced by soluble native HIV-1 envelope trimers in non-human primates. PLoS Pathog 17, e1009736. 10.1371/journal.ppat.1009736.

69. Pegu, A., Xu, L., DeMouth, M.E., Fabozzi, G., March, K., Almasri, C.G., Cully, M.D., Wang, K., Yang, E.S., Dias, J., et al. (2022). Potent anti-viral activity of a trispecific HIV neutralizing antibody in SHIV-infected monkeys. Cell Rep 38, 110199. 10.1016/j.celrep.2021.110199.

70. Li, H., Wang, S., Lee, F.H., Roark, R.S., Murphy, A.I., Smith, J., Zhao, C., Rando, J., Chohan, N., Ding, Y., et al. (2021). New SHIVs and Improved Design Strategy for Modeling HIV-1 Transmission, Immunopathogenesis, Prevention and Cure. J Virol 95. 10.1128/JVI.00071-21.

71. Hirsch, V.M., Santra, S., Goldstein, S., Plishka, R., Buckler-White, A., Seth, A., Ourmanov, I., Brown, C.R., Engle, R., Montefiori, D., et al. (2004). Immune failure in the absence of profound CD4+ T-lymphocyte depletion in simian immunodeficiency virus-infected rapid progressor macaques. J Virol 78, 275–284. 10.1128/jvi.78.1.275-284.2004.

72. Salazar-Gonzalez, J.F., Bailes, E., Pham, K.T., Salazar, M.G., Guffey, M.B., Keele, B.F., Derdeyn, C.A., Farmer, P., Hunter, E., Allen, S., et al. (2008). Deciphering human immunodeficiency virus type 1 transmission and early envelope diversification by single-genome amplification and sequencing. J Virol 82, 3952–3970. 10.1128/JVI.02660-07.

73. Keele, B.F., Giorgi, E.E., Salazar-Gonzalez, J.F., Decker, J.M., Pham, K.T., Salazar, M.G., Sun, C., Grayson, T., Wang, S., Li, H., et al. (2008). Identification and characterization of transmitted and early founder virus envelopes in primary HIV-1 infection. Proceedings of the National Academy of Sciences of the United States of America 105, 7552–7557. 10.1073/pnas.0802203105.

74. Pancera, M., McLellan, J.S., Wu, X., Zhu, J., Changela, A., Schmidt, S.D., Yang, Y., Zhou, T., Phogat, S., Mascola, J.R., and Kwong, P.D. (2010). Crystal structure of PG16 and chimeric dissection with somatically related PG9: structure-function analysis of two quaternary-specific antibodies that effectively neutralize HIV-1. J Virol 84, 8098–8110. 10.1128/JVI.00966-10.

75. Vazquez Bernat, N., Corcoran, M., Nowak, I., Kaduk, M., Castro Dopico, X., Narang, S., Maisonasse, P., Dereuddre-Bosquet, N., Murrell, B., and Karlsson Hedestam, G.B. (2021). Rhesus and cynomolgus macaque immunoglobulin heavy-chain genotyping yields comprehensive databases of germline VDJ alleles. Immunity 54, 355–366 e354. 10.1016/j.immuni.2020.12.018.

76. Mirdita, M., Schutze, K., Moriwaki, Y., Heo, L., Ovchinnikov, S., and Steinegger, M. (2022). ColabFold: making protein folding accessible to all. Nat Methods 19, 679–682. 10.1038/s41592-022-01488-1.

77. Pettersen, E.F., Goddard, T.D., Huang, C.C., Meng, E.C., Couch, G.S., Croll, T.I., Morris, J.H., and Ferrin, T.E. (2021). UCSF ChimeraX: Structure visualization for researchers, educators, and developers. Protein Sci 30, 70–82. 10.1002/pro.3943.

78. Mirdita, M., Steinegger, M., and Soding, J. (2019). MMseqs2 desktop and local web server app for fast, interactive sequence searches. Bioinformatics 35, 2856–2858. 10.1093/bioinformatics/bty1057.

79. Victora, G.D., and Nussenzweig, M.C. (2022). Germinal Centers. Annu Rev Immunol 40, 413–442. 10.1146/annurev-immunol-120419-022408.

80. Abbott, R.K., and Crotty, S. (2020). Factors in B cell competition and immunodominance. Immunol Rev 296, 120–131. 10.1111/imr.12861.

81. Kuraoka, M., Schmidt, A.G., Nojima, T., Feng, F., Watanabe, A., Kitamura, D., Harrison, S.C., Kepler, T.B., and Kelsoe, G. (2016). Complex Antigens Drive Permissive Clonal Selection in Germinal Centers. Immunity 44, 542–552. 10.1016/j.immuni.2016.02.010.

82. Berek, C., Berger, A., and Apel, M. (1991). Maturation of the immune response in germinal centers. Cell 67, 1121–1129.

83. Manser, T., Huang, S.Y., and Gefter, M.L. (1984). Influence of clonal selection on the expression of immunoglobulin variable region genes. Science 226, 1283–1288.

84. Jacob, J., Kelsoe, G., Rajewsky, K., and Weiss, U. (1991). Intraclonal generation of antibody mutants in germinal centres. Nature 354, 389–392. 10.1038/354389a0.

85. Tas, J.M., Mesin, L., Pasqual, G., Targ, S., Jacobsen, J.T., Mano, Y.M., Chen, C.S., Weill, J.C., Reynaud, C.A., Browne, E.P., et al. (2016). Visualizing antibody affinity maturation in germinal centers. Science 351, 1048–1054. 10.1126/science.aad3439.

86. Bonsignori, M., Zhou, T., Sheng, Z., Chen, L., Gao, F., Joyce, M.G., Ozorowski, G., Chuang, G.Y., Schramm, C.A., Wiehe, K., et al. (2016). Maturation Pathway from Germline to Broad HIV-1 Neutralizer of a CD4-Mimic Antibody. Cell 165, 449–463. 10.1016/j.cell.2016.02.022.

87. Garces, F., Sok, D., Kong, L., McBride, R., Kim, H.J., Saye-Francisco, K.F., Julien, J.P., Hua, Y., Cupo, A., Moore, J.P., et al. (2014). Structural evolution of glycan recognition by a family of potent HIV antibodies. Cell 159, 69–79. 10.1016/j.cell.2014.09.009.

88. Pauthner, M.G., Nkolola, J.P., Havenar-Daughton, C., Murrell, B., Reiss, S.M., Bastidas, R., Prevost, J., Nedellec, R., von Bredow, B., Abbink, P., et al. (2019). Vaccine-Induced Protection from Homologous Tier 2 SHIV Challenge in Nonhuman Primates Depends on Serum-Neutralizing Antibody Titers. Immunity 50, 241–252 e246. 10.1016/j.immuni.2018.11.011.

89. Andrabi, R., Pallesen, J., Allen, J.D., Song, G., Zhang, J., de Val, N., Gegg, G., Porter, K., Su, C.Y., Pauthner, M., et al. (2019). The Chimpanzee SIV Envelope Trimer: Structure and Deployment as an HIV Vaccine Template. Cell reports 27, 2426–2441.e2426. 10.1016/j.celrep.2019.04.082.

90. Li, H., Wang, S., Kong, R., Ding, W., Lee, F.H., Parker, Z., Kim, E., Learn, G.H., Hahn, P., Policicchio, B., et al. (2016). Envelope residue 375 substitutions in simian-human immunodeficiency viruses enhance CD4 binding and replication in rhesus macaques. Proc Natl Acad Sci U S A 113, E3413–3422. 10.1073/pnas.1606636113.

91. Zhao, F., Berndsen, Z.T., Pedreno-Lopez, N., Burns, A., Allen, J.D., Barman, S., Lee, W.H., Chakraborty, S., Gnanakaran, S., Sewall, L.M., et al. (2022). Molecular insights into antibody-mediated protection against the prototypic simian immunodeficiency virus. Nat Commun 13, 5236. 10.1038/s41467-022-32783-2.

92. Bangaru, S., Antanasijevic, A., Kose, N., Sewall, L.M., Jackson, A.M., Suryadevara, N., Zhan, X., Torres, J.L., Copps, J., de la Pena, A.T., et al. (2022). Structural mapping of antibody landscapes to human betacoronavirus spike proteins. Sci Adv 8, eabn2911. 10.1126/sciadv.abn2911.

93. Shlemov, A., Bankevich, S., Bzikadze, A., Turchaninova, M.A., Safonova, Y., and Pevzner, P.A. (2017). Reconstructing Antibody Repertoires from Error-Prone Immunosequencing Reads. J Immunol 199, 3369–3380. 10.4049/jimmunol.1700485.

94. Monigatti, F., Gasteiger, E., Bairoch, A., and Jung, E. (2002). The Sulfinator: predicting tyrosine sulfation sites in protein sequences. Bioinformatics 18, 769–770. 10.1093/bioinformatics/18.5.769.

95. Sievers, F., Wilm, A., Dineen, D., Gibson, T.J., Karplus, K., Li, W., Lopez, R., McWilliam, H., Remmert, M., Soding, J., et al. (2011). Fast, scalable generation of high-quality protein multiple sequence alignments using Clustal Omega. Mol Syst Biol 7, 539. 10.1038/msb.2011.75.

96. Moore, R.M., Harrison, A.O., McAllister, S.M., Polson, S.W., and Wommack, K.E. (2020). Iroki: automatic customization and visualization of phylogenetic trees. Peerj 8, e8584. 10.7717/peerj.8584.

97. Hraber, P., Korber, B., Wagh, K., Giorgi, E.E., Bhattacharya, T., Gnanakaran, S., Lapedes, A.S., Learn, G.H., Kreider, E.F., Li, Y., et al. (2015). Longitudinal Antigenic Sequences and Sites from Intra-Host Evolution (LASSIE) Identifies Immune-Selected HIV Variants. Viruses 7, 5443–5475. 10.3390/v7102881.

98. Tareen, A., and Kinney, J.B. (2020). Logomaker: beautiful sequence logos in Python. Bioinformatics 36, 2272–2274 10.1093/bioinformatics/btz921.

99. Ranwez, V., Douzery, E.J.P., Cambon, C., Chantret, N., and Delsuc, F. (2018). MACSE v2: Toolkit for the Alignment of Coding Sequences Accounting for Frameshifts and Stop Codons. Mol Biol Evol 35, 2582–2584 10.1093/molbev/msy159.

100. Altschul, S.F., Gish, W., Miller, W., Myers, E.W., and Lipman, D.J. (1990). Basic local alignment search tool. Journal of molecular biology 215, 403–410. 10.1016/S0022-2836(05)80360-2.

